# Age-associated failure of efficient cardiac proteostatic adaptations to chronic cAMP-stress is associated with accelerated heart aging

**DOI:** 10.1101/2023.08.13.553128

**Authors:** Maria Grazia Perino, Miguel Calvo-Rubio Barrera, Daniel R. Riordon, Giulio Agnetti, Admira Parveen, Christopher H. Morrell, Ismayil Ahmet, Khalid Chakir, Yelena S. Tarasova, Jia-Hua Qu, Kirill V Tarasov, Alexey E. Lyashkov, Yevgeniya O. Lukyanenko, Hikmet Kadioglu, Mark Ranek, Rafael deCabo, Edward G. Lakatta

## Abstract

Dysregulated proteostasis, leading to accumulation of misfolded proteins, electron-dense aggregates (lipofuscin, LF), preamyloid oligomers (PAOs), and proteotoxic stress is a hallmark of aging. We investigated how efficiently proteostatic adaptations to chronic cardiac cyclic adenosine monophosphate (cAMP)-dependent stress change with aging in mice harboring marked, cardiac-specific over-expression of adenylyl cyclase VIII (TG^AC8^). We assessed protein quality control (PQC) mechanisms: ubiquitin proteasome system (UPS), autophagic flux via macroautophagy, and mitophagy in left ventricles (LVs) of TG^AC8^ and wild type littermates (WT) at 3-4 months and at 17-21 months of age. At 3-4 months of age TG^AC8^ mice exhibited markers of increased autophagic flux, measured by levels of microtubule-associated protein 1 light chain 3 (LC3), p62, and their phospho-forms in TG^AC8^ LV; cathepsin L1 activity was also significantly increased. In addition, canonical mitophagy signaling was enhanced, as receptors PARKIN, p62^S405^ and p62^S351^ were all upregulated, confirming a more efficient proteostasis in TG^AC8^ at 3-4 months vs WT. In advanced age, however, the PQC mechanisms were overwhelmed by proteotoxic stress, manifested in insufficient proteasome activity and an unbalanced autophagic flux (accelerated for markers such as LC3A in the context of a slower *overall* flux), leading to an increase in the accumulation of protein aggregates (increased ratio of insoluble/soluble protein fractions). Although both canonical (PARKIN, p62^S405^ and p62^S351^ receptors) and non-canonical (FKBP8 receptor) mitophagy signaling were upregulated in advanced age in TG^AC8^, mitophagy was markedly impaired and mitochondrial dysfunction increased. Accumulation of LF bodies, of brownish-to-black pigments, and of LC3^+^ and p62^+^-inclusions of aberrant sizes, of desmin cardiac preamyloid oligomers (PAOs) and of cleaved desmin, tagged for ubiquitination, were all increased in TG^AC8^ compared to young TG^AC8^. In contrast, the rate of protein synthesis and levels of soluble aggregates were reduced in aged vs young TG^AC8^, a sign of “normal” aging. Thus, increased proteostatic mechanisms maintain cardiac health in TG^AC8^ in youth (3-4 months), but long-term exposure to chronic cardiac stress, imposed by sustained activation of the AC/cAMP/PKA/Ca^2+^ signaling axis, results in severely dysregulated proteostasis in TG^AC8^ vs WT mice, associated with proteostatic insufficiency and increased cardiomyopathy that leads to accelerated cardiac aging.

**GRAPHICAL ABSTRACT:** 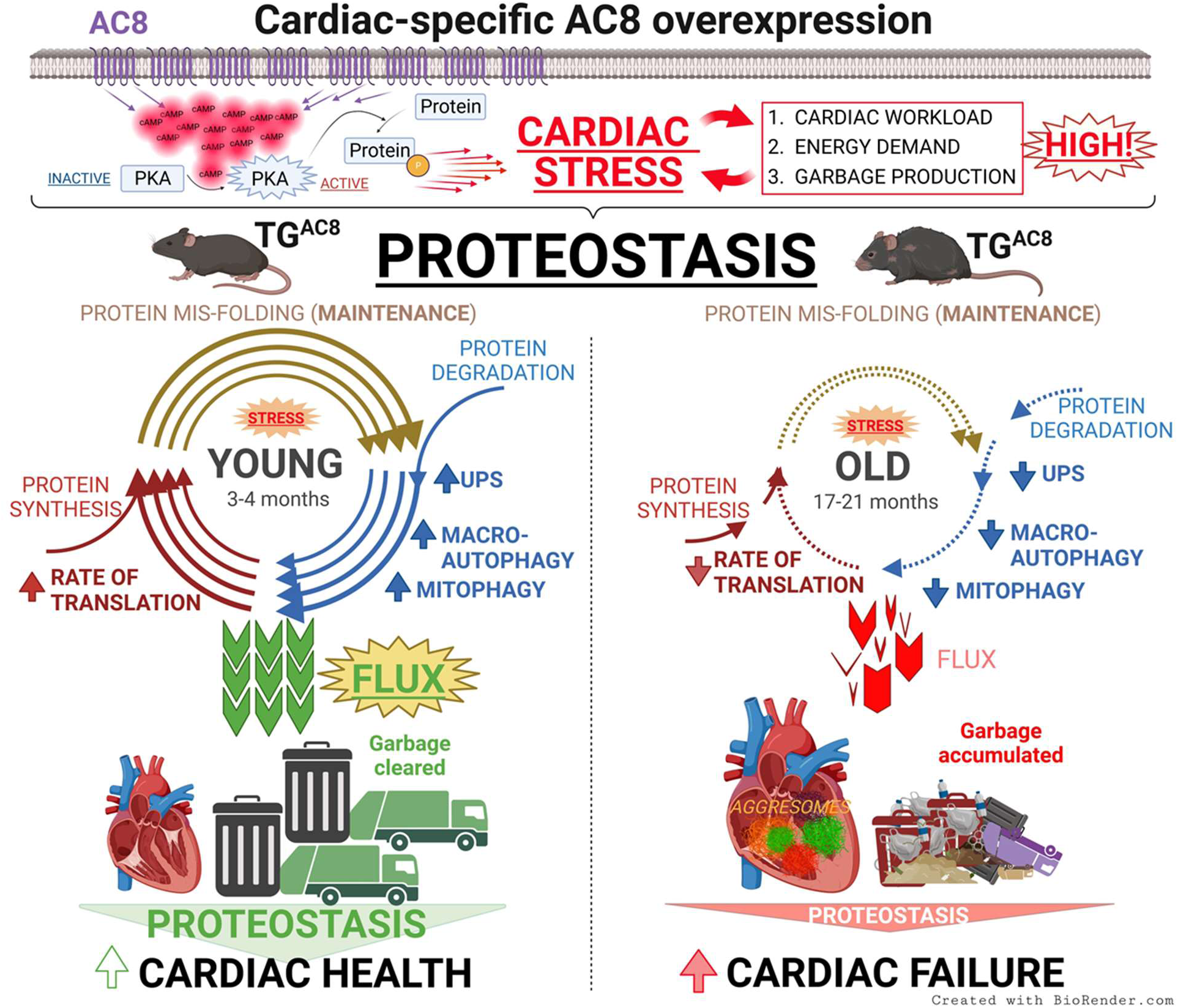

## INTRODUCTION

Cardiac protein quality control (PQC) is required for the maintenance of cardiac health and function. Pathogenic aspects of protein misfolding and aggregation, collectively referred to *proteotoxicity*, accompany advancing age and can ultimately lead to heart dysfunction^1^, and eventual cardiac failure^2^. Many protein aggregate–based diseases in the heart have been well characterized^3^. Plaque deposition (amyloidosis) or accumulation of soluble, oligomeric aggregates, and, more recently, desmin disorganization and the increase in cardiac desmin preamyloid oligomers (PAOs)^4^ have also emerged as common hallmarks of aging, and acquired heart failure (HF)^5,6^.

Many factors, including an imbalance of protein synthesis and degradation, exhaustion of proteostatic machinery, and downregulation of clearing mechanisms, e.g. the ubiquitin proteasome system (UPS)^7^ and autophagy^8^, can lead to collapse of proteostasis. Impaired autophagy and mitophagy that accompany advancing age due to reduced lysosomal^9^ and mitochondrial fitness resulting from increased oxidative stress^10^, have been linked to the accumulation of electron-dense aggregates, generally referred to lipofuscin (LF). As age advances, accumulation of LF occurs as PQC mechanisms become insufficient and begin to collapse^11^. The linear increase of LF as age advances is inversely correlated with longevity^12,13^.

Our recent study demonstrated that chronic severe cardiac stress imposed by over-expression of adenylyl cyclase VIII (AC8) in mice (TG^AC^^8^), induced concentric pattern of adaptations-signaling circuitry at a young age (3-4 months)^14^. This adaptive signaling circuitry included impressive upregulation of protein synthesis and degradation mechanisms (proteasome and autophagy) that protect against cardiac stress, maintaining the remarkably high performance of the heart for up to about one year of age^15,16^. However, signs of heart failure (HF) and cardiomyopathy begin to emerge in TG^AC^^8^ as age advances^17,18^. Here we tested the hypothesis that the adaptive proteostatic mechanisms in response to severe sustained cAMP-dependent chronic cardiac stress become *more impaired* (overwhelmed) in TG^AC8^ than in WT in advanced age.

## RESULTS

### Autophagy is <u>upregulated</u> in the TG^AC8^ heart at 3-4 months of age

To assess basal autophagy in LVs in young TG^AC8^ and WT, we performed WB of the established autophagy markers LC3 and p62/sequestosome-1, and assessed their activity via measuring their phosphorylation states. Both LC3A and LC3B isoforms, which localize on different autophagosomes^19^ and exhibit distinct expression patterns and functions^20^, were assessed using isoform-specific antibodies.

Significant changes in cytosolic LC3I and its lipidated product, LC3II (LC3-PE), recruited to the autophagosomal membrane during autophagy, were evident in TG^AC8^ vs WT. Specifically at 3-4 months of age: LC3AI was significantly upregulated in TG^AC8^ vs WT, whereas LC3AII was unchanged (**Figure 1Ai**); in contrast, cytosolic LC3BI did not change in TG^AC8^, whereas lipidated LC3BII was significantly downregulated (**Figure 1Aii**), compared to WT. Further, phosphorylation of LC3 at S12, which inhibits LC3 activity^21^, was reduced by 50% (**Figure 1Bi**), and phosphorylation of LC3 at T50, which increases LC3 activity^22^, was increased by 30% (**Figure 1Bii**) in TG^AC8^ vs WT, suggesting a <u>more efficient</u> autophagic context in TG^AC8^ vs WT.

**Figure 1.**
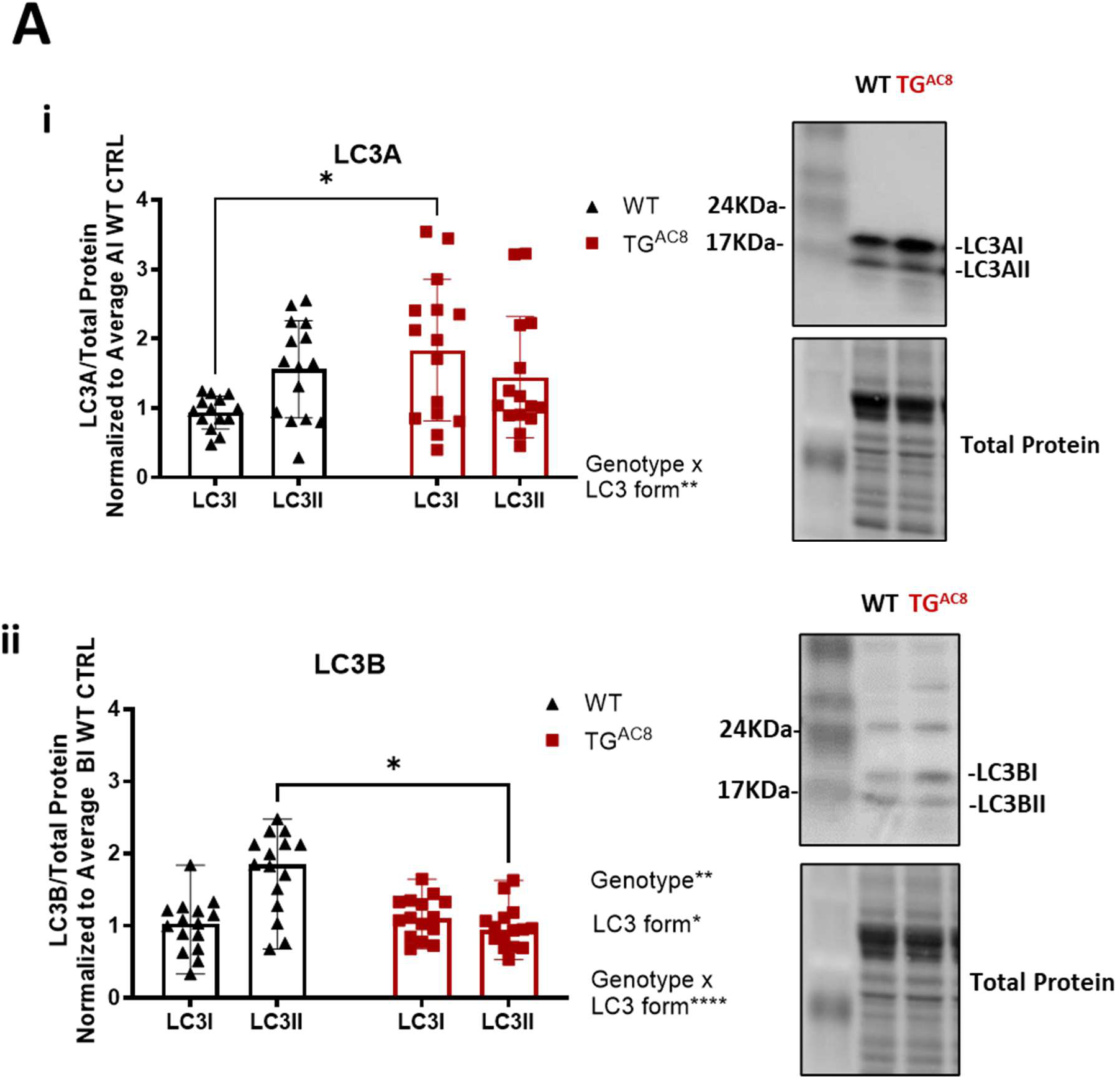
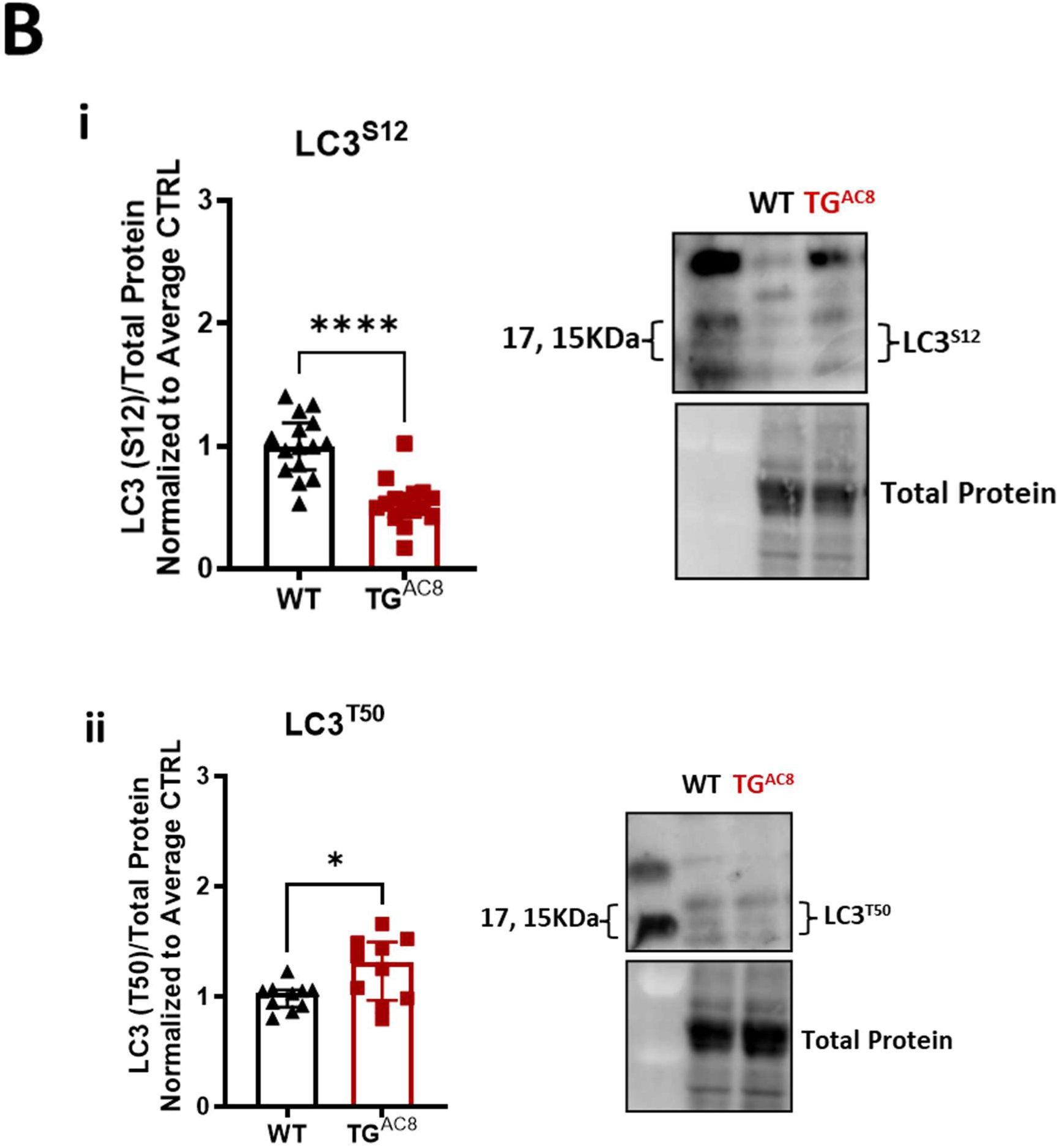
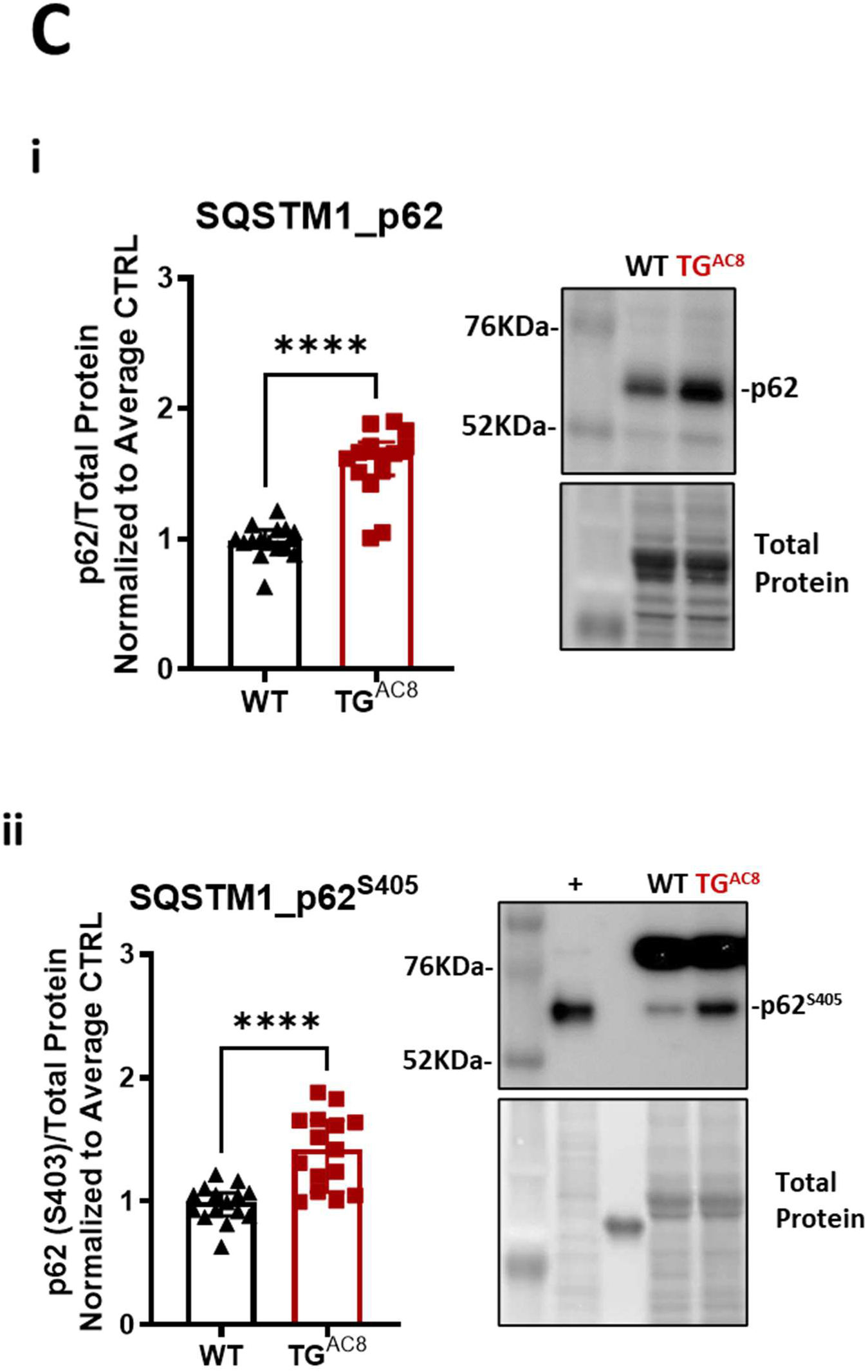
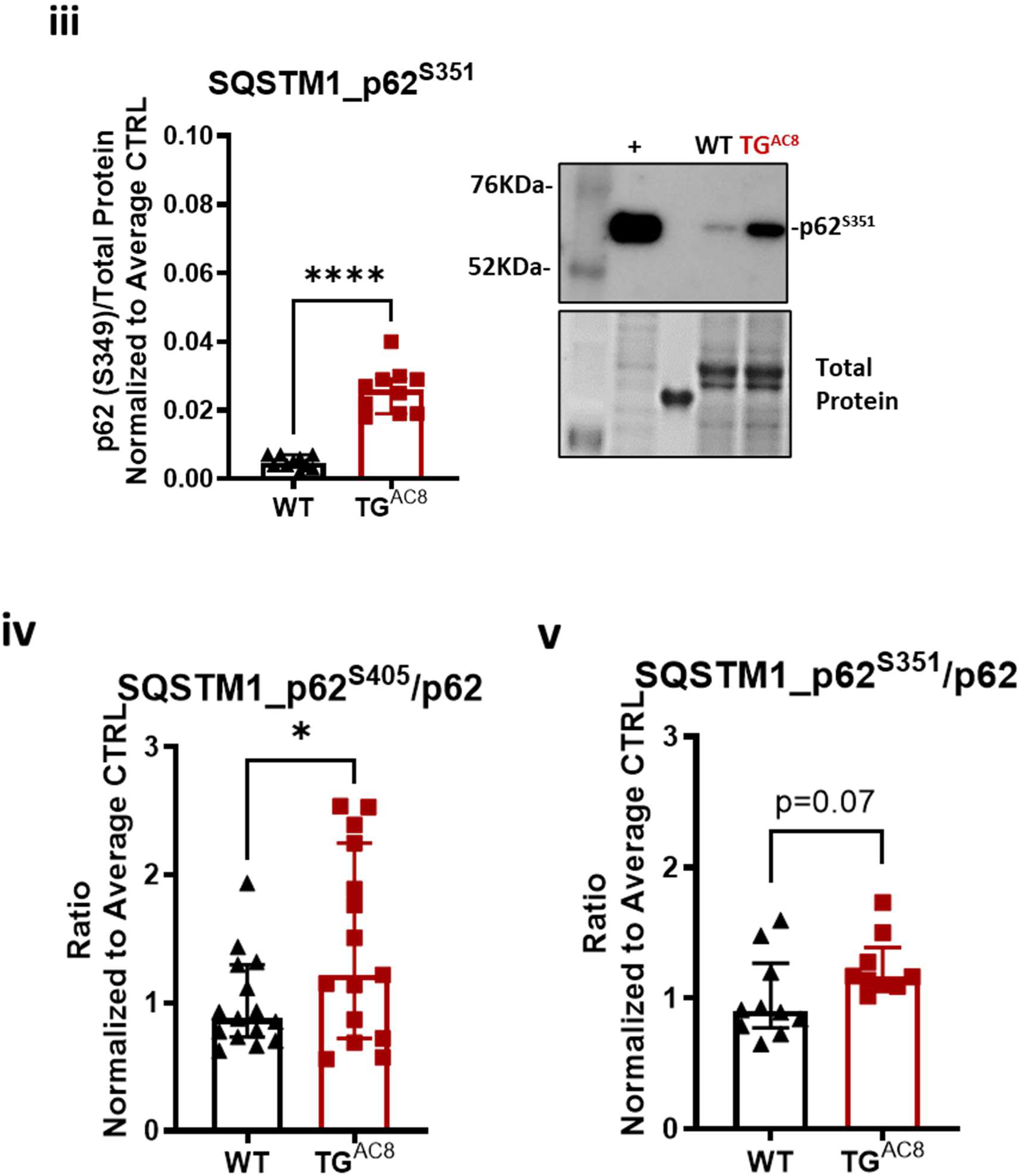
Selective and non-selective autophagy are both upregulated in TG^AC8^ at 3-4 months of age. Bar graphs and Western blot analysis of autophagy markers LC3 and its phosphorylation in young TG^AC8^ and wild type littermates (WT) at 3-4 months of age (n=15 mice per group). Data are presented as the median ±interquartile range. **Ai**), LC3A; **Aii**), LC3B; **Bi**), LC3^S12^; **Bii**), LC3^T50^. **Ci**), p62; **Cii**), p62^S405^; **C**), p62^S351^; **D**), p62^S405^/p62 ratio; **E**), p62^S351^/p62 ratio. **Ai-ii**: A 3-way ANOVA with mixed-effect model for repeated measurements, followed by Original FDR method of Benjamini and Hochberg post-hoc multi-comparison test was used. **B and C**: Unpaired 2-tailed Student’s tests with Welch’s correction was used. Symbol **\*** indicates significant (p<0.05) differences between genotypes (WT vs TG^AC8^).

Expression of total p62, an essential autophagy protein adaptor^23^, was significantly upregulated in TG^AC8^ vs WT (**Figure 1Ci**), and p62 was hyper-phosphorylated at S405 (p62^S405^) (**Figure 1Cii**) and at S351 (p62^S351^) (**Figure 1Ciii**), compared to WT; the ratios of phospho-proteins at both sites to total p62 were also significantly elevated (**Figures 1Civ-v**), suggesting a higher detection of “aggregates”^24^ (because phosphorylation at S405 increases affinity to ubiquitinated cargo^25^, including depolarized mitochondria^26^), and more efficient oxidative stress management (phosphorylation at S351 activates NRF2-pathway^27^) in the TG^AC8^ vs WT hearts, at 3-4 months of age.

Our previous work had shown that protein levels of lysosomal cathepsins are upregulated in the TG^AC8^ at 3-4 months of age^14^. Here, we quantified relative changes in different cathepsins in the TG^AC8^ LV, compared to WT. Cathepsins Z and S were *more enriched* in TG^AC8^, whereas cathepsin L1 showed the least significant change (**Figure 2A**, **Extended Data Table**). Because cathepsin L degrades not only lysosomal content but also autophagosome membrane-components GABARAP-II and LC3-II^28^, during the autophagic proteolysis, we assessed its activity as a measure of lysosome function and dispose of autophagy flux. A significantly higher cathepsin L1 activity was in TG^AC8^ vs WT LV (**Figure 2B**) in the context of an increase in the lysosomal acid phosphatase ACP2 protein **(Figure 2C**), suggesting an <u>overall higher lysosomal fitness</u> (function) in TG^AC8^ compared to WT.

**Figure 2.**
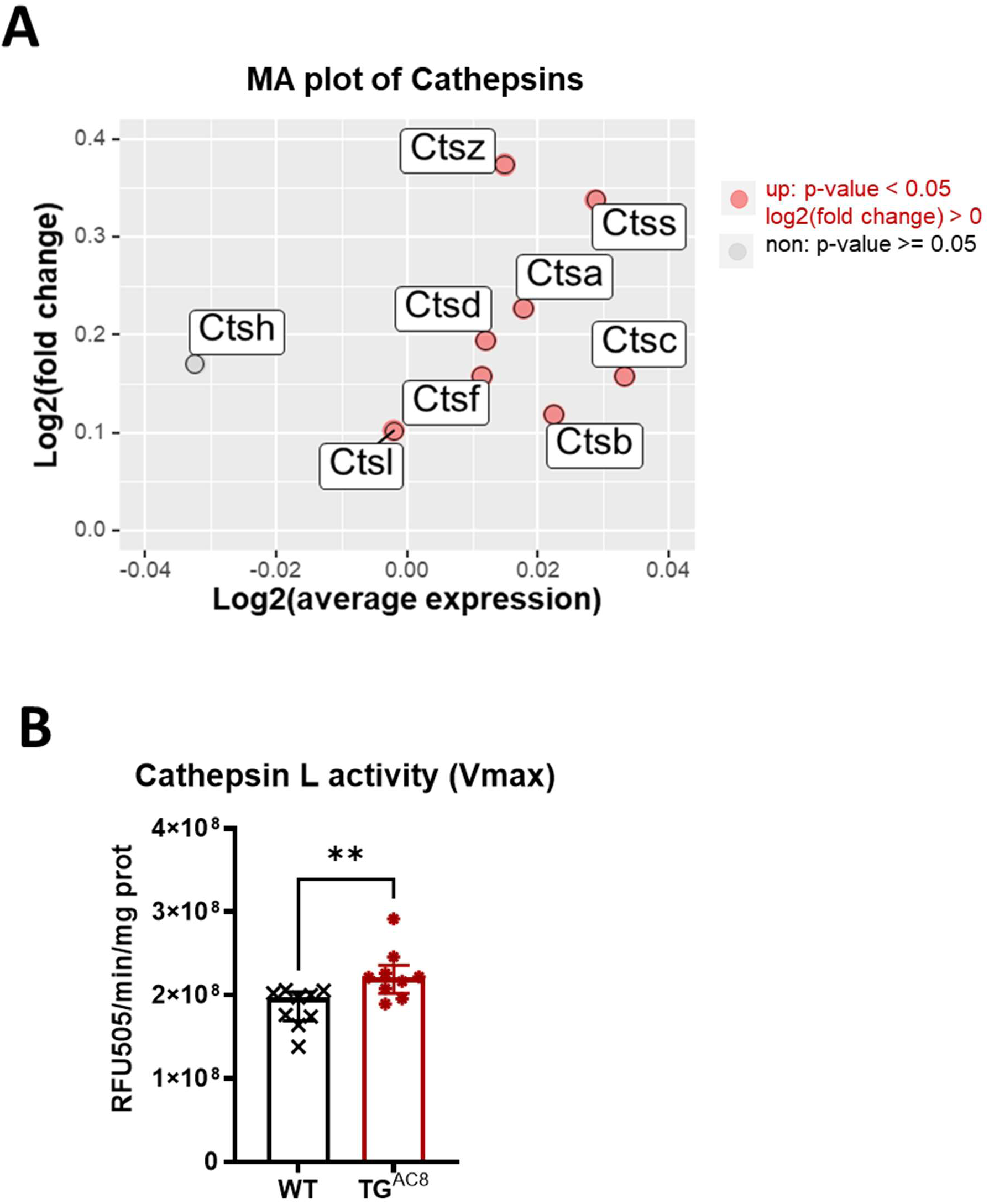
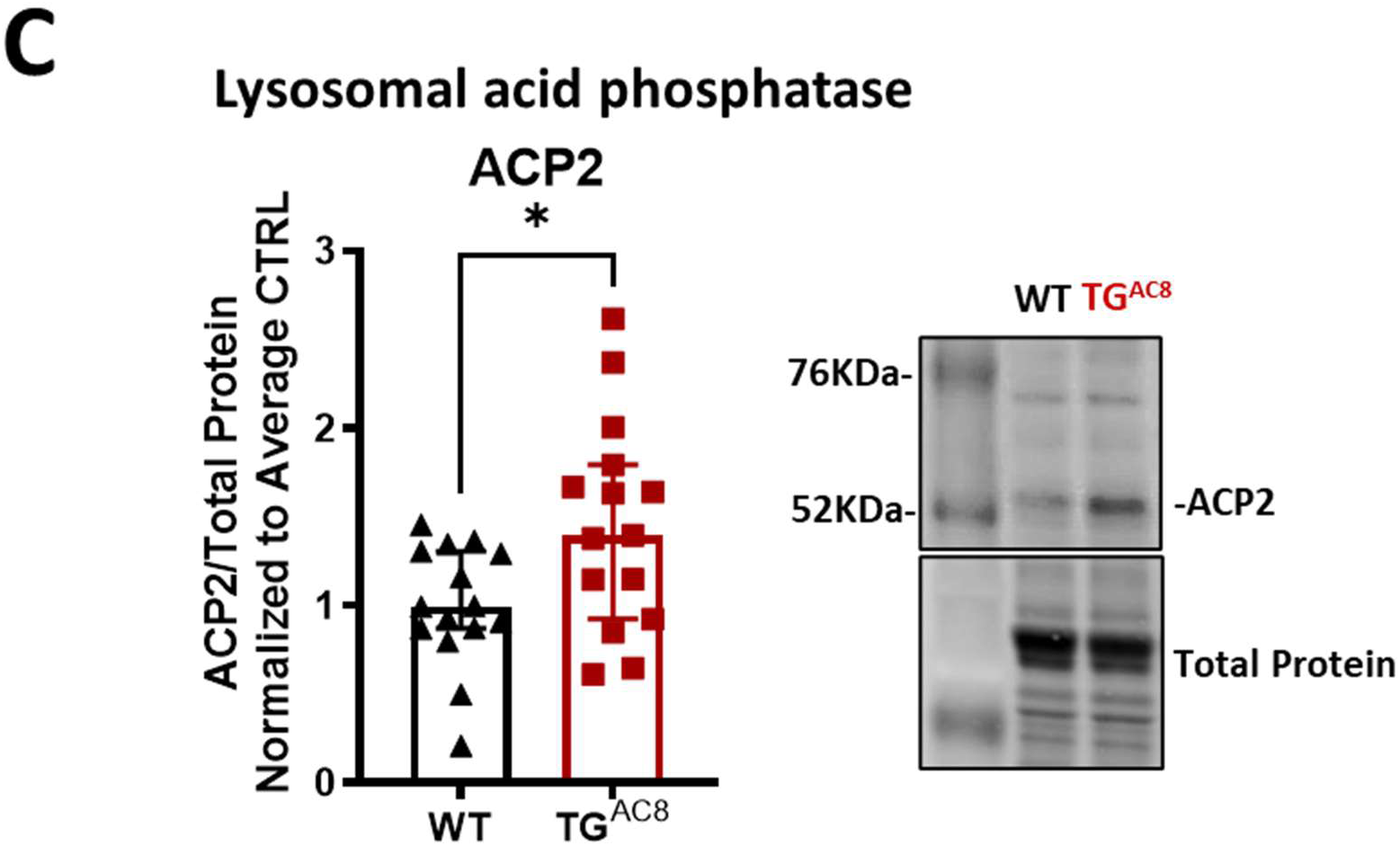
Lysosomal hydrolases are upregulated in TG^AC8^ at 3-4 months of age. **A**), MA plot showing enrichment in the relative amount of lysosomal cathepsin obtained from OMICs data in TG^AC8^ at 3-4 months of age. **B**), Cathepsin L1 activity measured in TG^AC8^ and WT littermates at 3-4 months of age (n=6 mice per group). Activity was measured at Vmax, and it is plotted as emitted fluorescence per mL, per mg of protein. **C**), Bar graphs and Western blot analysis of the lysosomal acid phosphatase, ACP2 (n=15 mice per group). Unpaired 2-tailed Student t test was used in **B** and unpaired 2-tailed Student t test with Welch’s correction was used in **C**. Data are presented as the median ± interquartile range. Symbol **\*** indicates significant (p<0.05) differences between genotypes (WT vs TG^AC8^).

### Autophagic flux is <u>optimized</u> to ensure efficient and effective clearance in the TG^AC8^ heart at 3-4 months of age

Because autophagy is a dynamic, multi-step process, we next evaluated stages of autophagic flux by inhibiting lysosomal function with the lysosomotropic drug chloroquine (CQ). Basal autophagy clearly differed in the young TG^AC8^ vs WT (**Figures 3Ai-iii, Extended Data Figure S1**): autophagic “carrier flux” (flux of the “carrier” protein not the actual cargo/substrate flux) of both <u>LC3AI</u> and <u>LC3BI</u> (**Figures 3Ai-ii**) became significantly reduced following CQ treatment, compared to saline, whereas for <u>LC3AII</u> and <u>LC3BII</u> remained unchanged; carrier-flux of the ubiquitin-binding protein^25^ and NRF2-signaling activator^29,30^ <u>p62</u> (**Figure 3Aiii**), was also downregulated (by 29%) in TG^AC8^ vs WT, whereas the pattern of the carrier-flux of other important players involved in autophagy following CQ was more variable. Specifically, the carrier-flux of the *autophagy related 16 like 1* protein (<u>ATG16L1</u>), which participates in LC3 lipidation^31^ and cell homeostasis by regulating membrane cellular trafficking^32^, was reduced (by 37%) (**Figure 3Bi**). In contrast, the carrier-flux of the *autophagy related 4B cysteine peptidase* (<u>ATG4B</u>) (**Figure 3Bii**), which is involved in LC3 lifecycle^33^ by processing pro-LC3 into LC3**I**^33^, and of the programmed cell death 6-interacting protein (<u>PDCD6IP/ALIX)</u>, ancillary protein contributing to the ESCRT signaling^34^, were both increased (by 42% and 28%, respectively) (**Figure 3Biii**). Finally, carrier-flux of the *peptidyl-prolyl cis-trans isomerase* (<u>FKBP8)</u> (**Figure 3Biv**), non-canonical mitophagy receptor^35^ that after recruitment on mitochondria from LC3A translocates to the ER (avoiding to be degraded by autophagosomes^36^), did not change between genotype.

**Figure 3.**
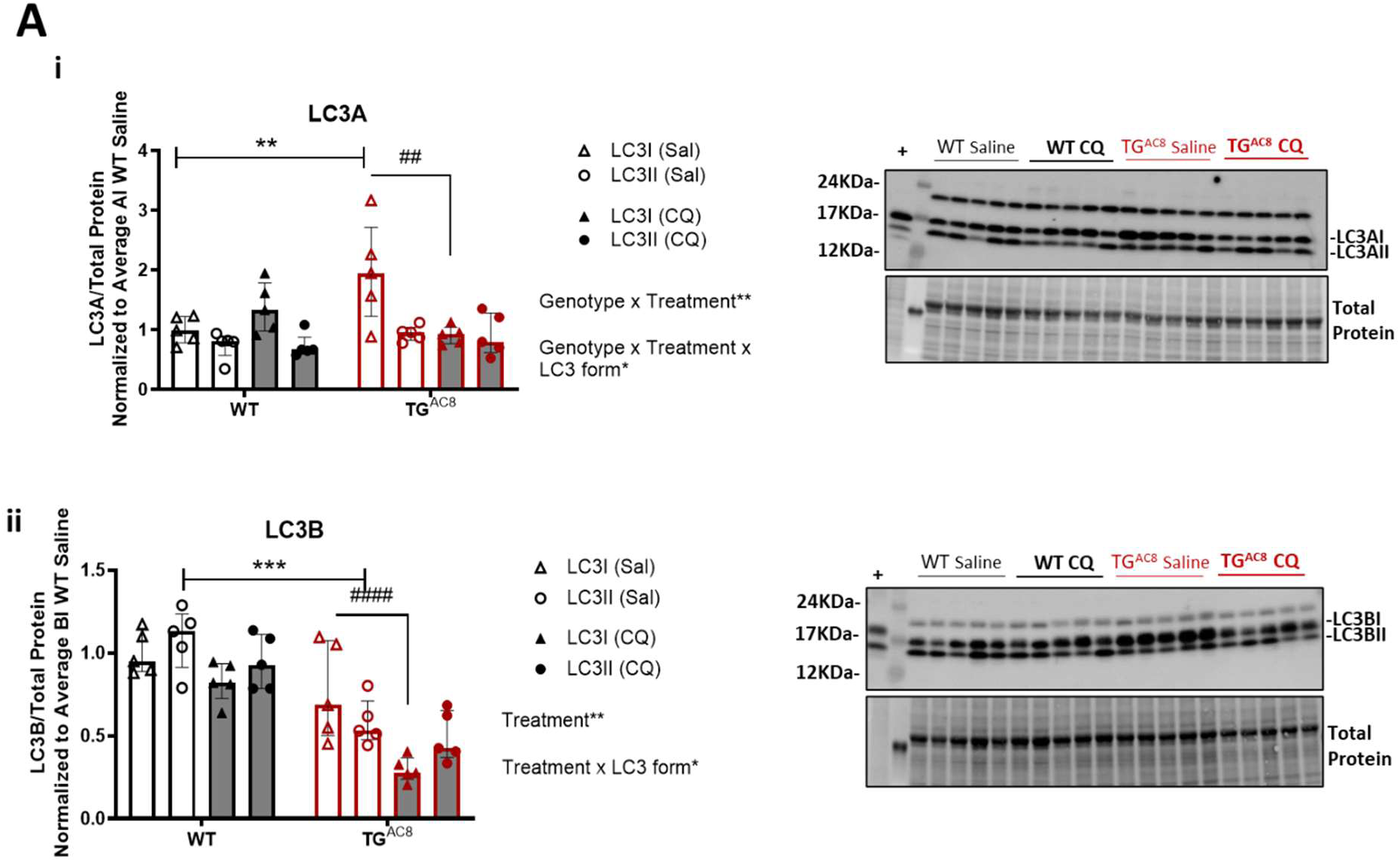
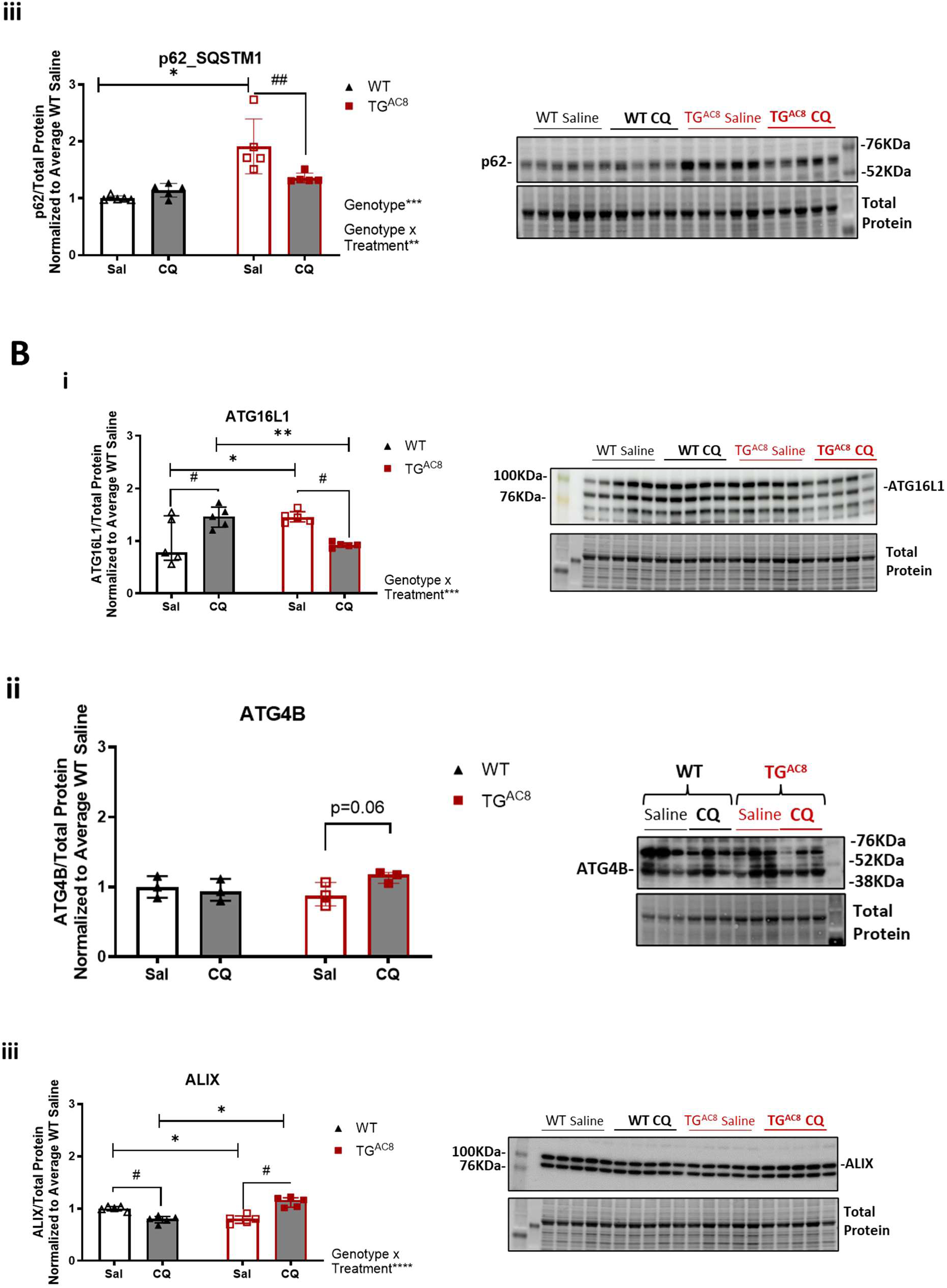
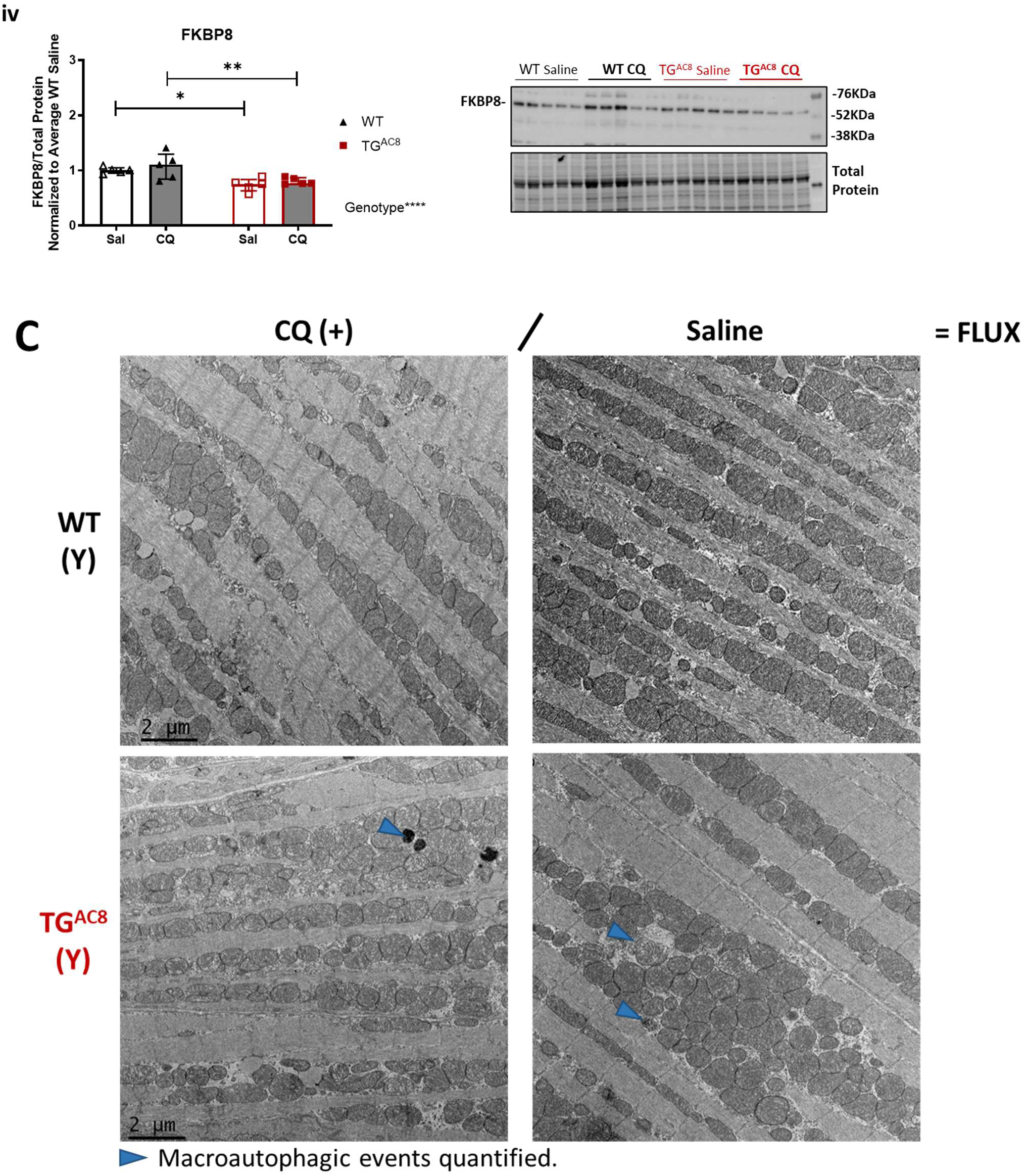
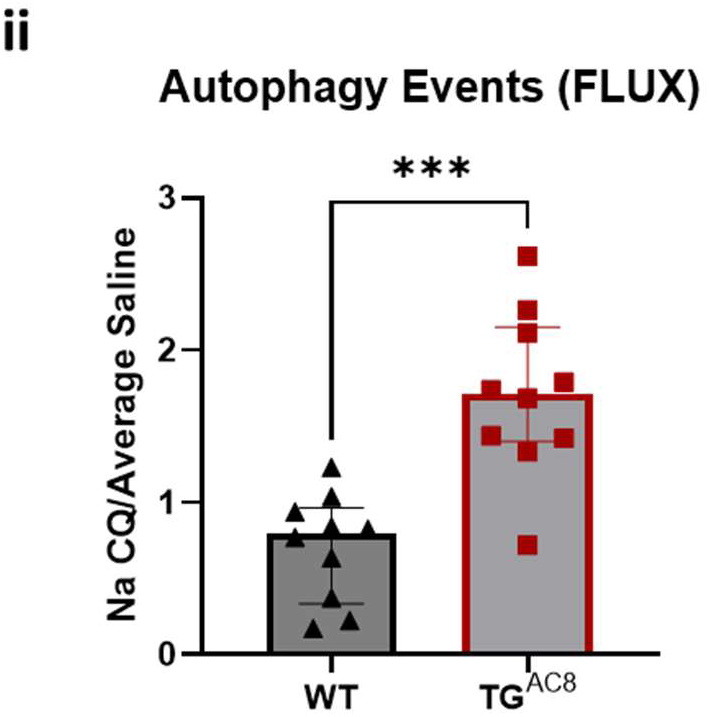
Autophagic flux is enhanced in TG^AC8^ at 3-4 months of age. TG^AC8^ and WT at 3-4 months of age were treated by intraperitoneal administration (IP) of chloroquine (CQ) (50 mg/kg) or saline, and LVs were collected 3 hours after drug administration and snap-frozen for analysis of autophagy markers LC3, p62, ATG16L1, ATG4B, ALIX and FKBP8 (n=5 mice per group). Bar graphs and Western blot analysis of **Ai**), LC3A; **Aii**), LC3B; **Aiii**), p62; **Bi**), ATG16L1; **Bii**), ATG4B; **Biii**), ALIX; **Biv**), FKBP8; **Ci**), TEM representative micrographs and **Cii**) Paired quantification of macroautophagic figures, normalized to average WT saline. **Ai-ii**: A three-way ANOVA with repeated measurements, followed by Original FDR method of Benjamini and Hochberg post-hoc multi-comparison test was used. **Aiii and B**: A two-way ANOVA, followed by Original FDR method of Benjamini and Hochberg post-hoc multi-comparison test was used. **C**: Unpaired 2-tailed Student t test with Welch’s correction was used. Data are presented as the median ±interquartile range. Symbols **\* ^#^** indicate significant (p<0.05) differences: between genotypes* (WT vs TG^AC8^), between treatments^#^ (CQ vs saline). Ai), LC3A; Aii), LC3B; Aiii), p62; Bi), ATG16L1; Bii), ATG4B; Biii), ALIX; Biv), FKBP8; Ci), TEM representative micrographs and Cii) Paired quantification of macroautophagic figures, normalized to average WT saline. Ai-ii: A three-way ANOVA with repeated measurements, followed by Original FDR method of Benjamini and Hochberg post-hoc multi-comparison test was used. Aiii and B: A two-way ANOVA, followed by Original FDR method of Benjamini and Hochberg post-hoc multi-comparison test was used. C: Unpaired 2-tailed Student t test with Welch’s correction was used. Data are presented as the median ±interquartile range. Symbols * # indicate significant (p<0.05) differences: between genotypes* (WT vs TGAC8), between treatments# (CQ vs saline).

Using transmission electron microscopy (TEM), we next assessed events-flux by directly visualizing autophagic figures in saline vs CQ-treated TG^AC8^ and age-matched WT controls. A significant increase in the numbers of autophagic events was detected in TG^AC8^ vs WT (**Figures 3Ci-ii**), confirming the <u>increase in autophagy/autophagic flux</u> in the TG^AC8^ LV at this age.

In summary, chronic cardiac-specific over-expression of AC8 in the young TG^AC8^ heart^14^ modulates protein levels of the autophagic machinery by fine-tuning key markers of selective cargo recognition (LC3 isoforms), and of selective autophagy (LC3/p62 activation by phosphorylation), activates protection mechanisms against oxidative stress, and enhances lysosomal degradation ensuring a more efficient and effective clearance of misfolded proteins without accumulation of insoluble protein aggregates.

### <u>Increased efficiency</u> of autophagy/autophagic flux of the youthful TG^AC8^ heart <u>becomes insufficient</u> in advanced age

Because autophagy induction and clearance are known to decrease with aging^9^, we anticipated that expression of key autophagic proteins and dissipated autophagic flux became reduced in LV of both aged TG^AC8^ and matched WT. Indeed, both LC3 isoforms (**Figures 4Ai-ii**) and ATG4B (**Extended Data Figure S2**) were significantly downregulated in both genotypes at 17-21 vs 3-4 months of age, with specific accumulation of LC3II, an age-associated decrease in event number/volumetric density at *early* stages (**Figures 4Bi-ii, 4Bv**), and an increase in event number/volumetric density in *late* stages, measured by paired TEM micrograph analysis (planimetry and stereology) (**Figures 4Biii-v**).

**Figure 4.**
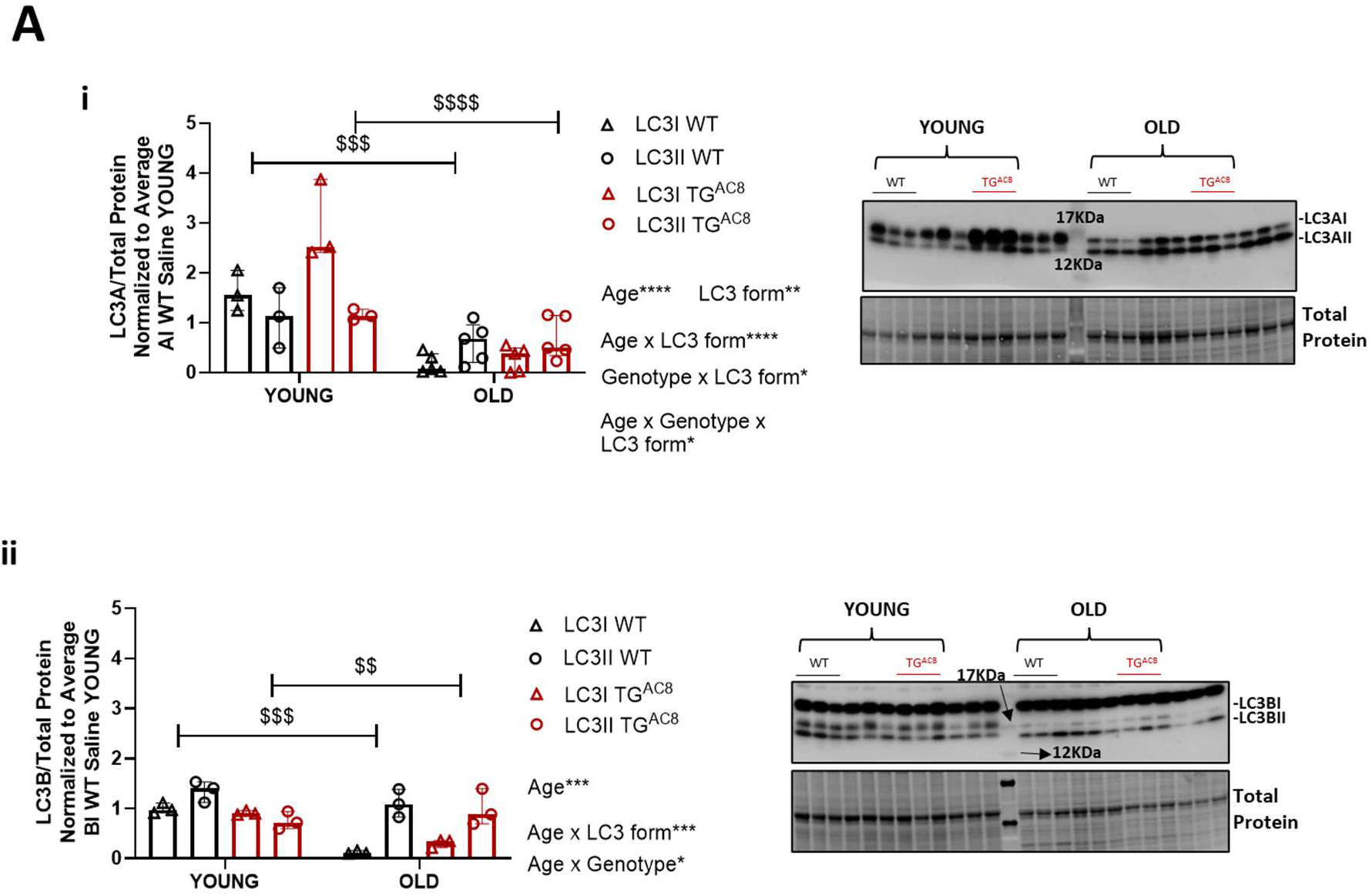
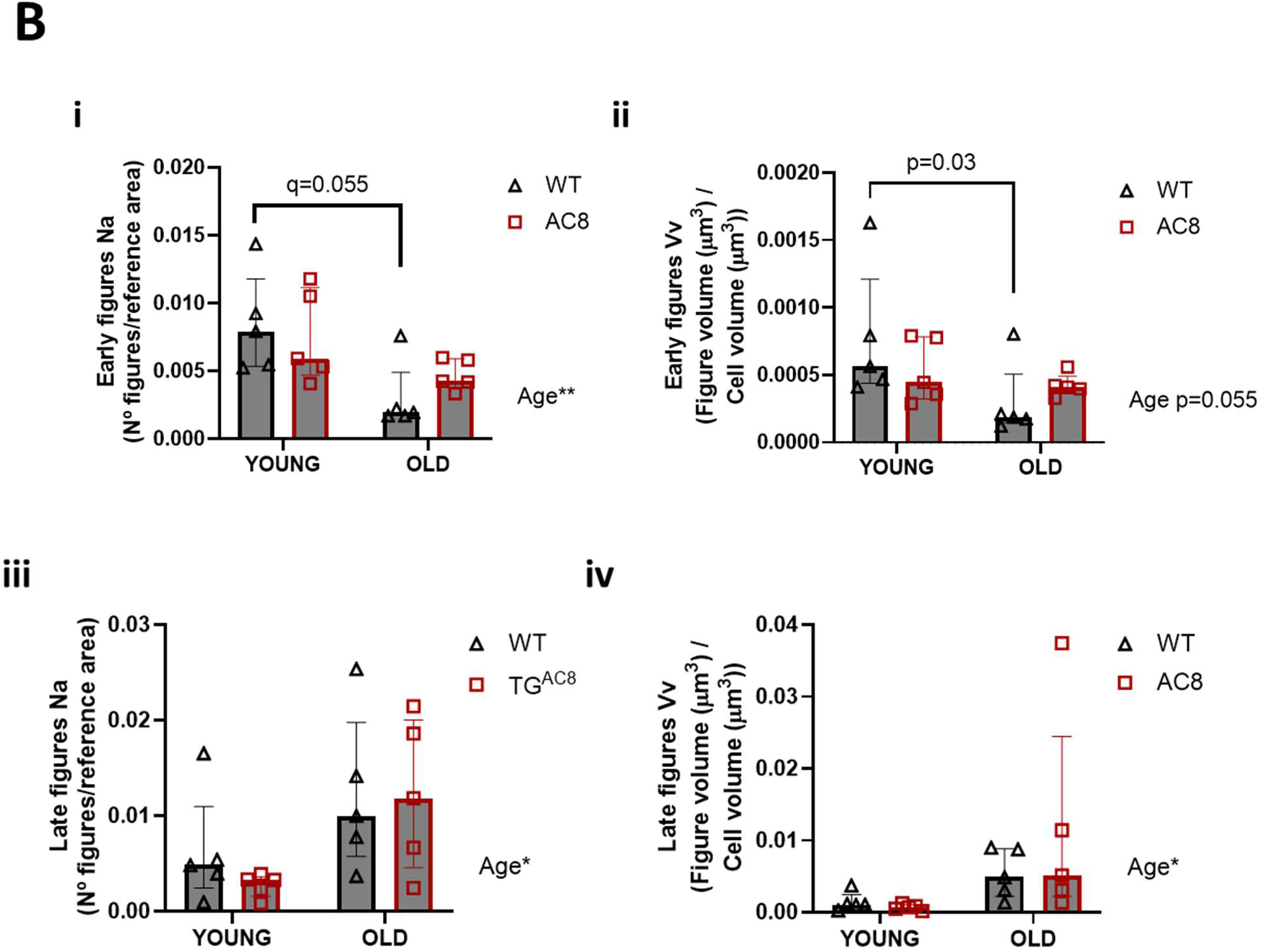
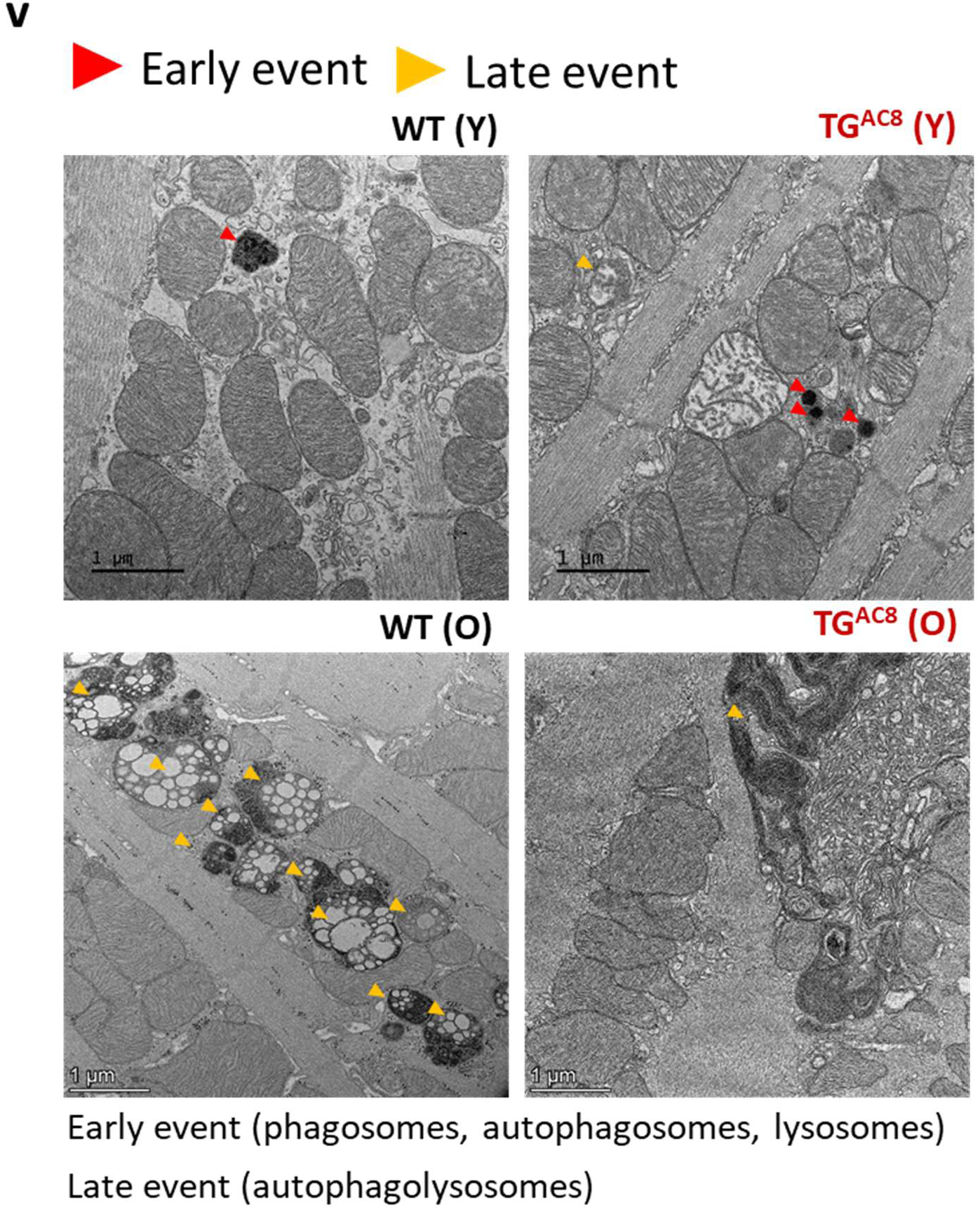
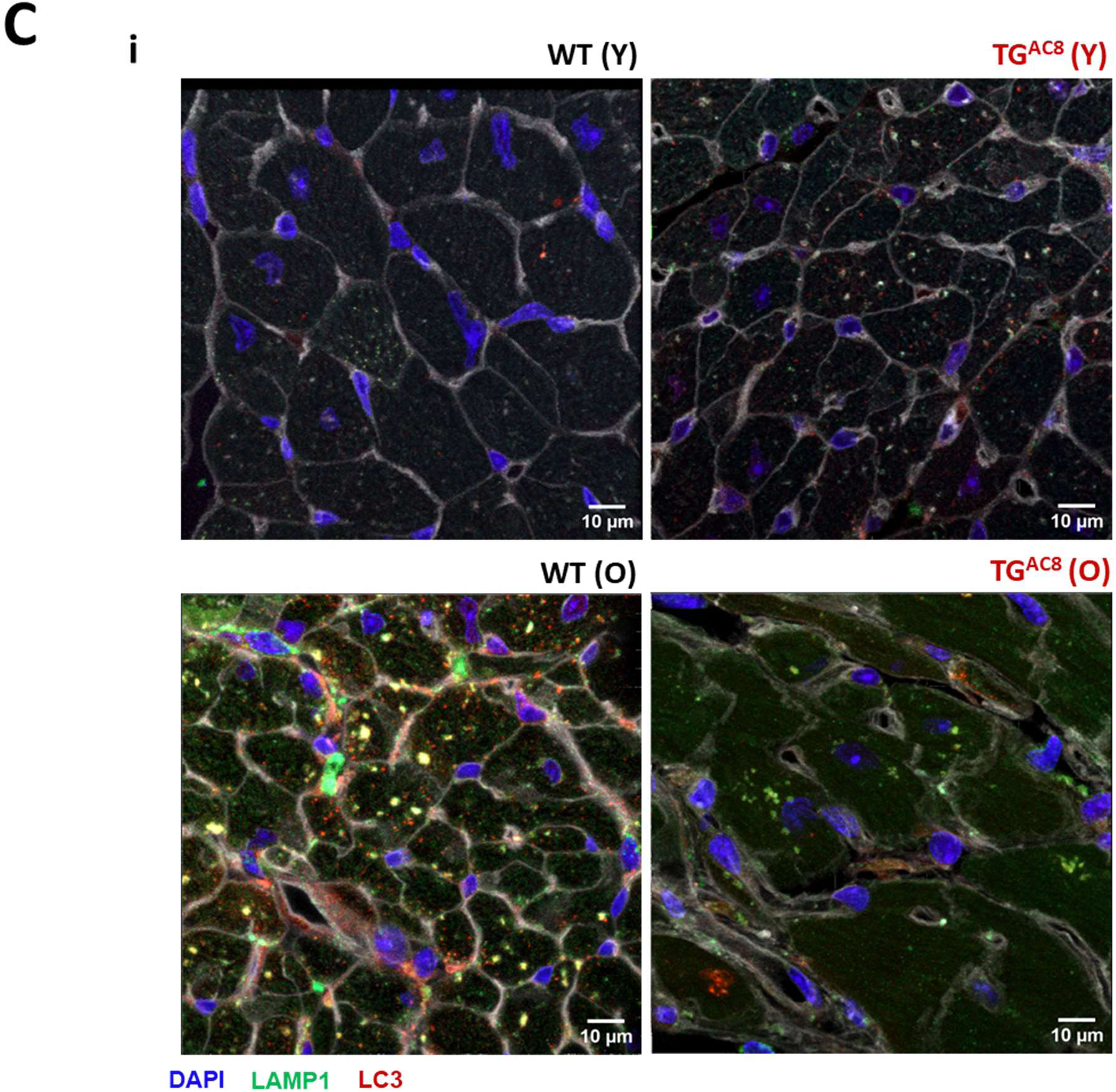
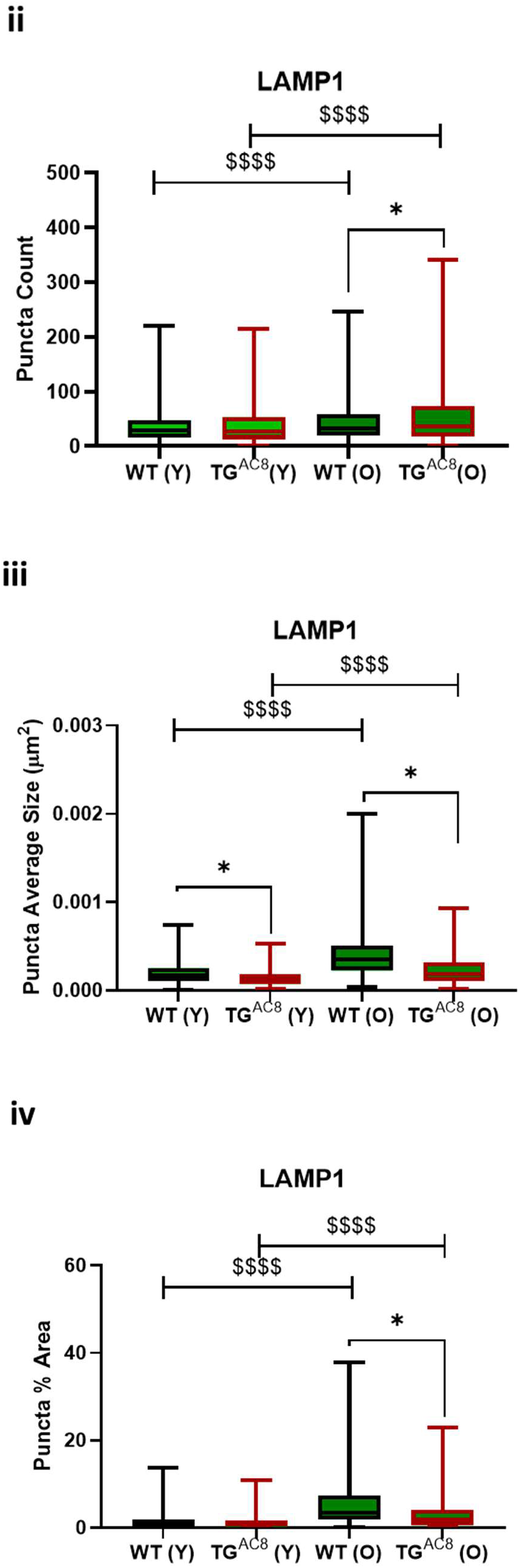
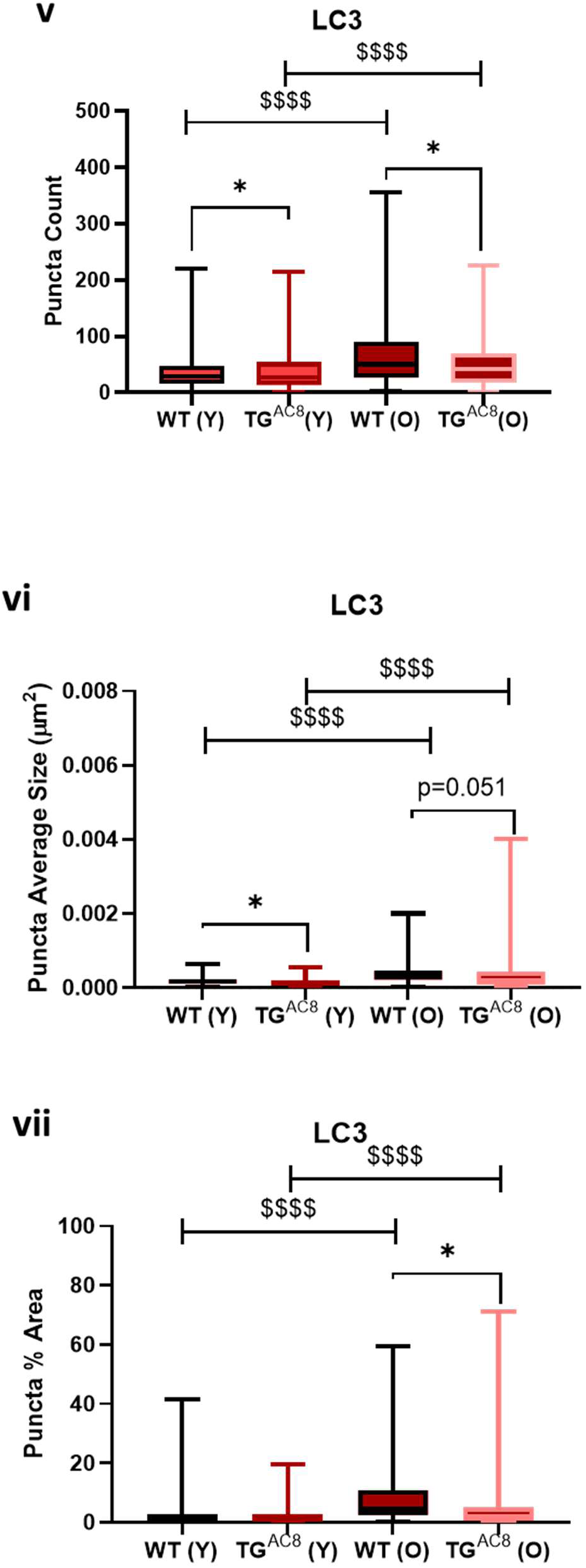
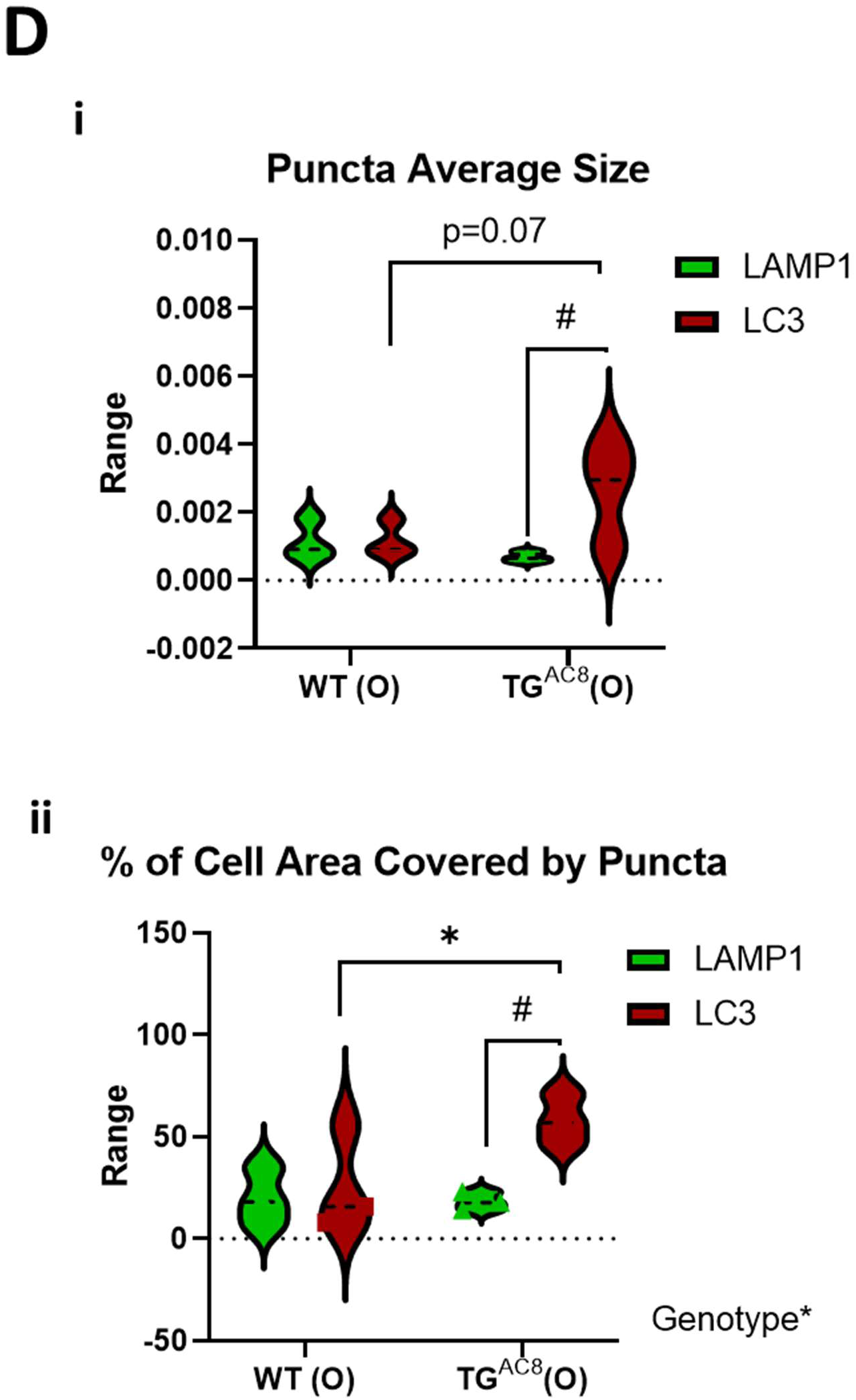
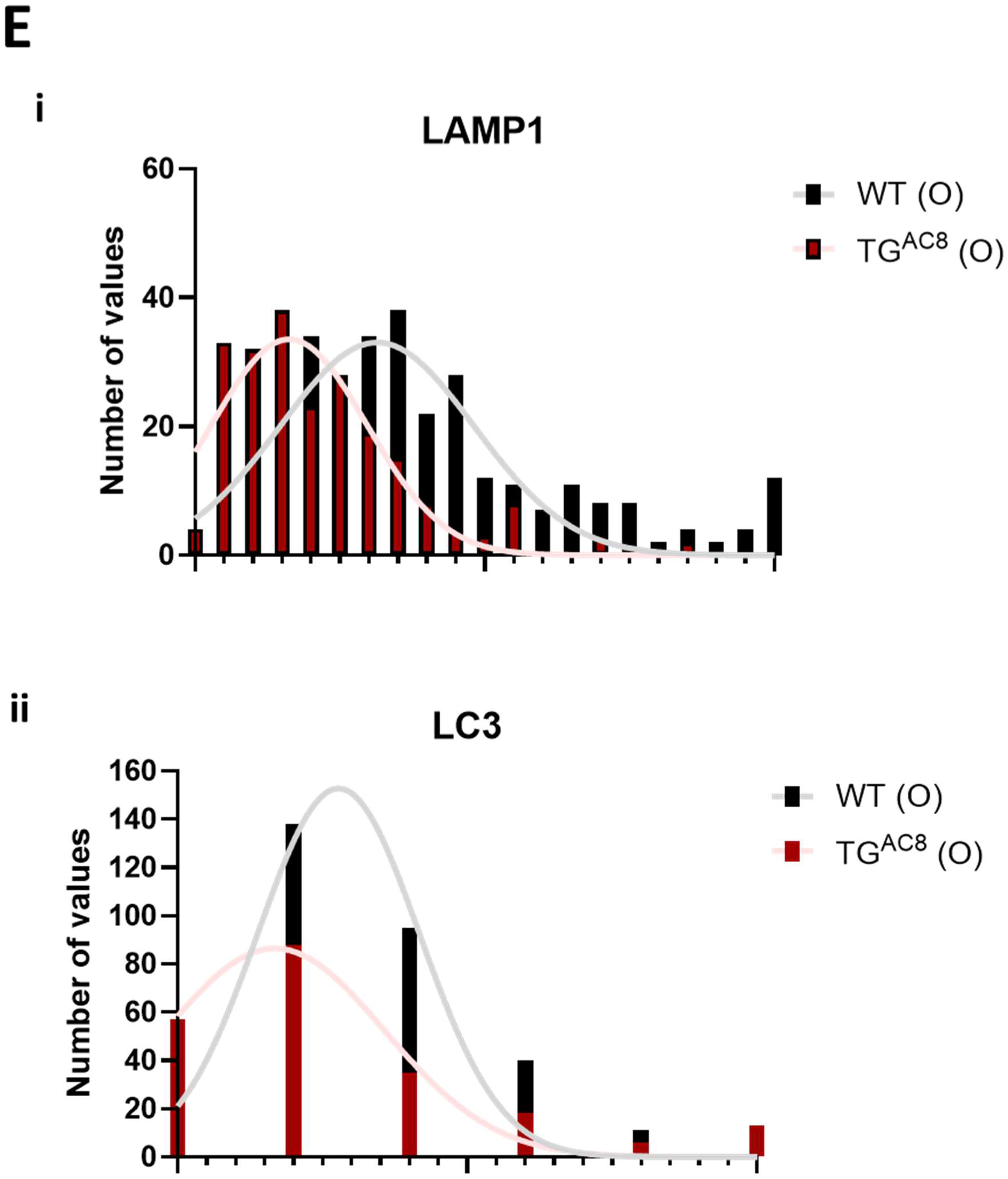
Autophagy failure in aged TG^AC8^ increases inclusions’ aggregation and size heterogeneity. (**A**) Bar graphs and Western blot analysis of autophagy markers LC3, and (**B**) bar graphs showing event quantification of macroautophagic figures (TEM) and (**C**) of endogenous LC3^+^- and LAMP1^+^-puncta (IHC) in TG^AC8^ and WT at 3-4 months (n=3-5 mice per group) and 21 months of age (n=5 mice per group) LAMP1^+^-puncta are in green, LC3^+^-puncta are in red. **Ai**), LC3A; **Aii**), LC3B. Quantification of early events **Bi**), number of per reference area; **Bii**) volume per cell volume, and quantification of late events; **Biii**), number of per reference area; **Biv**) volumes per cell volume; **Bv**) TEM representative images. **Ci**), IHC representative images; **Cii**), quantification of LAMP1^+^-puncta count; **Ciii**), quantification of LAMP1^+^-puncta average size; **Civ**) quantification of the % of cell area covered by LAMP1^+^-puncta; **Cv**), quantification of LC3^+^-puncta count; **Cvi**), quantification of LC3^+^-puncta average size; **Cvii**) quantification of the % of cell area covered by LC3^+^-puncta; **Di**), Range of LAMP1^+^- and LC3^+^-puncta average size in TG^AC8^ and WT at 21 months of age; **Dii**), Range of the % of cell area covered by LAMP1^+^- and LC3^+^-puncta in TG^AC8^ and WT at 21 months of age; **Ei-ii**) LAMP1^+^- and LC3^+^-puncta size distribution in TG^AC8^ and WT at 21 months of age. **A**: A three-way ANOVA with repeated measurements, followed by Original FDR method of Benjamini and Hochberg post-hoc multi-comparison test was used. **B**, **C**, and **D**, A two-way ANOVA, followed by Original FDR method of Benjamini and Hochberg post-hoc multi-comparison test was used. Data are presented as the median ±range in **A**, and as median ±interquartile range in all others. Symbols * $ indicate significant (p<0.05) differences: between genotypes* (WT vs TG^AC8^), between ages^$^ (young vs old) and between markers^#^ (LAMP1 vs LC3).

Assessment by fluorescence microscopy (IHC) of the number and size of endogenous LC3^+^-puncta (autophagosomes) and LAMP1^+^-puncta (lysosomes/late endosomes), (**Figure 4Ci**), in young and aged TG^AC8^ LVs and age-matched WT, revealed that puncta count and size significantly increased in *aged* compared to *young* mice (**Figures 4Cii-vii**) independently of genotype, suggesting downregulation of autophagy induction and clearance, in aged mice.

TG^AC8^, however, differed from WT in several aspects: both LAMP1^+^-puncta and LC3^+^-puncta were *smaller* in TG^AC8^ vs matched WT, at *both* ages (**Figures 4Ciii, 4Cvi**); while LC3^+^-puncta counts were higher in numbers in *young* TG^AC8^ (**Figure 4Cv**), there was no difference in the % of cell area covered by LC3^+^-puncta in young TG^AC8^ vs WT (**Figure 4Cvii**), but as the TG^AC8^ *aged*, LC3^+^-puncta count number decreased (**Figure 4Cv**), whereas LAMP1^+^-puncta counts significantly increased (**Figure 4Cii**), vs WT. Additionally, independent of counts, both LAMP1^+^-puncta and LC3^+^-puncta covered a smaller area of the cell in *aged* TG^AC8^ LV vs aged WT (**Figures 4Civ, 4Cvii**), but the latter were much more heterogeneous in size (**Figures 4Di-ii**), and had a different frequency distribution vs age-matched WT (**4Ei-ii**). A reduction in puncta size in TG^AC8^ vs WT at both ages, indicates <u>differential modulation</u> of the autophagic process; relative changes in LAMP1^+^-vs LC3^+^-puncta counts, in *aged* TG^AC8^, suggest a <u>different</u> requirement in <u>cargo detection</u> and transport (LAMP1 is also a marker of late endosomes) compared to aging WT. Nonetheless, the greater accumulation and increased aggregation (increased LC3^+^-puncta size and smaller % of cell area covered by LC3^+^-puncta) of insoluble inclusions in the old TG^AC8^ heart compared to age-matched WT, suggests that effects of aging are more severe in TG^AC8^ vs WT.

Measurement of specific LC3 isoforms, p62, and other proteins (ATG9A) involved in the autophagic process (by WB), and assessment of autophagic flux (with CQ), showed that LC3AI was significantly upregulated in old TG^AC8^, compared to age-matched WT (**Figure 5Ai**), similar to the genotypic difference at 3-4 months (**Figure 1Ai**), whereas LC3AII protein levels did not significantly differ between genotypes at 17-21 months, either in the soluble (**Figure 5Ai**) or insoluble (pellet) fraction (**Extended Data Figures S3i, S3iii**). In contrast, LC3B forms did not differ in aged TG^AC8^ (**Figure 5Aii**), demonstrating that LC3 isoform-specific modulation persists in old age, as we previously observed in TG^AC8^ at younger age (**Figure 1Ai-ii**). p62 protein levels were also significantly upregulated in old TG^AC8^ LVs, both in the soluble fraction (**Figure 5Aiii**), similar to the younger age (**Figure 3Ci**), and in the insoluble fraction (**Extended Data Figures S3ii-iii**), where *LC3AII-bound* p62 is present in LC3^+^/p62^+^ aggregates/inclusions^24^, suggesting that unprocessed protein aggregates were increased in the aged TG^AC8^, vs WT. Protein levels of ATG9A were elevated at both ages in TG^AC8^ vs age-matched WT (**Extended Data Figure S4**), but in *aged* TG^AC8^ was the highest level, suggesting increased membrane delivery to the phagophore assembly site (PAS)^37^, a process that is essential for autophagosome formation^38^, transport of lysosomal hydrolases^39^, and lipid mobilization to mitochondria^40^.

**Figure 5.**
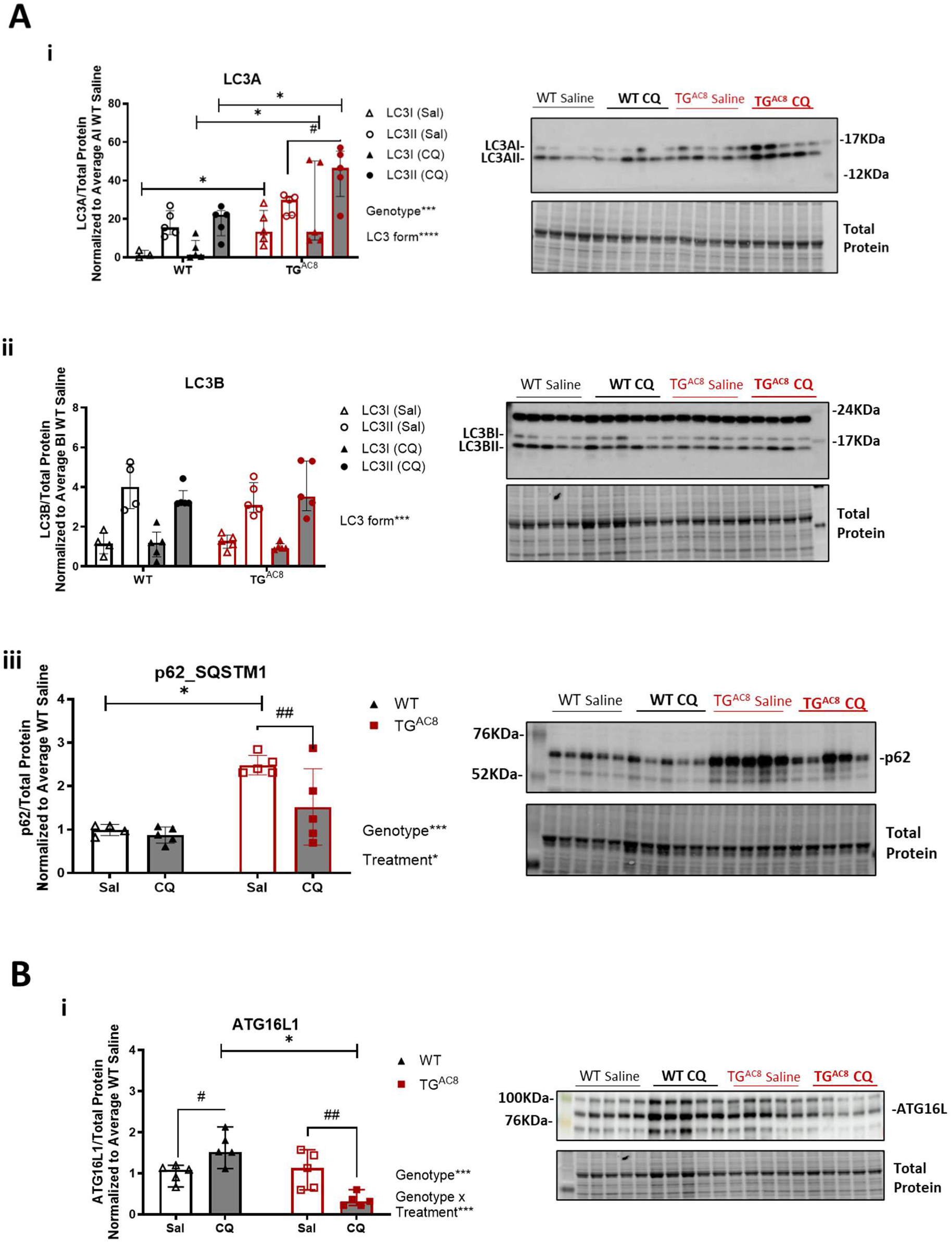
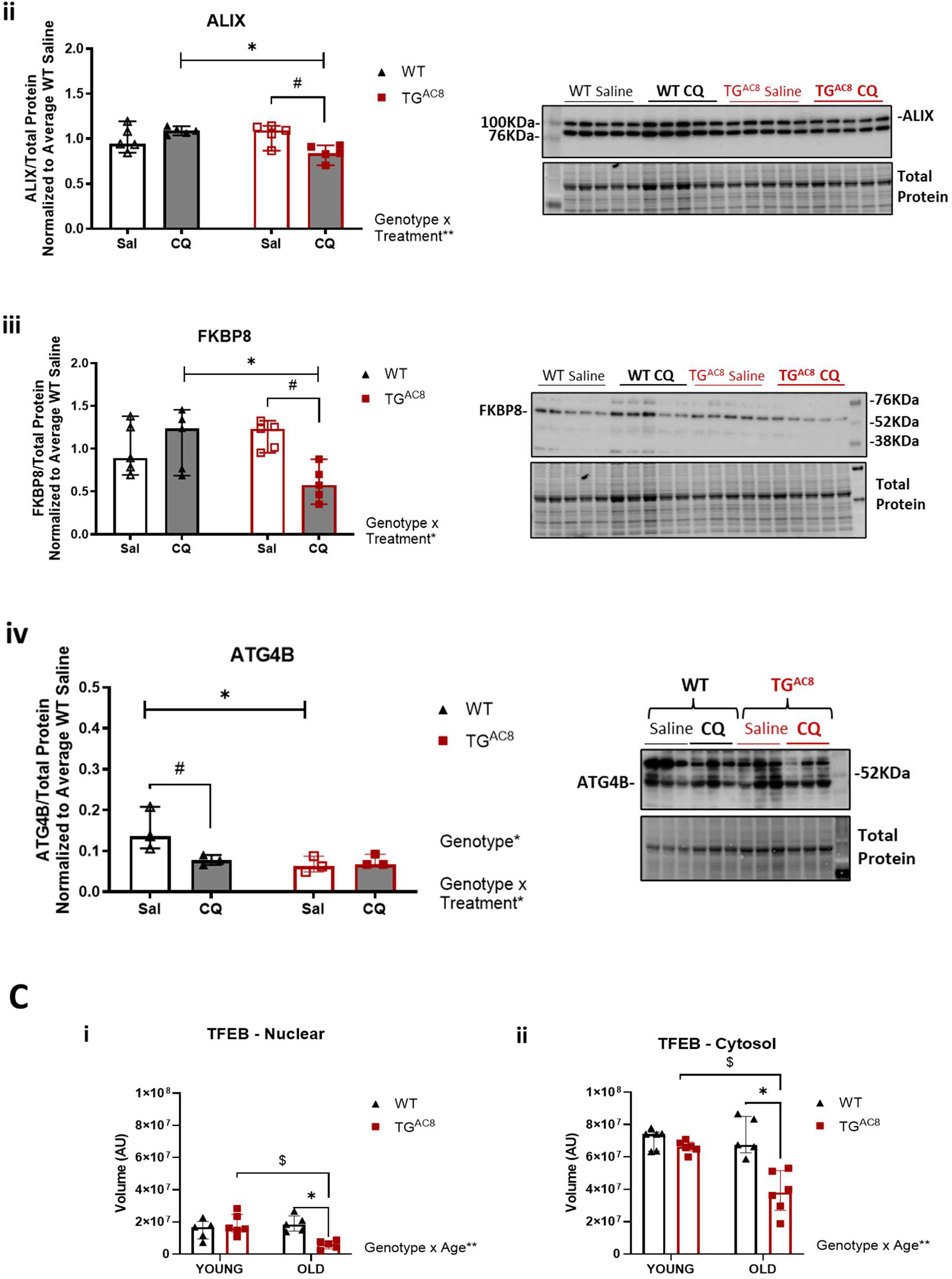
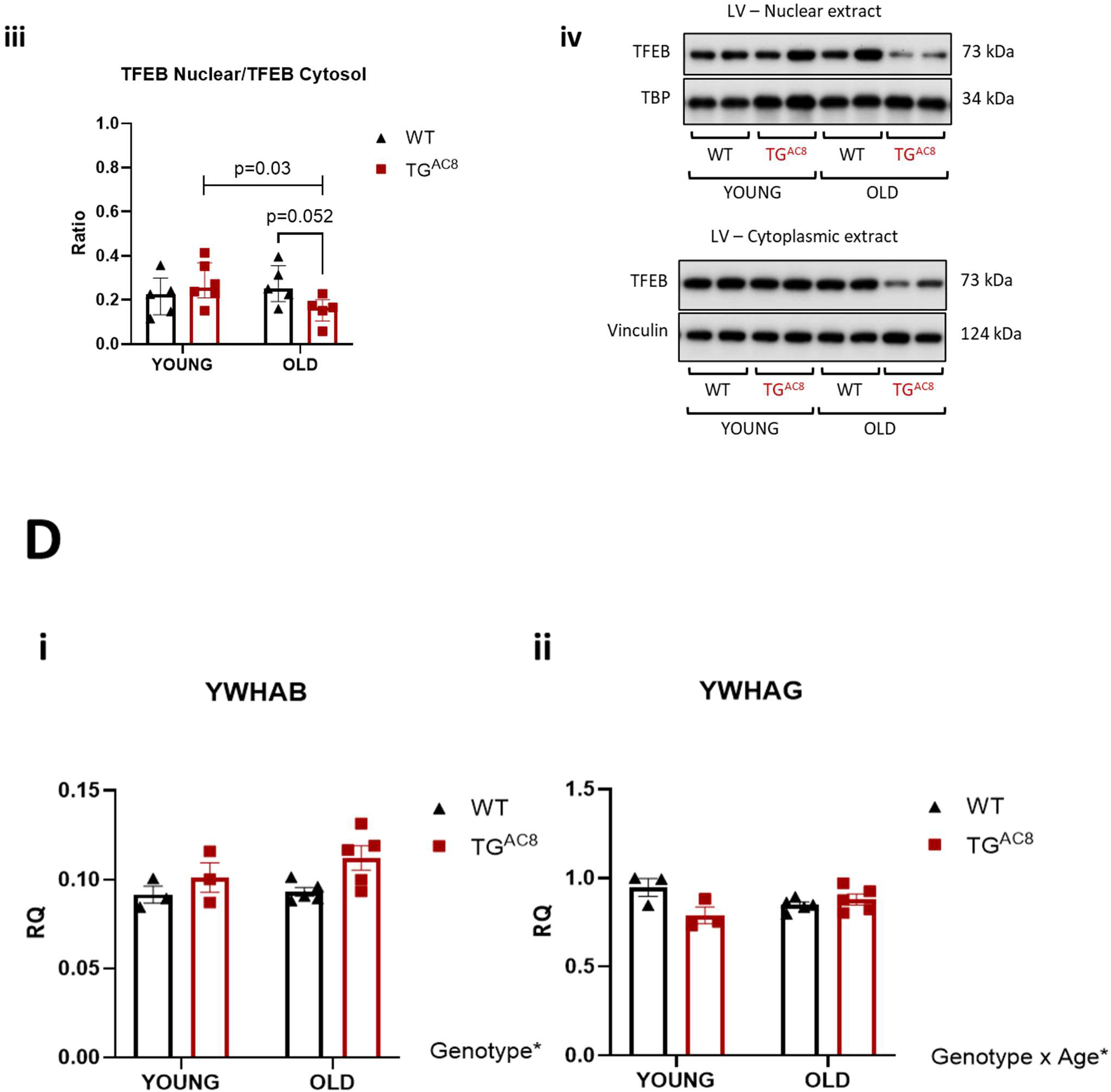
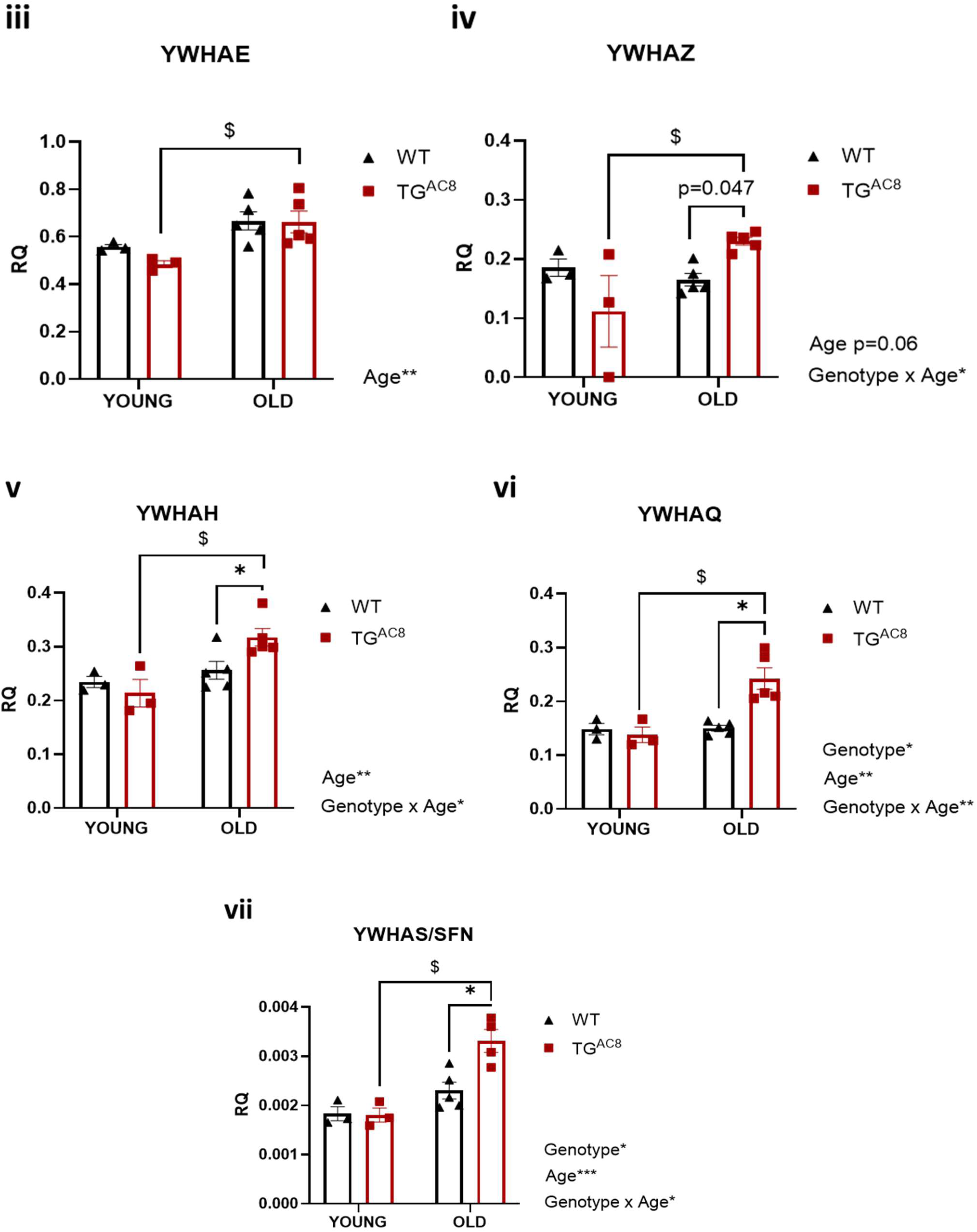
Accelerated autophagic flux in young TGAC8 heart becomes maladaptive and insufficient in old age, leading to more severe suppression of the autophagy program. **A-B** TG^AC8^ and WT littermates at 21 months of age were treated by intraperitoneal administration (IP) with chloroquine (CQ) (50 mg/kg) or saline, and LVs (n=5 mice per group) were collected 3 hours after and snap-frozen for analysis of autophagy markers LC3, p62, ATG16L1, ALIX, FKBP8 and ATG4B. Bar graphs and Western blot analysis of **Ai**), LC3A; **Aii**), LC3B; **Aiii**), total p62; **Bi**), ATG16L1; **Bii**),.ALIX; **Biii**), FKBP8; **Biv**),.ATG4B; **C** Total TFEB protein levels in cytosolic and nuclear enriched fractions in young (3-4 months of age) and old (20 months of age) TG^AC8^ and WT littermates (n=5 mice per group). **Ci**), Total TFEB in nuclear fraction; **Cii**), Total TFEB in cytosolic fraction; **Ciii**), TFEB Nuclear/TFEB Cytosol ratio; **Civ**), Representative WB images of nuclear and cytoplasmic extracts. **D** Quantification of 14-3-3s transcripts in young and old TG^AC8^ and WT littermates using the primers indicated in Supplemental Table 2 (n=3-5 mice per group). HPRT was used as housekeeping gene. **Di**), YWHAB transcript **Dii**), YWHAG transcript; **Diii**), YWHAE transcript; **Dii**), YWHAZ transcript; **Dv**), YWHAH transcript; **Dvi**), YWHAQ transcript and **Dvii**), SFN transcript. **Ai-ii**: A three-way ANOVA with repeated measurements, followed by Original FDR method of Benjamini and Hochberg post-hoc multi-comparison test was used. **Aiii, B, C and D**, A two-way ANOVA, followed by Original FDR method of Benjamini and Hochberg post-hoc multi-comparison test was used. Data are presented as the median ±interquartile range except for qPCR, which are presented as the mean ±SEM. Symbols * ^$ #^ indicate significant (<0.05) differences between: genotypes* (WT vs TG^AC8^), between ages^$^ (young vs old), between treatments^#^ (CQ vs saline).

Following CQ treatment, both LC3AI and LC3AII significantly *increased* in TG^AC8^ vs WT-saline and WT-CQ (**Figure 5Ai**); LC3AII also increased vs TG^AC8^-saline, an indication of an *accelerated carrier-flux* for <u>LC3A</u> in old TG^AC8^ vs age-matched WT. <u>LC3B</u> forms were not affected by CQ (**Figure 5Aii**), whereas CQ remarkably reduced <u>p62</u> protein levels (soluble fraction) in old TG^AC8^ (**Figure 5Aiii**), similar to younger age (**Figure 3Ci**). Additionally, the carrier-flux of <u>ATG16L1</u> (**Figure 5Bi**), of <u>ALIX</u> (**Figure 5Bii**) and of <u>FKPB8</u> (**Figure 5Biii**) were all significantly reduced in old TG^AC8^ LVs (by 68%, 20% and 49% respectively), an indication that *overall* autophagic flux in aged TG^AC8^ is significantly reduced, compared to both younger TG^AC8^ and to aging WT. In contrast, the carrier-flux of <u>ATG4B</u> (**Figure 5Biv**), LC3 phosphorylation (S12 and T50) and ACP2 protein levels (**Extended Data Figures S5i-iii**) did not differ by genotype at 17-21 months of age.

Because the *Coordinated Lysosomal Expression and Regulation* (CLEAR) network^48^ orchestrates upstream signaling pathways of autophagy, lysosome biogenesis and function^41^, we assessed protein levels and activation of its master regulator, the transcription factor TFEB, in young and old TG^AC8^ LVs and age-matched WT (**Extended Data Figure S6i-iv**). We observed a significant increase in the phosphorylation at S210 (TFEB^S2^^10^), together with an increase in the ratio of TFEB^S210^ to total TFEB (**Extended Data Figure S6iii**), with aging in both genotypes (**Extended Data Figure S6i-iv**), suggesting the lack of a genotype-effect on autophagy induction. However, and remarkably, cytoplasmic and nuclear enrichment of LV tissue showed that the nuclear to cytoplasmic ratio of total TFEB was significantly reduced in aged TG^AC8^ vs age-matched WT (**Figures 5Ci-iv**). TFEB phosphorylation at S210 (by mTORC1^42^) increases its interaction with 14-3-3 proteins in the cytoplasm, which prevents its translocation to the nucleus^42^ consequently turning off the activation of TFEB downstream pathways, including autophagy (14-3-3:RAPTOR interaction), and mitophagy/ER-phagy (14-3-3:BAD interaction, which shifts the balance towards survival, and BCL-2/BCL-XL binding to BECLIN-1 (ATG6))^43^. 14-3-3 is a family of ubiquitously expressed adaptor proteins that typically bind to “client proteins” at phosphorylated serine/threonine motifs, regulating their stability, activity and/or localization^44^. We therefore assessed 14-3-3s’ transcriptional expression (by qPCR) in TG^AC8^ vs WT with aging, and found that five (YWHAE, YWHAZ, YWHAH, YWHAQ and Stratifin) of the 7 isoforms of 14-3-3s were significantly increased in aged vs young TG^AC8^, whereas 14-3-3’s transcripts did not change in WT with age (**Figures 5Di-vii**). Additionally, mRNA of YWHAZ, YWHAH, YWHAQ and Stratifin were also significantly increased in aged TG^AC8^ vs old WT (**Figures 5Div-vii**). Transcriptional upregulation of 14-3-3s together with reduced levels of nuclear TFEB explain the greater reduction in the activation of the CLEAR network in old TG^AC8^ vs WT hearts, strongly suggesting that autophagic signaling becomes <u>more severely downregulated</u> in TG^AC8^ with aging, compared to that in age-matched WT.

Taken together, data in figures suggest that autophagy declines with age in TG^AC8^ and WT regardless of genotype, as shown by reduced protein levels of markers involved in autophagy induction (LC3, ATG4B), accumulation of LC3II, increased puncta count and size, and reduced numbers of *early* autophagic events in conjunction with higher numbers/volumes of *late* figures.

However, further downregulation of the CLEAR network in old TG^AC8^ LVs compared to age-matched WT, together with increased heterogeneity in LC3^+^-puncta size, the reduction in the cell area covered by LC3^+^-puncta and the different modulation pattern of LAMP1^+^-puncta and LC3^+^-puncta in old TG^AC8^, all point to much more severe autophagy downregulation in old TG^AC8^ vs age-matched WT, thus that the *optimized, highly efficient autophagic flux* in young age that accommodated proper cargo clearance in TG^AC8^, *manifests reduced kinetics and becomes dysfunctional* as TG^AC8^ ages, and would clearly be insufficient for proper clearance of aggregates and inclusions of aberrant size, and more prone to aggregation, compared to the young-TG^AC8^ and age-matched WT.

### <u>Proteasome insufficiency</u> and <u>mitochondrial dysfunction</u> lead to exaggerated age-associated increase in aggregates accumulation in TG^AC8^

We next assessed PQC mechanisms (ubiquitin proteasome system (UPS) activity, autophagy and mitophagy) and evaluated protein translation rates and aggregates accumulation in young and aged TG^AC8^, and age-matched WT. Our previous work in young mice (3-4 months of age) demonstrated that: 1) UPS activity and autophagy, including mitophagy, were all upregulated in TG^AC8^, compared to WT, indicating cooperation among PQC mechanisms^14^; 2) although the protein synthesis rate was 40% higher in TG^AC8^ vs WT, UPS clearance of soluble aggregates was also upregulated, and insoluble protein aggregates did not accumulate within cells^14^; 3) the numbers of healthy mitochondria and the percent of cell volume these occupied, did not differ between TG^AC8^ and WT^14^; and 4) although the canonical cargo receptors PARKIN was significantly upregulated, mitochondrial fitness was comparable between genotypes^14^.

Proteasome activity was not increased in old TG^AC8^, compared to age-matched WT and in contrast to younger age (**Figure 6A**). UPS function in TG^AC8^ declined with aging, from 3 to 18 months, whereas it remained the same with aging in WT (**Figure 6A**). The density of *soluble* aggregates also did not differ by genotype at both ages, but it decreased with aging independent of genotype (**Figure 6Bi**). The ratio of insoluble to soluble aggregates, however, significantly increased in TG^AC8^ between 3 and 18 months, with no change in aging WT (**Figure 6Bii**), indicating the greater accumulation of *insoluble* protein aggregates in the TG^AC8^ heart, during the aging process, compared to “normal” aging WT.

**Figure 6.**
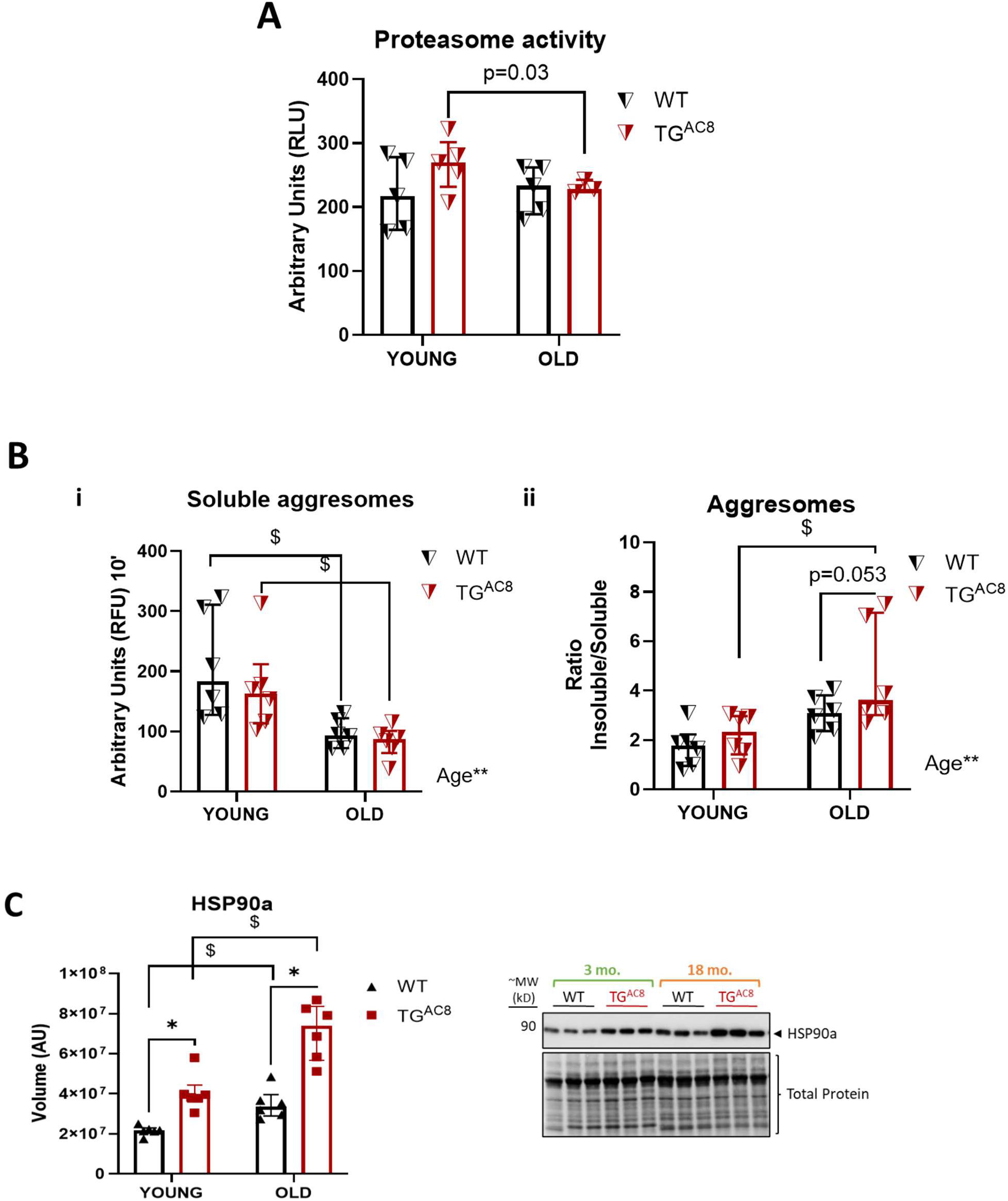
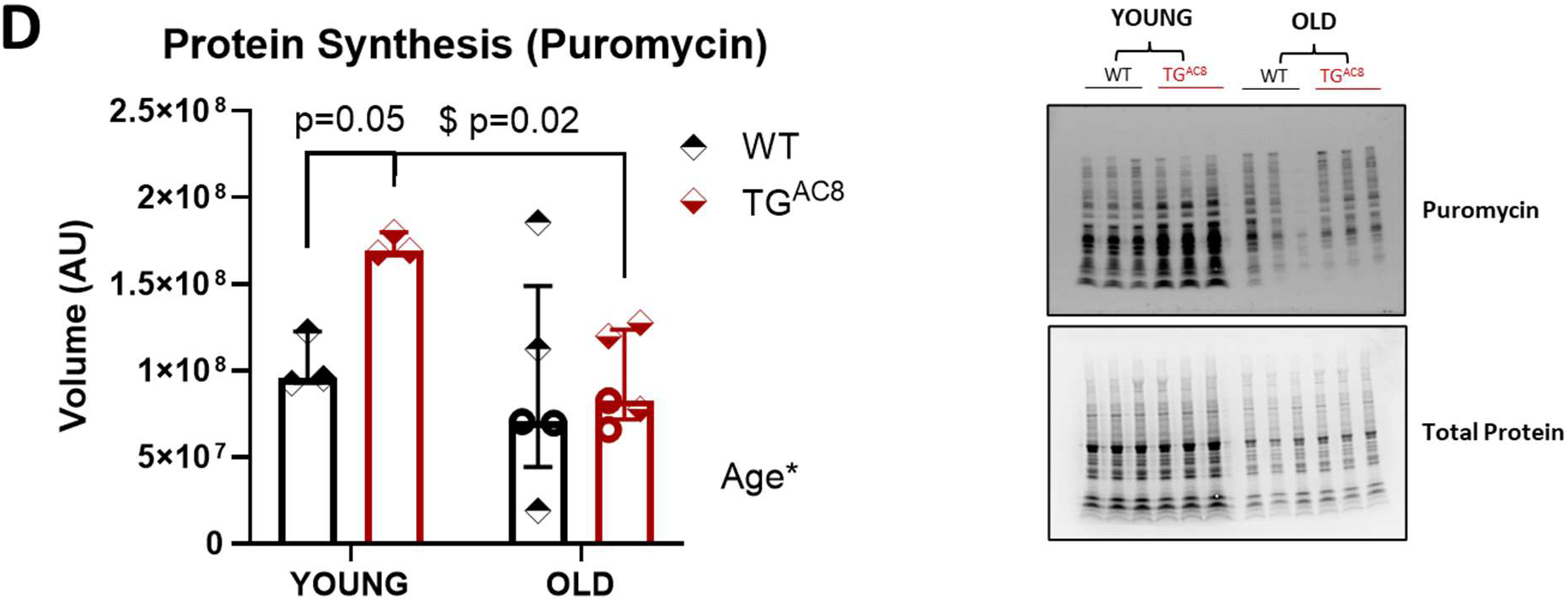
Proteostasis dysregulation is more severe in old TGAC8 heart. Bar graphs of **A**), Proteasome activity assessment and of **B**), Aggregates quantification in soluble fractions and the ratio of insoluble vs soluble protein aggregates in young (3-4 months) and old (17-18 months) TG^AC8^ and age-matched WT (n=3-5 per group). **C**) Bar graphs and Western blot analysis of the stress marker HSP90. **D**) TG^AC8^ and WT at 19 months of age (n=5 mice per group) were treated by intraperitoneal administration (IP) with puromycin (0.022g/g mouse) according to the SUnSET method, and LVs were collected 30 minutes after drug administration and snap-frozen for puromycin analysis. **A-D**, A two-way ANOVA, followed by Original FDR method of Benjamini and Hochberg post-hoc multi-comparison test was used. Data are presented as the median ±interquartile range. Symbols * ^$ #^ indicate significant (<0.05) differences between genotypes* (WT vs TG^AC8^), between ages^$^ (young vs old).

Because accumulation of *insoluble* aggregates leads to proteotoxic stress^45^, we assessed, by WB, protein levels of heat shock protein 90 (HSP90), a stress-sensor that, with its chaperon activity^46^ activates protective mechanisms to reduce stress and maintain cellular homeostasis and intracellular transport^47^. HSP90α, the HSP90 stress-inducible form, significantly increased between 3 and 18 months in both genotypes (**Figure 6C**), an indication that age, *per se*, increases cellular stress, as well as the need for increased protein folding/refolding, maintenance and degradation. Additionally, HSP90α was significantly upregulated in TG^AC8^ (**Figure 6C**) vs WT at both ages^14^, but was increased to a further extent in old TG^AC8^, in parallel with the highest level of proteotoxic stress (**Figure 6Bii**), and the persistent higher cAMP-derived chronic stress of the TG^AC8^ heart, compared to “normal’ aging WT. Although protein synthesis did not differ by genotype in aged mice (**Figure 6D**), protein translation rates, a hallmark of aging in health^48^, significantly decreased in TG^AC8^, from 3 to 18 months.

Mitochondrial failure, a hallmark of aging, occurs in diverse pathologic diseases conditions, including heart failure^49^, in response of a wide variety of stressors. We evaluated protein levels (by WB) of key players of mitochondrial dynamics, including fusion/fission^50^, function^51^, integrity^52^ and clearance^53^ in young and aged TG^AC8^ LVs, and age-matched WT. *Mitofusin 1* (MFN1), located on the outer mitochondrial membrane (OMM) and involved in the orchestration of OMM fusion, was significantly increased in old TG^AC8^ (**Figure 7A**), suggesting that conditions for mitochondrial fusion and docking^54^ were more favorable, compared to old WT; however, levels of the mitochondrial GTPase *dynamin-related protein 1* (DRP1), a cytosolic protein that controls late stages of the mitofission process^55^, and is recruited to the mitochondrial surface in response to various physiological cues^56^, did not differ between genotypes (**Figures 7B**). In contrast, levels of the protein *dynamin-related GTPase optic atrophy type 1* (OPA1), which is localized in the mitochondrial intermembrane space on the outer leaflet of the inner membrane (IMM)^57^ differed in TG^AC8^ vs WT. The expression of both the full length OPA1 (L-OPA1), which has a role in IMM fusion and cristae remodeling^58^, and of its cleaved form, S-OPA1, which is associated with mitochondrial fission^59^, and results from the proteolytic processing of L-OPA1 constitutively (YME1L^60^), and upon stress-activation (metalloprotease OMA1^61^), were significantly reduced in TG^AC8^ vs WT independent on age (**Figure 7Ci-ii**). Remarkably, although TG^AC8^ expressed less S-OPA1 vs WT (**Figure 7Cii**), the ratio of S-OPA1 to L-OPA1 was significantly elevated in TG^AC8^ vs WT at both ages (**Figure 7Ciii**), and was increased to a further extent in aged TG^AC8^ vs old WT, suggesting that persistent chronic cardiac cAMP-derived stress in the TG^AC8^ heart exacerbates the age-related increase in cellular stress^61^ (**Figure 7Ciii**). Protein levels of the mitochondrial heat-shock protein 60 (mtHSP60/HSPD1), an indispensable chaperonin that regulates mitochondrial protein homeostasis and function by preventing the aggregation of misfolded proteins^62^, were also reduced in aged TG^AC8^ vs WT (**Figure 7D**), which in the context of imbalanced levels of OPA1 and the persistent chronic cAMP-derived cardiac stress, show that decreased protection^52^ of the mitochondrial matrix and its associated functions^63^ in the TG^AC8^ heart as age advances vs age-matched WT, contribute to the greater mitochondrial dysfunction, eventually leading to heart failure^51^.

**Figure 7.**
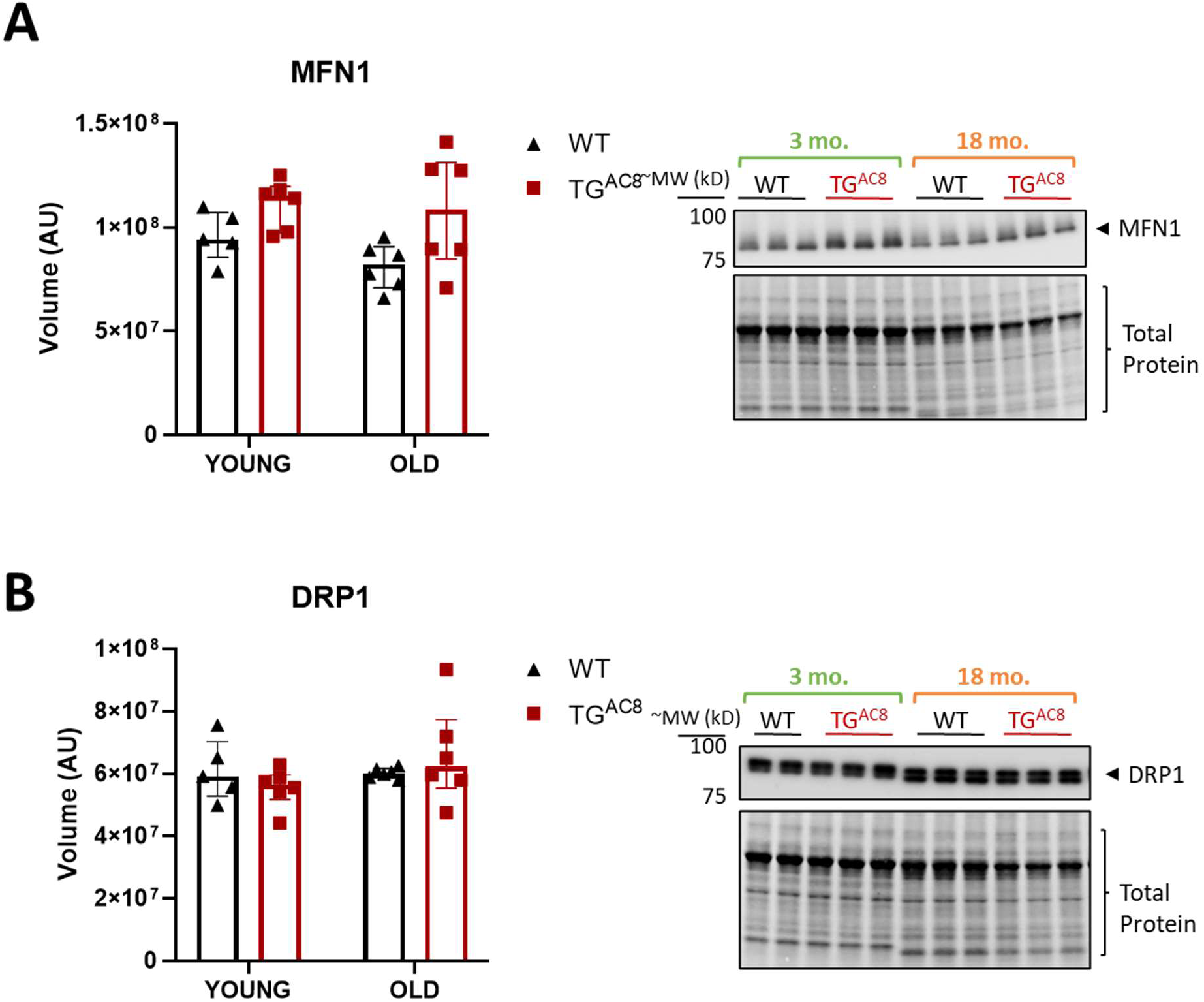
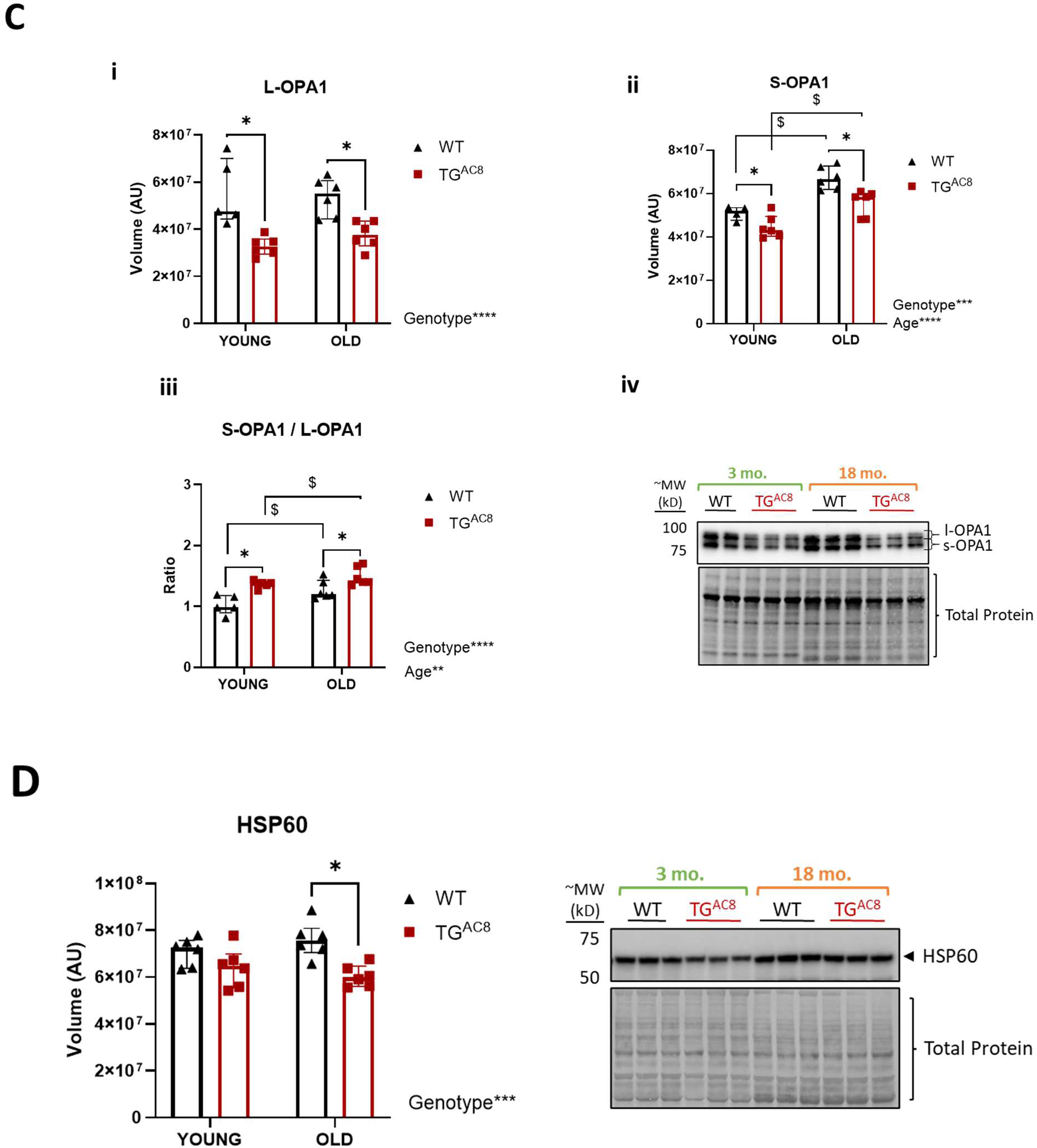
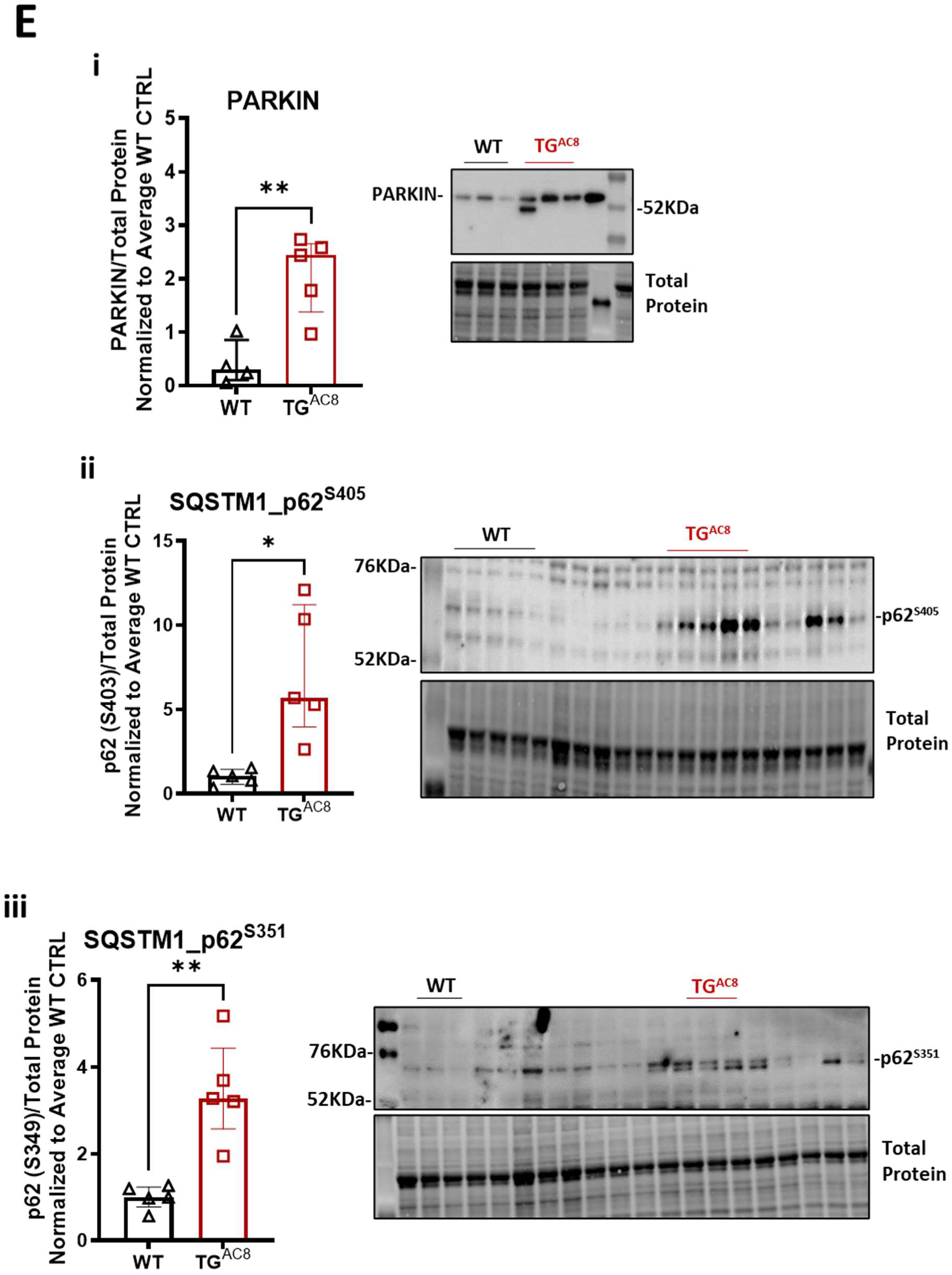
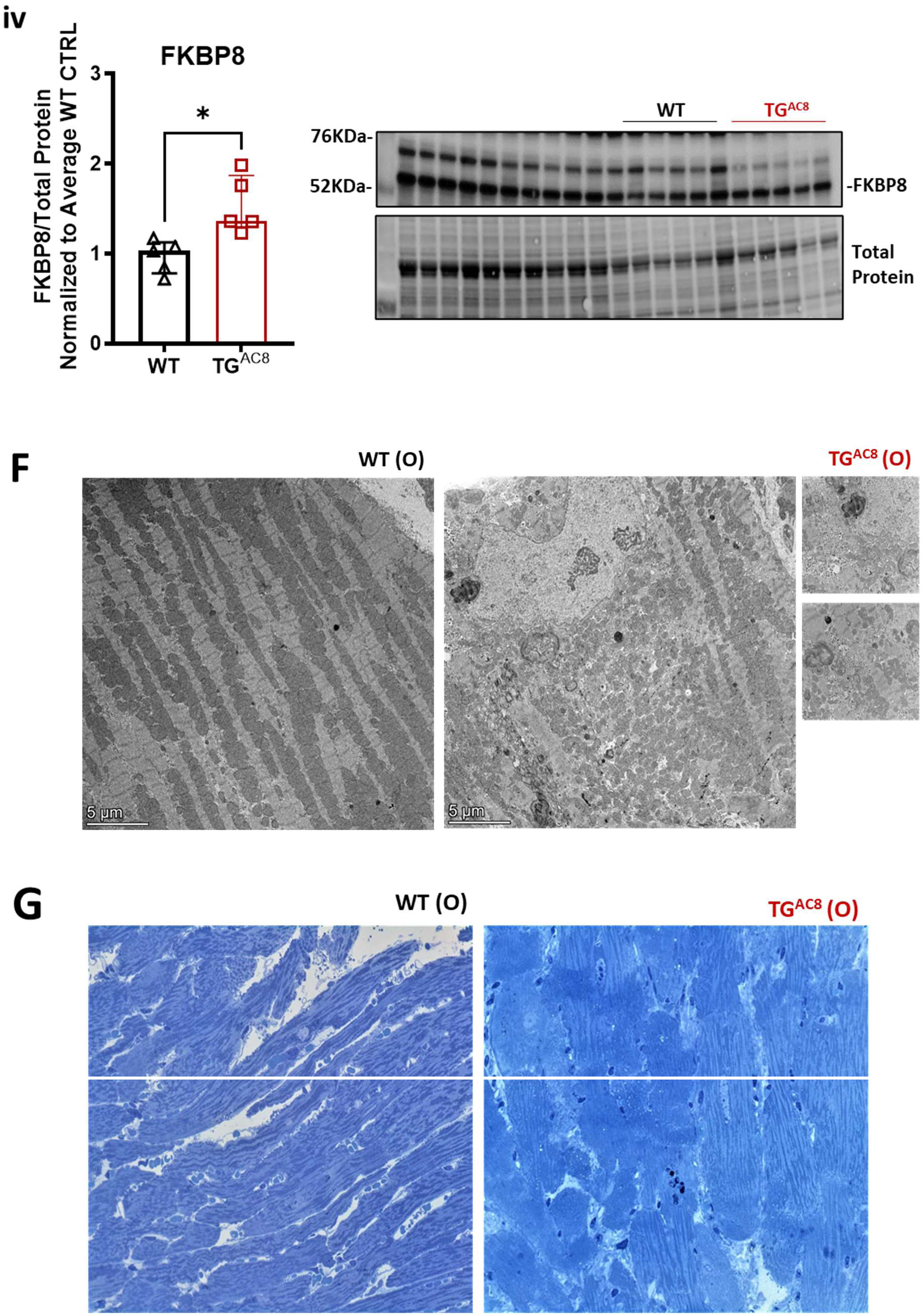
PQC insufficiency leads to mitochondria dysfunction and disruption of the mitochondrial network in old TGAC8 heart. Bar graphs and Western blot analysis of mitochondrial markers in young (3-4 months) and old (19 months) TG^AC8^ and WT littermates (n=5 mice per group). **A**), MFN1; **B**), DRP1; **Ci**) L-OPA1; **Cii**) S-OPA1; **Ciii**) S-OPA/L-OPA1 ratio; **Civ**) representative WB images; **D**) HSP60; **Ei**) PARKIN; **Eii**) p62^S405^; **Eiii**) p62^S351^; **Eiv**) FKBP8; **F**) representative TEM and **G**) light microscopy images of semithin sections showing disrupted mitochondrial network. **A-D**: A two-way ANOVA, followed by Original FDR method of Benjamini and Hochberg post-hoc multi-comparison test was used. **E**: Unpaired 2-tailed Student t test with Welch’s correction was used. Data are presented as the median ±interquartile range. Symbols * ^$^ indicate significant (<0.05) differences between genotypes* (WT vs TG^AC8^), between ages^$^ (young vs old).

Because the mitophagy process ensures disposal of malfunctional and/or dysfunctional mitochondria, we next evaluated the expression of specific cargo receptors involved in canonical and non-canonical mitophagy^64^ in aged TG^AC8^ and age-matched WT. Similar to young TG^AC8^, the canonical cargo receptors PARKIN (**Figure 7Ei**), p62^S405^ (**Figure 7Eii**), recruited to depolarized^65^ and polyubitiquinated^26^ mitochondria, respectively, and p62^S351^ (**Figure 7Eiii**), upregulated in conditions of oxidative stress^27^, were all significantly increased in aged TG^AC8^ vs WT, suggesting that canonical mitophagy is incremented in TG^AC8^ vs WT. In contrast to young TG^AC8^ (**Figure 3Biv**), the non-canonical cargo receptor FKBP8^66^, which assists LC3A during mitophagy and in the clearance of misfolded proteins, was also significantly upregulated (**Figure 7Eiv**) in the old TG^AC8^ vs WT. In line with these results, TEM and light microscopic assessment of semi-thin sections of LV cardiomyocytes clearly demonstrated disruption of the mitochondrial network in old TG^AC8^ vs old WT (**Figures 7F-G**). Reduced signaling for disposal of oxidized lipoproteins, in old TG^AC8^, together with disruption of the mitochondrial network, suggests that mitophagy failed in old TG^AC8^, because the rate of <u>damaged mitochondria accumulation</u> *exceeded* the <u>rate of aggregates clearance</u>. In other terms, despite the concurrent activation of both canonical and non-canonical mitophagy in old TG^AC8^, the system cannot cope with the increased mitochondrial dysfunction in old TG^AC8^.

LF body quantification of the extent of accumulation of undigested material (in Palade’s-stained semi-thin sections by light microscopy) (**Figure 8A**), showed that there was no difference in LF number (**Figure 8Bi**) or average body size (**Figure 8Bii**) in old TG^AC8^ vs WT. However, there was a marked heterogeneity within the LF population, and increased total volume of LF (**Figures 8Biii, 8Biv**), with LF covering a larger percentage of the cardiomyocyte area (**Figure 8Bv**), in TG^AC8^ vs WT. Interestingly, those vast insoluble aggregates reminded a myelin-like figure, with high electron-dense concentric lamellations, and displayed different pigmentation (brownish to blueish-black) (**Figure 8C**), and very heterogenous LC3^+^-puncta/inclusions (**Figure 8D**) almost entirely covering the cardiac myocyte cell area (LC3^+^-CMs) (**Figures 8Ei-iii**), and were only present in TG^AC8^ LVs, suggesting the presence of an “aged” (further oxidized) cargo accumulation of different lipoproteins and/or a denser composition.

**Figure 8.**
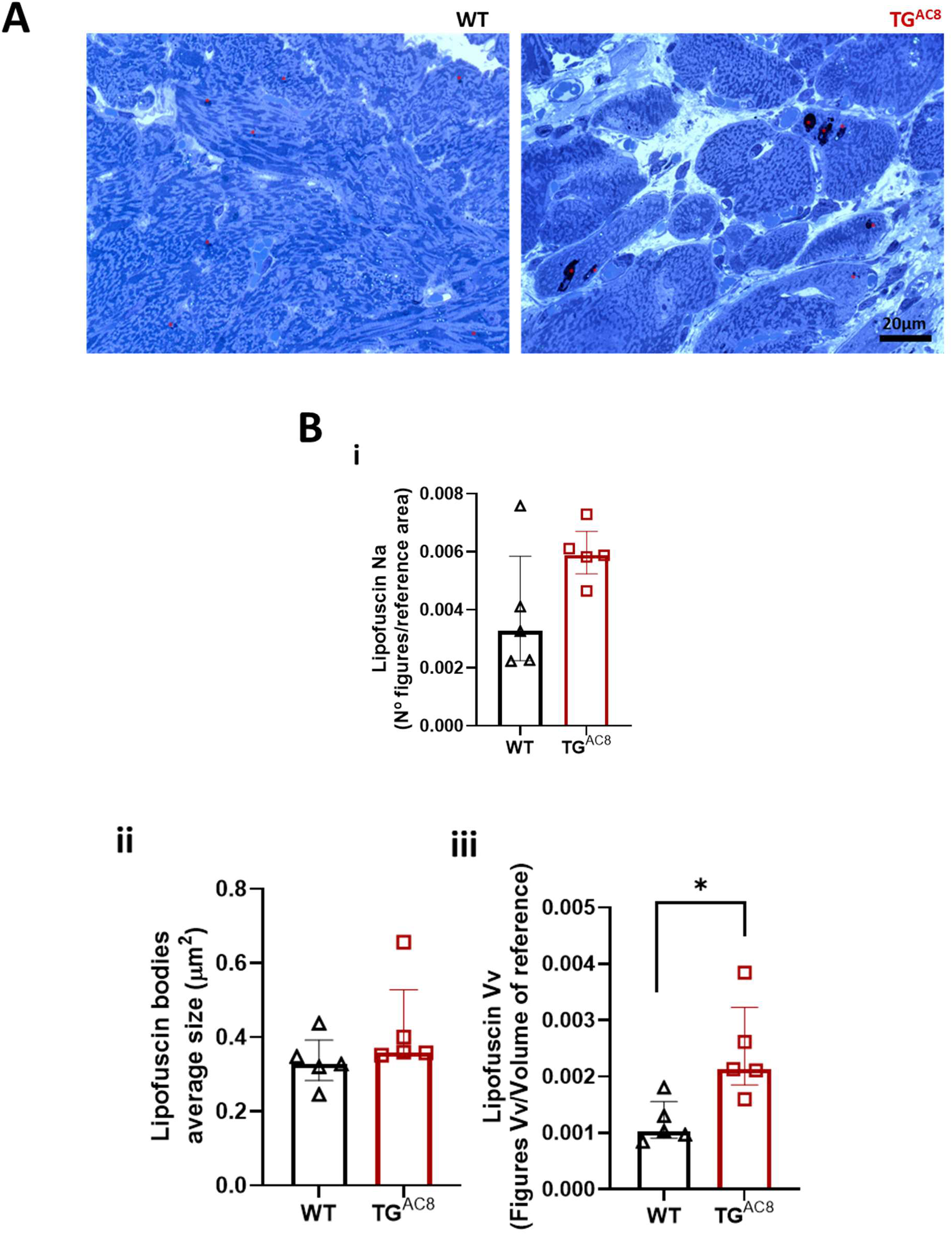
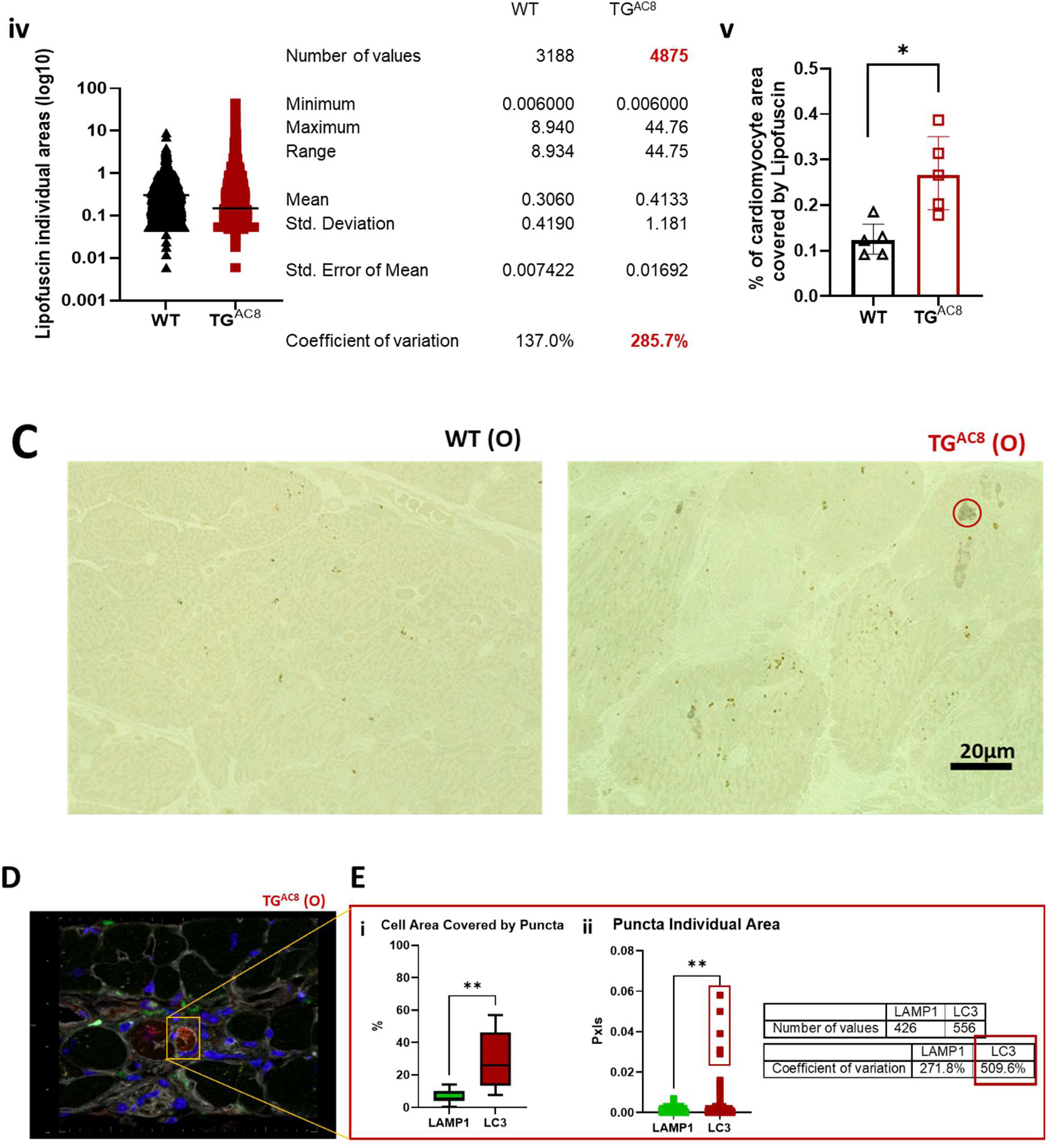
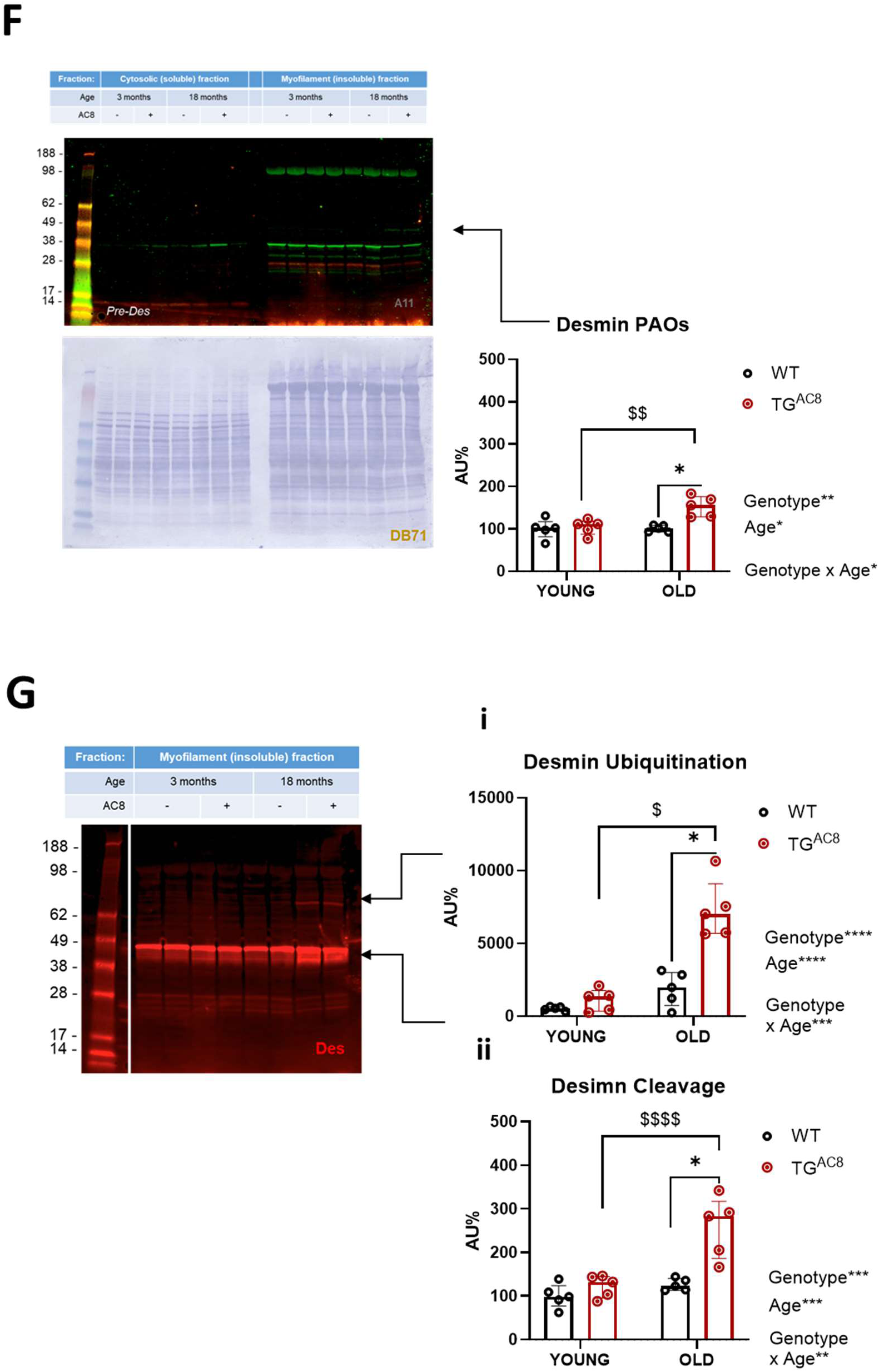
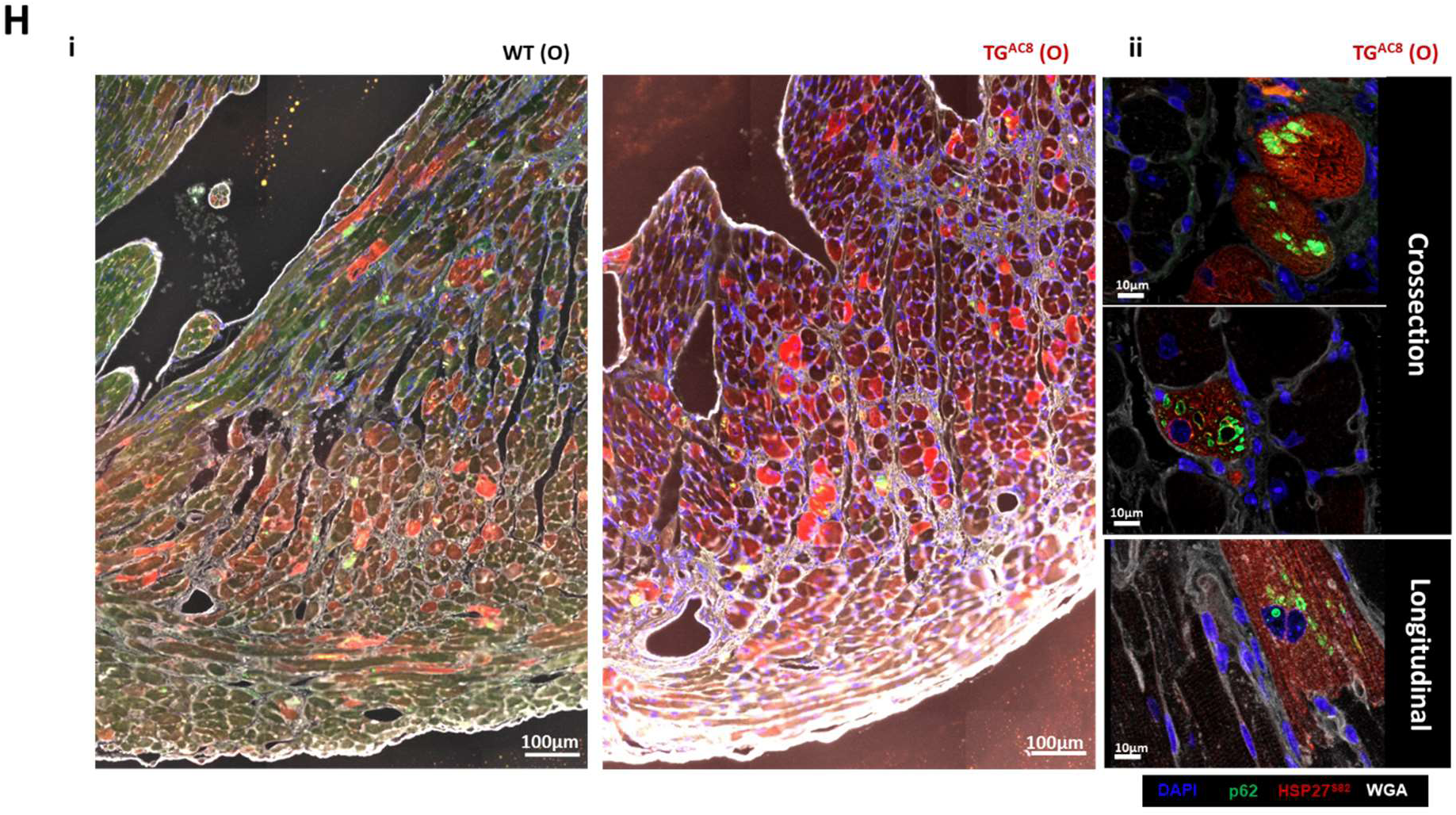
Proteostasis dysregulation from PQC insufficiency leads to increased lipofuscin accumulation and deposition of desmin Pre-Amyloid Oligomers (PAOs) in aged TGAC8. Bar graphs representing the quantification of undigested material (LF and desmin) in old TG^AC8^ and WT LVs (n=5 per group); **A**), representative semi-thin staining of lipofuscin bodies; **Bi**), number of lipofuscin figures per reference area; **Bii**), average size of lipofuscin bodies; **Biii**), volume of figures per volume of reference area; **Biv**), violin plot showing the more aberrant sizes of lipofuscin bodies; **Bv**), percentage of cardiomyocyte area covered by lipofuscin; **C**), representative image of Palade-staining of TEM semi-thin sections showing the presence of brown-to-blueish/black pigments in old TG^AC8^; **D**), representative image of IHC sections showing cardiomyocytes covered with LC3^+^-inclusions in old TG^AC8^; **Ei**), quantification of cell area covered by LAMP1^+^- and LC3^+^-inclusions in old TG^AC8^; **Eii**), quantification LAMP1^+^- and LC3^+^-puncta individual areas in old TG^AC8^; **F**), soluble and insoluble (myofilament-enriched) fractions probed with the A11 antibody for pre-amyloid oligomers (top), and total protein staining with Direct Blue71/DB71 (bottom), with relative densitometry analysis; **G**), insoluble (myofilament-enriched) fractions probed with the desmin antibody, and relative densitometry for **i**) ubiquitinated and **ii**) cleaved desmin proteoforms; **H**), representative IHC staining old TG^AC8^ and WT littermates with HSP27^S86^ (red), p62 (green), and wheat germ agglutinin WGA (white) while nuclei are in blue (DAPI); **B-E**: Unpaired 2-tailed Student t test with Welch’s correction was used. **F-G**: A two-way ANOVA, followed by Original FDR method of Benjamini and Hochberg post-hoc multi-comparison test was used. Data are presented as the median ±interquartile range. Symbols * ^$^ indicate significant differences (p<0.05) between genotypes* (WT vs TG^AC8^), between ages^$^ (young vs old).

Recent evidence has shown the involvement of desmin, whose filaments interlinks the contractile myofibrillar apparatus to mitochondria, nuclei and the sarcolemma, to impaired mitochondrial fission^67^ and proteostasis^67^. Furthermore, desmin disorganization has been documented to be a component of the accumulation of preamyloid oligomer (PAO) aggregates in the heart^68^. Desmin PAOs were increased with age in TG^AC8^ but not in WT, and were higher in aged TG^AC8^ vs WT (**Figure 8F**). Further, a higher percentage of desmin was tagged for ubiquitination (**Figure 8Gi**), and cleaved (**Figure 8Gii**), an indication of severe alterations of the myofilament ultrastructure in the old TG^AC8^ heart vs WT.

HSP27^S86^ (S82 in human), the stress-induced phosphoform (by MAPK^69^) of the small heat shock protein (sHSP) HSP27/HSPB1, which is a biomarker of cardiac damage^70^ and is upregulated during stress conditions to maintain myocardial function^71^, was significantly elevated in old TG^AC8^ vs WT LVs (**Figures 8Hi**), and specifically co-localized (via fluorescence microscopy IHC) with p62^+^-puncta/inclusions (**Figure 8Hii**), suggesting higher stress and myofilament disruption, exactly at the site of electron-dense aggregates accumulation, in aged TG^AC8^ cardiac myocytes vs old WT (**Figure 8Hii**). Thus, damaged TG^AC8^-cardiac myocytes (HSP27^S86+^) present higher garbage accumulation (p62^+^ aggregates), have a thinner wall and a sclerotic stroma, compared to old WT. Therefore, failure of PQC mechanisms leads to poor cardiomyocyte “health”, a phenotype consistently associated with the context of cardiomyopathy.

### <u>Increased failure</u> of PQC mechanisms <u>exacerbates</u> cardiac aging in TG^AC8^

We next addressed what changes occurred in cardiac structure and function in the context of dysregulated proteostasis. To this end, assessment of cardiac function and structure by echocardiogram revealed that the small hyperdynamic heart at 3 months of age in the AC8 group (significantly smaller LV chamber size and higher ejection fraction (EF) and heart rate (HR), compared to young WT)^14^, became a hypertrophic and dilated heart at 19 months of age (**Figure 9Ai-iii**). Although the HR remained higher throughout life in TG^AC8^ vs WT (**Figure 9Bi**), in aged TG^AC8^ the EF started to decline (**Figure 9Bii**) between 14 and 19 months of age, whereas the left ventricular (LV) mass (**Figure 9Ci**), the end diastolic volume (EDV) (**Figure 9Cii**) and the end systolic volume (ESV) (**Figure 9Ciii**) increased, demonstrating a reduced heart performance, vs old WT. Interestingly, some hearts were more dilated and had a greater reduction in EF than others within the aged TG^AC8^ group (**Figures 9Aii-iii, 9Bii**). This heterogeneity in LV reduced function matches the increased heterogeneity present in aggregates (**Figures 4C-E, 6Bii**) and oxidized lipoprotein accumulation in old TG^AC8^ (**Figures 8B, 8E**), compared to old WT (**Figure 9Ai**).

**Figure 9.**
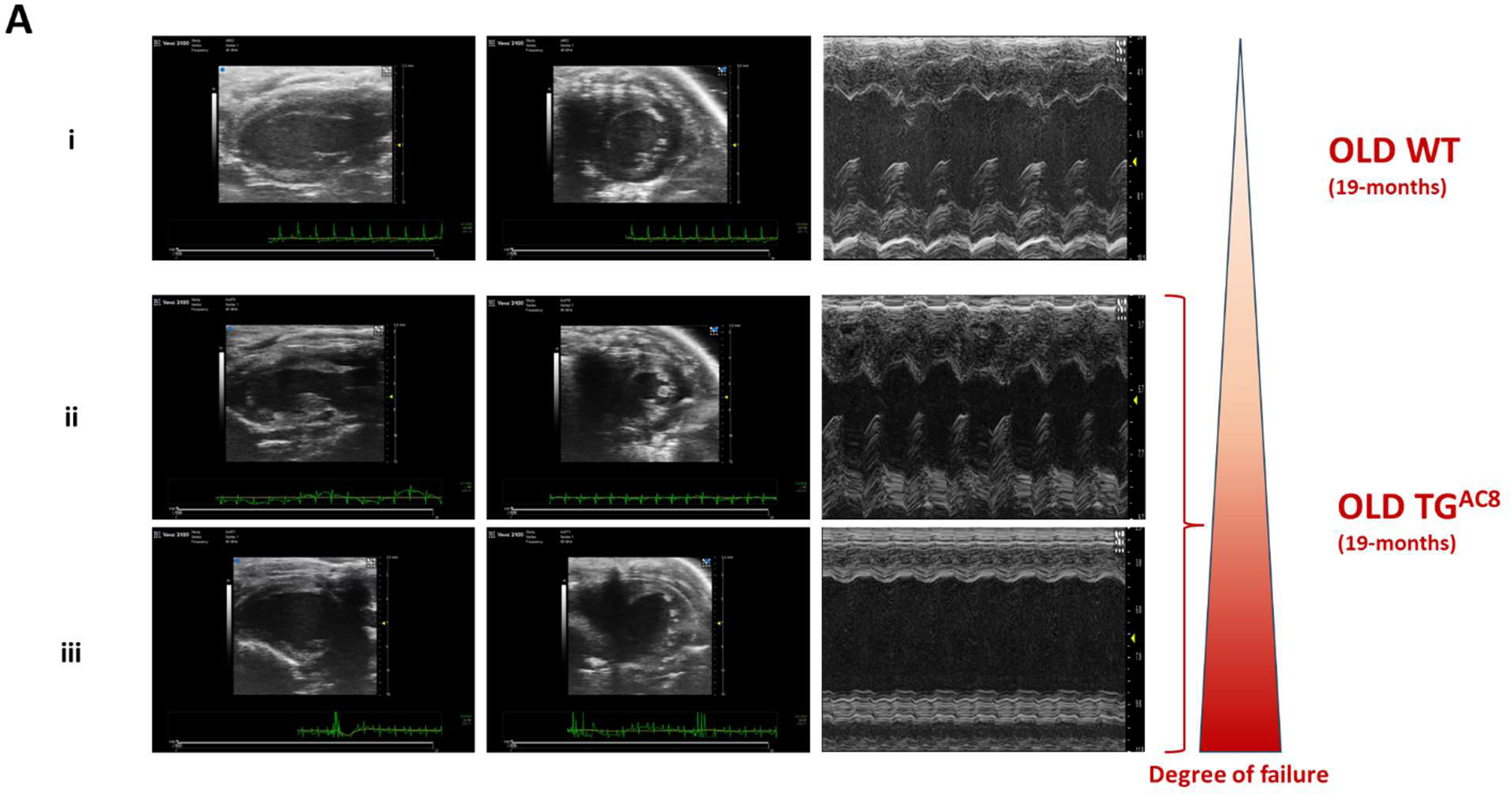
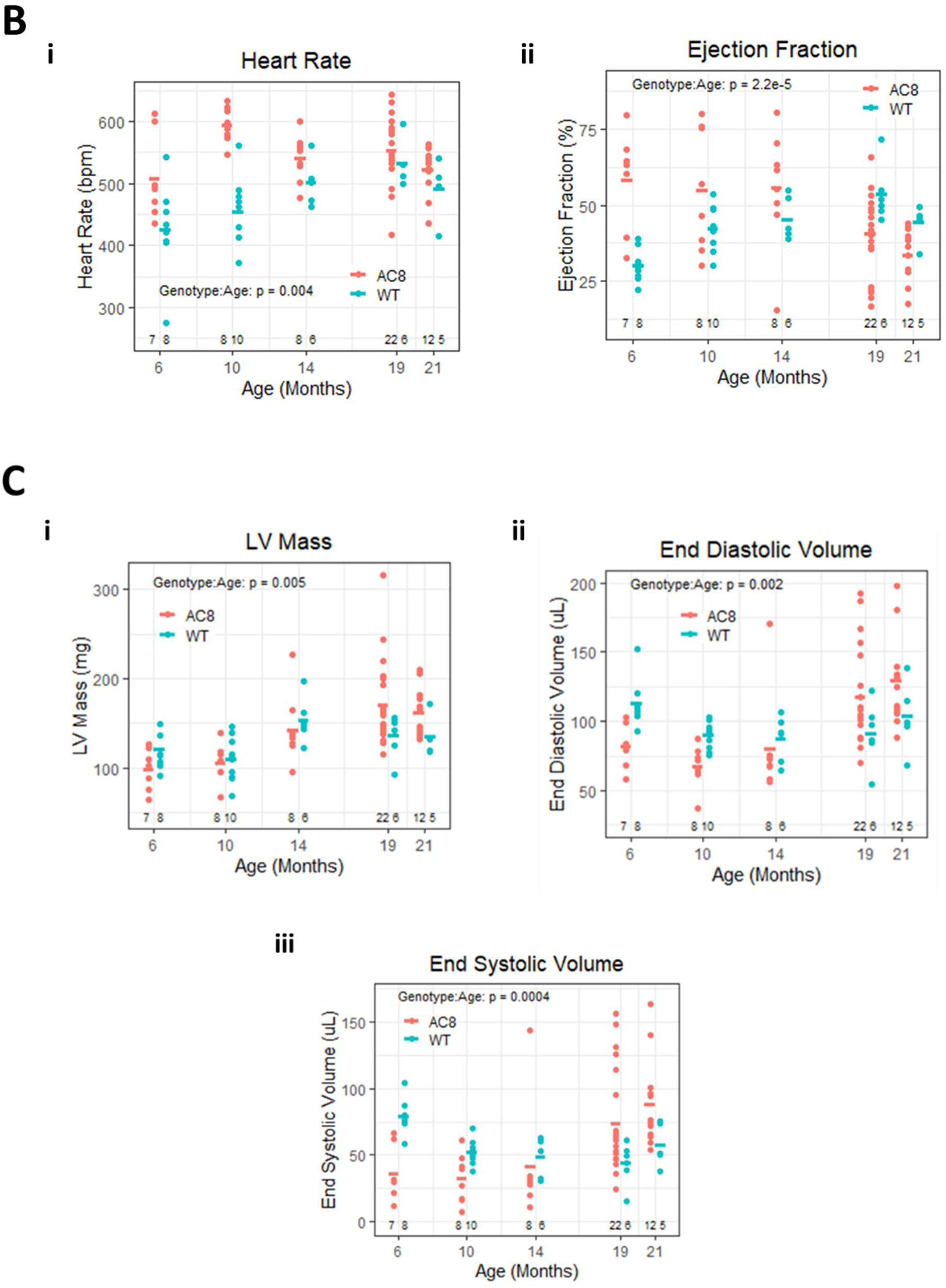
Heart damage by PQC insufficiency and mitochondrial dysfunction leads to heart failure contributing to TGAC8 accelerated cardiac aging. **A**), Representative images of echocardiograms in old TG^AC8^ and WT littermates (N=6-15 mice per group). Echocardiographic parameters **Bi**), heart rate (HR); **Bii**), ejection fraction (EF); **Ci**), left ventricular mass (LVM); **Cii**), end diastolic volume (EDV); **Ciii**), end systolic volume (ESV). Data are presented as mean along with all the data points. 2-way mixed ANOVAs was employed for the analyses. Statistical significance was assumed at p<0.05.

In conclusion, increased proteasome insufficiency and greater proteotoxic stress, in conjunction with a greater mitochondrial dysfunction and severe UPS and autophagy dysregulation in aged TGAC8, lead to excessive accumulation of enlarged LF bodies, brownish-to-black pigments and LC3+-/p62+-inclusions of heterogeneous size, and to the disruption of the cardiac contractile machinery by desmin disorganization and increased desmin PAOs. In addition, higher expression of HSP27S86 suggests increased cardiac myocyte damage in the old TGAC8 heart vs WT, and its co-localization with p62+-inclusions and LF aggregates points to these as mechanisms that underlie the upregulation of stress responses to preserve cardiomyocyte structure and function. Thus, increased failure of PQC mechanisms with aging leads to decreased heart function and highly contributes to TGAC8 accelerated cardiac aging.

## DISCUSSION

It is widely recognized that marked chronic catecholamine-induced cardiac stress leads to heart failure^72,73^. Our previous work in TG^AC8^ mice had shown that, up to 3-4 months of age, sustained upregulation of the AC/cAMP/PKA/Ca^2+^ axis^14^ activates numerous concentric adaptation mechanisms that protect heart health and function in TG^AC8^ mice. Among other mechanisms, protein quality control (PQC), including the ubiquitin proteasome system (UPS) and the autophagic machinery, became activated to ensure more efficient proteostasis in the TG^AC8^ heart. The present study evaluated proteostasis in the TG^AC8^ heart in more detail, focusing on key players involved in the autophagic process at 3-4 months of age in TG^AC8^, and probed whether PQC mechanisms remained adequate in advanced age.

### Autophagy in the TG^AC8^ heart at 3-4 months of age

Our results show that efficient modulation of LC3’s isotype-specific functions^19^ (by interacting with specific adapters for cargo recruitment^20^ and selection^74^) guarantees a <u>more favorable</u> autophagic process and effective clearance of damaged proteins, that essential PQC features in the context of a high cardiac workload and energy demands. One manifestation of upregulated autophagy in TG^AC8^ vs WT is that LC3, and specifically its isoforms LCA and LC3B, which localize to different phagophores/autophagosomes^19^, exhibit distinct expression patterns (**Figure 1A-B**). Specifically of the TG^AC8^ heart, at 3-4 months of age, LC3A orchestrates steady state autophagy, and *LC3A-mediated autophagy* modulates the protein quality control network^75^ to effectively resolve both aggresome formation and disposal of failing mitochondria by mitophagy^35^ (together with FKBP8). We speculate that LC3A *upregulation* is important for efficient PQC in the “stressed-out” TG^AC8^ heart, and that *downregulation* of LC3B is consistent with the idea that LC3A and LC3B are involved in different signaling pathways (UPR-activated response^75^ vs mitophagy^35,54^ vs apoptosis^76^ vs endosome-mediated autophagy^77^). LC3 post-translational modifications (PTMs) by phosphorylation, that affect its activity^21,22^, were also differentially modulated in the young TG^AC8^ vs WT heart in a way that <u>favored</u> both *early* and *late* stages of autophagy: specifically, phosphorylation of LC3 at S12 was significantly reduced, whereas its phosphorylation at T50 was significantly increased. Phosphorylation at S12 by PKA/PKC^78^ inhibits autophagy by preventing autophagosome wall elongation, whereas phosphorylation at T50 by STK3/4^22^ stimulates the autophagic process by promoting autophagolysosome fusion.

Another manifestation of fine-tuned, regulated efficient autophagy in TG^AC8^ vs WT was that p62/sequestosome-1, an essential autophagy adaptor protein^79^ that co-localizes with ubiquitin and LC3, was upregulated in TG^AC8^ vs WT at 3-4 months (**Figure 1Ci**). p62 plays a critical role in “aggresome” formation^80^ and clearance^49^ by activating autophagy while, itself, becoming an autophagy substrate^81^. p62 phospho-forms p62^S405^ and p62^S351^ (and their ratios to total p62) were also significantly upregulated in TG^AC8^ vs WT at 3-4 months (**Figure 1Cii-v**). p62^S405^ correlates to ubiquitinated cargo^25^ (including protein aggregates^24^), cellular inclusion bodies (the aggregates of aggregates^45^), and polyubitiquinated mitochondria^25^, whereas p62^S351^ has a higher affinity for the ubiquitin ligase adaptor Kelch-like ECH-associated protein 1 (KEAP1)^29^, resulting in constitutive activation of the transcription factor NF-E2-related factor 2 (NRF2) and regulation of its downstream signaling pathways^30^, with a positive feedback on p62 transcriptional activation itself^27^. Transient accumulation of p62 bodies by upregulation of p62 and its phospho-forms in the young TG^AC8^ heart, demonstrates its important role in the induction of autophagy^82^, and in both adaptation to the sequelae evoked by upregulation of signaling for protein aggregate clearance, and increased ROS scavenging, a protection mechanism from oxidative stress by sustained activation of the AC/cAMP/PKA/Ca^2+^ axis. ATG9A, which is the only integral membrane ATG protein, and is essential for autophagosome formation, was elevated in young TG^AC8^ LVs, vs WT (**Extended Data Figure S3**). ATG9A partially co-localizes with LC3^38^, and, as well as LC3, correlates with the number of autophagic bodies and autophagosomes^37^. In addition, it also participates in Parkin-mediated mitophagy via activation of the optineurin (OPTN)/ATG9A axis^83^, crucial for maintenance of healthy mitochondrial dynamics. LC3^+^-puncta were more numerous but smaller in TG^AC8^ at 3-4 months of age (**Figure 4Cv-vi**), and since cargo detection is regulated by *flexibility in autophagosome size*^37^, we interpret elevated levels of ATG9A together with higher counts of smaller LC3^+^-puncta, to indicate an increased *frequency of autophagosome formation* and enhanced *“selective”* autophagy and mitophagy in young TG^AC8^ vs WT.

Events-flux numbers were increased in TG**^AC8^** vs WT at 3-4 months, following CQ (**Figure 3C**), indicating accelerated flux. However, remarkably, LC3II did not accumulate, following CQ (**Figure 3Ai-ii**). Interestingly, protein levels of LC3**I** (both LC3AI and LC3BI) were *always decreased* in TG^AC8^, following CQ (**Figure 3Ai-ii**), whether LC3 was transcriptionally *upregulated* (LC3A) or *downregulated* (LC3B). We interpret this reduction to be the result of an *accelerated LC3 turnover* that prevents accumulation of LC3II at the lysosome. Specifically, increased protein levels of ATG4B (**Supplemental Information Table 2**), ATG16L1 (**Figure 3Bi, Supplemental Information Table 2**), and the trend towards accelerated flux of ATG4B in TG^AC8^ (**Figures 3Bii**), promote a *<u>faster</u> <u>LC3I to LC3II processing</u>*; additionally, increased protein levels of ATG9A (**Extended Data Figure S3**), of the lysosomal acid phosphatase ACP2 (**Figure 2C**), and of several cathepsins (**Figure 2A**)^14^, together with the upregulation of cathepsin L1 activity (**Figure 2B**), induce a *<u>faster degradation of LC3II</u>*. However, it might be argued that our conditions to assess the autophagic flux were suboptimal for our transgenic mouse, since the protocol was optimized to detect LC3II accumulation (following CQ) in WT controls. In fact, saturating concentrations of lysosomal inhibitors are needed, considering the well-known enhanced drug clearance of the heart compared to other organs^84^, together with the “high performance” of the TG^AC8^ heart^14^ and the increase in lysosomal (cathepsin L1) activity. Thus, we cannot exclude that the lack of accumulation of LC3II and p62 in TG^AC8^ LVs, following CQ treatment, is due to a faster clearance of CQ from the TG^AC8^ heart and/or because of the consequent reduced CQ concentration in TG^AC8^ LVs, the latter been shown to further increase lysosomal activity^85^, and autophagy flux in general. And if that were to the case, lysosomal inhibition would be incomplete in young TG^AC8^ LVs, leaving the existence of a remaining residual LC3-carrier flux in the TG^AC8^ heart, which further accelerates endogenous LC3 turnover that deprives the cellular pool of LC3I (lower levels of LC3AI and LC3BI) and prevents the accumulation of LC3II and p62 (no changes in LC3II and lower levels of p62), following CQ.

Nonetheless, *in toto*, the present results, together with those of our previous study^14^ showing increased proteasome activity without accumulation of insoluble aggregates, and increased levels of proteins involved in the autophagic machinery^14^, strongly suggest that UPS and autophagic pathways concurrently operate in a more efficient mode in the young TG^AC8^ heart to manage its state of chronic, marked cardiac cAMP-derived stress^49,86^.

### PQC mechanisms in aged TG^AC8^ and WT Autophagy

Autophagy markers LC3 and ATG4B became significantly downregulated with age in both TG^AC8^ and WT (**Figure 4A**, **Extended Data Figure S2**), and the number and the size of autophagosomes (LC3^+^-puncta) and lysosomes/late endosomes (LAMP1^+^-puncta), measured by fluorescence microscopy, dramatically increased (**Figure 4Cii-vi**). Downregulation of proteins involved in induction of autophagy, in conjunction with reduced numbers of *early* autophagy figures, and an increased number of *late* autophagic events (by TEM quantification and fluorescence microscopy) (**Figure 4B**), an indication of clearance impairment and of aggregates/inclusions accumulation, with age, demonstrate that cardiac aging, *per se*, negatively impacts on PQC.

Although the present results clearly point to less efficient autophagy due to aging, *per se*, several pieces of evidence suggest that some PQC mechanisms fail more severely than others in a context of chronic exposure (up to 21 to months) to marked cardiac cAMP-induced stress, and that autophagy is more severely compromised in TG^AC8^ than in WT. Firstly, LAMP1^+^-puncta *increased* in number whereas LC3^+^-puncta *decreased*, in old TG^AC8^, vs old WT (**Figure 4Cii, 4Cv**); and, both LAMP1^+^- and LC3^+^-puncta covered a smaller % of cell area in old TG^AC8^, vs old WT (**Figure 4Civ, 4Cvii**). Secondly, a reduction (by WB) of the nuclear to cytoplasm ratio of the master regulator protein of the CLEAR network, TFEB, in old TG^AC8^ vs old WT (**Figure 5C**), together with the transcriptional upregulation of several 14-3-3s isoforms in aged TG^AC8^ vs young TG^AC8^, and vs old WT(**Figure 5D**), demonstrate that the autophagic program is more severely suppressed in the aged TG^AC8^ than that due to “normal” aging in WT, and that lysosomal clearance more significantly impaired, manifested as the increased accumulation of insoluble LC3^+^-inclusions. Indeed, the fact that endogenous LAMP1^+^- and LC3^+^-puncta covered a smaller % of cell area in TG^AC8^ vs WT of advanced age, and were highly heterogeneous in size (**Figure 4D**), portrays a scenario compatible with the presence of increased “aggresomes”^45,75^ (the “aggregate of aggregates” encaged by intermediate filament proteins^60^) resulting from a buildup of numerous small pre-aggresomal bodies which are, at younger age, compartmentalized as a cytoprotective mechanism^87,88^.

<u>LC3A</u> was transcriptionally *upregulated*^14^ and its carrier-flux *accelerated* in old TG^AC8^, vs WT (**Figure 5Ai**); however, *overall* autophagic flux was reduced more in TG^AC8^ than in WT, as demonstrated by the reduced levels of <u>p62</u> (**Figure 5Aiii**), <u>ATG16L1, ATG4B</u> and <u>ALIX</u> (**Figure 5Bi-iii**), following CQ treatment. Inhibition of proteasome activity has been linked to activation of LC3A specifically, as a general stress response^75^, without involvement of LC3B; additionally, LC3A silencing (by promoter methylation), primes cells for aggresome formation to achieve cellular homeostasis, in conditions of altered autophagy, such as in cancer cells^89^. Thus, we correlate *increased* LC3A levels and the *accelerated LC3A-flux* to the presence of increased protein aggregates in aged TG^AC8^ vs old WT. Indeed, accelerated flux has been shown to be a compensation mechanism that can be maladaptive in disease conditions that contribute to the remodeling of the myocardium under stress^90^. Thus, increased LC3A-flux in aged TG^AC8^ vs WT is another index of an indication of initiated signaling of oxidized lipoprotein accumulation within the context of chronic upregulation of the AC/cAMP/PKA/Ca^2+^ axis in old TG^AC8^.

Damaged and/or dysfunctional mitochondria^8,49^ are unable to support high energy demands, and overall mitochondrial fitness declines, with deterioration of heart function leading to disease conditions. Significant alterations in mitochondrial morphology, reflected by mitochondrial dysfunction and increased ROS production, in response to LC3A-induced autophagy, via the PERK-eIF2a-ATF4 axis of the UPR pathway, have been shown to result in altered mitochondrial dynamics and stress induced senescence^75^. LC3A activation was greater, and its flux accelerated. In aged TG^AC8^, it was accompanied by mitochondrial dysfunction, also increased, vs WT, suggesting that greater mitochondrial dysfunction may contribute to increased LC3A expression in old TG^AC8^ LVs. The higher cellular stress-response and the need of a “*specialized autophagy*” to maintain cellular/mitochondrial homeostasis, further emphasizes the intricate interplay between autophagy and mitochondrial dynamics^75^.

Although in aged TG^AC8^, both L-OPA1 and S-OPA1 protein levels were significantly decreased vs WT (by WB) (**Figure 7Ci-ii**), the ratio of S-OPA1 to L-OPA1 was significantly increased (**Figure 7Ciii**). In contrast, protein levels of HSP60 (WB) were reduced, in TG^AC8^ vs WT (**Figure 7D**). These results point to a reduction in mitochondrial fusion in aged TG^AC8^ vs WT: excessive stress-induced processing of OPA1 by OMA1^60^ triggers mitochondrial fragmentation, accelerating mitochondrial fission^69^. Because myocardial function depends on balanced mitochondrial fusion and fission^91^, we conclude that although mitophagy signaling was upregulated through a variety of pathways^53^ (**Figure 7E**), such altered mitochondrial dynamics and reduced protection by HSP60 contribute to mitochondrial dysfunction and failing of mitophagy to a greater extent, in aged TG^AC8^ vs WT, resulting in poor mitochondrial fitness/function, and is further aggravated by <u>insufficient</u> lysosomal activity.

Accumulation of LC3^+^-bodies (**Figure 4C**) and p62^+^-inclusions (**Figure 8H**), like LF aggregates (**Figure 8A-B**) in late stages of autophagy that cannot be removed from the cytosol by the UPS^92^ because they are covalently cross-linked aggregates, is an indication of a greater lysosomal impairment^93^ in aged TG^AC8^ vs WT. In addition, the black-to blueish pigment aggregates (**Figure 8C**), and LC3^+^-cardiac cells (**Figure 8D-E**), in older TG^AC8^, but not in age-matched WT, suggests changes in cargo processing, and accumulation of different material in TG^AC8^ LV vs WT. Because aggregates/inclusions are a manifestation of “normal” aging, we interpret more marked changes in TG^AC8^ than in WT to be an induction of accelerated aging in TG^AC8^.

### Proteasome

Although proteasome activity did not differ between genotypes at 18 months, between 3-4 months and 18 months, proteasome function progressively declined in TG^AC8^ but not in WT (**Figure 6A**). In addition to failure at the lysosome, variations in UPS activity and expression, reflecting age-associated metabolic, functional, or simply structural changes, occur in response to the different demands for the turnover of signaling and structural proteins. Specifically, cardiac proteasomal insufficiency has been linked to accumulation of “aggresomes”, leading to partial disorganization of protein filaments^60,61^, that are displaced from their normal cellular distribution^45^. In line with these results, accumulation of insoluble misfolded proteins was increased in aged TG^AC8^ vs WT, whereas soluble aggregates (soluble proteins undergoing normal turnover and misfolded proteins *en route* to degradation by the UPS), became significantly reduced in *both* genotypes at 18 months (**Figure 6B**). We interpret this result to indicate that protein insolubility increases with age, *per se*, and that, with aging, misfolded proteins are more “aggregate-prone”^45^. Thus, the presence of high molecular weight oligomers^45^ in TG^AC8^, more stable than the intermediate conformers from which they are derived, clearly demonstrate that the proteasome of the old TG^AC8^ heart is “out-won” in the competition for misfolded substrates prior to their aggregation^94^.

Although protein translation rates at older ages did not differ between genotypes, protein synthesis decreased significantly more with aging in TG^AC8^ heart (**Figure 6D**) in accordance with a “normal” aging process^48,95^. We interpret this reduction to indicate that, exactly as in the age-matched WT, the old TG^AC8^ heart reduces energy use and load on the protein quality control machinery, by directing proteases and chaperons activity (**Figure 6C**) towards repair of existing proteins rather than the folding of new ones^48,95,96^. Thus, the old TG^AC8^ heart utilizes a mechanism that preserves and increases life span as in healthy aging^96^, to optimize energy use by reducing the formation of *de-novo* proteins, in order to focus on repairing the existing proteome.

Disorganization of desmin filaments that interlink the contractile myofibrillar apparatus with mitochondria, nuclei and the sarcolemma, has been shown to affect mitochondrial positioning, therefore compromising mitochondrial function, and leading to cardiomyocyte dysfunction^67,97^. Negative modulation of desmin’s physical properties and assembly by its PTMs leads to the build-up of cardiac preamyloid oligomers (PAOs) and has been recently indicated to be an index of premature and acquired heart failure^4,6^. Desmin tagging for ubiquitination and cleavage, and desmin PAOs accumulation (assessed by protein fractionation and WB) was also highly increased in old vs WT TG^AC8^ hearts (**Figures 8F-G**).

The number of cardiomyocytes expressing HSP27^S^^86^ (assessed by fluorescence microscopy) was also elevated in aged TG^AC8^ vs WT, and this increased expression was co-localized with p62^+^-inclusions (**Figures 8H**). HSP27^S86^ co-localizes with contractile proteins (cardiac troponin T) at sarcomeres^71,98^ and it is upregulated in stress conditions (including cardiac injury^99^) where it enhances heart tolerance to stress^100^ while maintaining myocardial function^71,98^. Co-localization of HSP27^S86^ with p62^+^-inclusions in the aged TG^AC8^ suggests increased heart damage. Indeed, myocardial function and cardiac structure progressively declined from 6 months to 21 months of age, leading to a failing hypertrophic and dilated heart at 19 months of age (**Figures 9A-C**).

In conclusion, in response to chronic cardiac AC-dependent cAMP-stress that activates protective concentric signaling circuitry, including autophagy/autophagic flux and proteasome activity, and maintains PQC that preserve heart health early in life^14^, with age, <u>proteasomal insufficiency</u> and <u>compromised PQC</u>, in a context of a <u>dysregulated</u> <u>autophagy flux</u> in the older TG^AC8^, indicates the overall presence of maladaptive responses with <u>ineffective cargo clearance</u> and <u>mitochondrial dysfunction</u>. This scenario leads to severe loss of proteostasis in the old TG^AC8^ heart vs WT, contributing to increased cardiac damage, accelerated cardiac aging (cardiomyopathy^15^, desminopathy^90^) and reduced cardiac health-span.

## METHODS

### Experimental animals

All studies were performed in accordance with the Guide for the Care and Use of Laboratory Animals published by the National Institutes of Health (NIH Publication no. 85-23, revised 1996). All experimental protocols were approved by the Animal Care and Use Committee of the National Institutes of Health (protocol #441-LCS-2025). Breeder pairs of mice in which human TG^AC8^ was over-expressed under the murine α-myosin heavy chain (α-MHC) promoter (TG^AC8^), and wild-type littermates (WT) (background strain C57BL/6 from Jackson Labs, Stock # 000664), were a gift from Nicole Defer/Jacques Hanoune, Unite de Recherches, INSERM U-99, Hôpital Henri Mondor, F-94010 Créteil, France^15^. HAC8^+^ mice were crossed with C57Bl6 mice, and housed in a climate-controlled room with 12-hour light cycle and free access to food and water, as previously described^72^. All assays were performed in 3-4-month-old (young) and 17-21-month-old (old) males and compared to age-matched wild-type littermates (WT). Mice were sacrificed using Ketamine/Xylazine mixture, by intraperitoneal (IP) administration, in accordance with the approved protocol (#441-LCS-2025).

### Heart and cardiac tissue isolation

The heart was quickly removed and placed into cold PBS solution. The left ventricle (LV) free wall, without the septum, was identified anatomically under a dissecting microscope, excised, snap-frozen in liquid nitrogen and stored at −80°C for further analyses.

### Protein extraction and WB

Flash frozen tissue was homogenized as previously described^14^. Briefly, snap-frozen LVs were homogenized and lysed in ice cold RIPA buffer (Thermo Fisher Scientific: 25 mM Tris-HCl (pH 7.6), 150 mM NaCl, 1% NP-40, 1% sodium deoxycholate, 0.1% SDS) supplemented with a Halt protease inhibitor cocktail (Thermo Fisher Scientific), Halt phosphatase inhibitor cocktail (Thermo Fisher Scientific) and 1 mM phenylmethyl sulfonyl fluoride, using a Precellys homogenizer (Bertin Instruments) with a tissue homogenization kit CKMix (Bertin Instruments) at 4°C. Lysates were then centrifuged at 10,000g for 10 min at 4°C to separate insoluble material. Nuclear and cytosolic lysates were prepared from snap-frozen LV using a subcellular protein fractionation kit for tissues (Thermo Fisher Scientific) as per the manufacturer’s instructions. The protein concentration of samples was determined using the Bicinchoninic Acid (BCA) Assay (Thermo Fisher Scientific). Samples were denatured in 4X Laemmli sample buffer (BioRad Laboratories) containing 355 mM 2-mercaptoethanol at 95°C for 5 min, or DTT at 70°C for 10 min, and proteins (25μg/lane) were resolved on 4-20% Criterion TGX Stain Free gels (Bio-Rad Laboratories) by SDS/PAGE. For detection of HSP60, samples were denatured in XT sample buffer containing 1X XT reducing reagent (BioRad Laboratories) at 95°C for 5 min, and 25μg protein resolved on a 12% Criterion™ XT Bis-Tris gel using XT MES running buffer (BioRad Laboratories). For Stain-Free™ gels, to induce crosslinking of trihalo compound with protein tryptophan residues, gels were exposed to UV transillumination for 2-3 mins. Proteins were then transferred to low fluorescence polyvinylidene difluoride (LF-PVDF) membranes (BioRad Laboratories), using an electrophoretic transfer cell (Mini Trans-Blot, Bio-Rad). Membrane total protein was visualized using an Amersham Imager 600 (AI600) (GE Healthcare Life Sciences) with UV transillumination to induce and capture a fluorescence signal. Membranes were blocked with either 5% milk/tris-buffered saline with Tween-20 (TBST) or EveryBlot Blocking Buffer (BioRad Laboratories) as appropriate, and were then incubated with the primary antibodies indicated in the **Supplemental Information Table 1** Primary antibodies were then detected using horseradish peroxidase (HRP) conjugated antibody (Invitrogen) at 1:10,000, and bands were visualized using Pierce SuperSignal West Pico Plus ECL substrate kits (Thermo Scientific), the signal captured using an Amersham Imager 600 (AI600) (GE Healthcare Life Sciences). Total protein for HSP60 blot was visualized by staining membrane with 1X Amido Black solution (Millipore Sigma) according to manufacturer’s protocol. Band density was quantified using ImageQuant TL software (GE Healthcare Life Sciences), normalized to total protein, and TG^AC8^ to average of WT.

### Immunostaining (IHC)

Endogenous macroautophagic puncta in young and aged TG^AC8^ and aged-matched WT were determined in 5 µm thick paraffin sagittal sections of cardiac tissue. Following standard procedures, deparaffinization and rehydration were followed by heat antigen retrieval in a citric acid base solution (H-3300, VectorLabs). Autofluorescence was quenched with a 10 min incubation with 1mg/ml Sodium Borohydride solution in PBS. Cardiac sections were blocked in 10% goat serum for 30 min, and incubated at 4°C overnight with the following primary antibodies: rabbit monoclonal LC3A/B (1:50, cloneD3U4C, #12741 Cell Signaling), mouse monoclonal p62 (1:50, clone 2C11, #ab56416; Abcam), rat monoclonal LAMP-1 (1:100, clone 1D4B, #1D4B; DSHB), rabbit monoclonal p(S82)-HSP27 (1:100, clone D1H2F6, #9709; Cell Signaling), and Wheat Germ Agglutinin (WGA) (1:100, #W21404; ThermoFisher Scientific). After incubation with corresponding conjugated secondaries and counterstaining with 500 nM 4′,6-diamidino-2-phenylindole (DAPI) for about 1 hr at RT, immunolabeled samples were mounted using an antifade gel mounting medium (Vectashield Vibrance; Vectorlabs) and examined with a Zeiss LSM 900 confocal microscope. At least 10 random fields of optical regions (100 µm2) of 5 µm thickness per cardiac section (n=3) were collected with a Plan-Apochromat 63x/1.40Oil DIC M27 objective and projected on a single extended projection image to analyze. Puncta counting and areas of vesicles were measured using manual tracking with ImageJ software by a blinded investigator.

### Transmission Electron Microscopy (TEM)

Samples for transmission electron microscopy were fixed using 2.5% glutaraldehyde in 0.1 M sodium cacodylate buffer, pH 7. Then, they were post-fixed in 1% osmium tetroxide for 1 hr at 4°C in the same buffer, dehydrated, and then embedded in Embed 812 resin (Electron Microscopy Sciences, Hatfield, PA) through a series of resin resin-propylene oxide gradients to pure resin. Blocks were formed in fresh resin contained in silicon molds, and the resin was polymerized for 24-48 hrs at 65°C; blocks were then trimmed and sectioned in an EM UC7 ultramicrotome (Leica Microsystems, Buffalo Grove, IL) to obtain both semi-thick (0.5-1 µm width) and ultrathin (40-60 nm width) sections. Semi-thick sections were mounted on glass slides and stained with 1% toluidine blue in a 1% borax aqueous solution for 2 min. Palade-stained (OsO4) and Toluidine blue-stained semi-thick sections were then imaged using a Leica AXIO Imager light microscope with a 63x oil immersion objective, for quality control and Lipofuscin (LF) quantification. Ultrathin sections were stained with uranyl acetate and lead citrate and then imaged on a Talos L120C Transmission Electron Microscope (TEM) with a 4K Ceta CMOS camera, for macroautophagic events quantification. Micrographs at ×2000 ×5000 and ×11000 magnification were obtained from randomly selected cytoplasmic areas of cardiomyocytes, for illustration purposes and quantitative analysis of macroautophagic events population. Two stereological parameters were determined: (a) the numerical profile density Na (number of figures of interest / μm^2^ of cell fraction), and (b) volume density of figures of interest (Vv; i.e., the volume fraction of cardiomyocyte cytoplasm occupied by figures of interest). Volume density was obtained following a point analysis using a simple square lattice test system^101^, with superposition of a virtual grid over the micrographs where the user performs a point-counting method. Autophagosomes (also referred in the text as early events) were identified as double-membrane vesicles with identifiable cargo and comparable density to the surrounding cytosol. Autolysosomes (late events) and residual bodies were denoted as single membrane vesicles containing non-identifiable cargo of higher density than the surrounded cytosol and fragmented organelles. Lipofuscin bodies were identified following Palade staining^102^ by the presence of densely packed lipids with a dark brown-black color against cardiomyocyte cytosol, which allowed a straightforward segmentation of the image and posterior quantification of the planimetric and stereological parameters. Point counting and areas of vesicles and cytosol were measured using manual tracking with ImageJ software by a blinded investigator.

### RT-qPCR

RT-qPCR of LV tissue was performed to determine the transcript abundance of 14-3-3 subunits. RNA was extracted from left ventricular myocytes (VM) with RNeasy Mini Kit (Qiagen, Valencia, CA) and DNAse on column digestion. 1 µg of total RNA was used for cDNA synthesis with MMLV reverse transcriptase (Promega). RT-qPCR was performed using a QuantStudio 6 Flex Real-Time PCR System (Thermo Fisher Scientific) with a 384-well platform. The reaction was performed with a FastStart Universal SYBR Green Master Kit with Rox (Roche) using the manufacturer’s recommended conditions; the sizes of amplicons were verified. Each well contained 0.5 μl of cDNA solution and 10 μl of reaction mixture. Each sample was quadruplicated and repeated twice using de novo synthesized cDNA sets. Preliminary reactions were performed to determine the efficiency of amplification. RT-qPCR analysis was performed using the ddCt method. Primers were selected with Primer Express 3.0 software (Applied Biosystems). The primers sequence is indicated in the **Supplemental Information Table 2**.

### Cathepsin L1 activity assay

Cathepsin L1 activity was measured using the fluorimetric kit (Abcam, ab65306) according to the manufacturer’s instructions. Briefly, snap-frozen LVs stored at −80°C were thawed on ice, and 10 mg of tissue was resuspended in 50 µl of CL Buffer, then homogenized with a Dounce homogenizer sitting on ice, with 10–15 passes. Samples were centrifuged for 2–5 mins at 4°C at top speed using a cold microcentrifuge to remove any insoluble material, then supernatants were collected and transferred to a clean tube, on ice. Activity was performed at different dilutions to reach Vmax, in the presence of Ac-FR-AFC fluorogenic substrate (200 µM final concentration), at 37°C for 1–2 hours, at an excitation wavelength of 400 nm and an emission wavelength of 505 nm using the Cytation 5 microplate reader (Biotek). Fold-increase in Cathepsin L activity was determined by comparing the relative fluorescence units (RFU) with the level of the uninduced control, after subtraction of the negative control sample. TG^AC8^ values were normalized to average of WT.

### Proteasome activity assay

Proteasome activity was measured with Proteasome-Glo™ Chymotrypsin-Like Assay G8621, accordingly to the manufacturer’s instructions. Briefly, lysed tissue was diluted with the provided buffer and 50 μg of the sample was loaded on a 96 well microplate. The resulting “glow-type” luminescent signal was then quantified utilizing the Spectramax microplate reader (San Jose), with luminescence recorded at 500. Human 20S Proteosome was used as a positive control.

### Protein aggregation assay

The isolation of protein aggregates and the assessment of protein aggregation was conducted using the Proteostat kit (Enzo Life Science, ENZ-51023), as previously published^14^. Specifically, 50 μg of proteins were loaded into a 96-well plate for analysis. Fluorescence measurements were obtained with an excitation wavelength of 550 nm and emission wavelength of 600 nm, employing a time interval of 15 minutes between readings, and background readings were subtracted from sample recordings.

### Protein synthesis assessment

Protein synthesis was assessed by SUnSET-Western Blot as previously described^14,103^. Briefly, the puromycin solution was prepared in PBS, sterilized by filtration, and a volume of 200 μl was injected in TG^AC8^ mice and age-matched WT controls intraperitoneally, to achieve a final concentration of 0.04 μmol/g of body mass. After 30 min, mice were sacrificed, the LV was harvested and snap frozen in liquid nitrogen. Protein extraction was performed using Precellys, quantified with BCA assay 25 μg of total protein were separated by SDS-PAGE; proteins were then transferred onto PVDF membrane and incubated overnight in the anti-puromycin primary antibody (MABE343, Sigma-Aldrich, St. Louis, MO). Visualization of puromycin-labeled bands was obtained using horseradish peroxidase conjugated anti-mouse IgG Fc 2 a secondary antibody (Jackson ImmunoResearch Laboratories Inc, West Grove, PA, USA), using Pierce Super Signal ECL substrate kit (Pierce/ Thermo Scientific Rockford, IL). Chemiluminescence was captured with the Imager AI600 and densitometry analysis was performed using ImageQuantTL software (both by GE, Boston, MA). Total protein was used as control for protein loading. Genotypic differences of protein synthesis were tested via an as unpaired t-test.

### Quantification of desmin preamyloid oligomers (PAOs)

Proteins were extracted according to the method described in *Li Z et al* (https://doi.org/10.1101/2023.05.09.540017). Myofilament-enriched, insoluble fractions were prepared as described previously^6^. Fractions were separated using precast NuPAGE gels (Thermo Fisher Scientific) with MES buffer, and probed with the desmin antibody (DE-U-10, Sigma, 1:10,000) or the A11 antibody (kindly provided by Dr. Charles Glabe at UC Irvine^104^, 1:1,500). TG^AC8^ values were normalized to WT.

### Echocardiography

Mice underwent echocardiographic (Echo) examination (40-MHz transducer; Visual Sonics 3100; Fuji Film Inc, Seattle, WA) under light anesthesia with isoflurane (2% in oxygen) via nose cone; temperature was maintained at 37°C using a heating pad. Mice were placed in the supine position; skin hair in the chest area was shaved. Standard electrocardiogram (ECG) electrodes were placed on the limbs and ECG Lead II was recorded simultaneously with acquisition of echo images. Each Echo examination was completed within 10 min. Parasternal long-axis views of the LV were obtained and recorded to ensure that mitral and aortic valves and the LV apex were visualized. From the parasternal long-axis view of the LV, M-mode tracings of LV were obtained at mid-papillary muscle level. Parasternal short-axis views of the LV were recorded at the mid-papillary muscle level. Endocardial area tracings, using the leading-edge method, were performed in the 2D mode (short-axis and long-axis views) from digital images captured on a cine loop to calculate the end-diastolic and end-systolic LV areas. LV End-Diastolic Volume (EDV) and End-Systolic Volume (ESV) were calculated by a Hemisphere Cylinder Model method. Ejection Fraction (EF) was derived as EF = 100*(EDV - ESV)/EDV. LV mass (LVM) was calculated from EDV, septal thickness (IVS) and LV Posterior Wall (PW). All measurements were made by a single observer who was blinded to the identity of the tracings. All measurements were reported an average of five consecutive cardiac cycles covering at least one respiration cycle (100 times/min in average). The reproducibility of measurements was assessed by repeated measurement a week apart in randomly selected images; the repeated measure variability was less than 5%.

### Sample extraction and proteoOMICs analysis

LV proteome analysis was done accordingly to our previous work^14^. Briefly, four LV samples from WT and TG^AC8^ mouse hearts were snap frozen in liquid nitrogen and stored at −80°C. On average, 2 mg of muscle tissue from each sample was pulverized in liquid nitrogen and mixed with a lysis buffer containing (4% SDS, 1% Triton X-114, 50 mM Tris, 150 mM NaCl, protease inhibitor cocktail (Sigma)), pH 7.6. Samples were sonicated on ice using a tip sonicator for 1 min with 3s pulses and 15s rest periods at 40% power. Lysates were centrifuged at +4°C for 15 min at 14,000 rpm, aliquoted and stored at −80°C until further processing. Protein concentration was determined using the commercially available 2-D quant kit (GE Healthcare Life Sciences). Sample quality was confirmed using NuPAGE protein gels stained with fluorescent SyproRuby protein stain (ThermoFisher). In order to remove detergents and lipids, 500 µg of muscle tissue lysate was precipitated using a methanol/chloroform extraction protocol (sample:methanol:chloroform:water 1:4:1:3) (Wessel and Flugge^105^). Proteins were resuspended in 50 µl of concentrated urea buffer (8 M Urea, 150 mM NaCl [Sigma]), reduced with 50 mM DTT for 1 hour at 36°C and alkylated with 100 mM iodoacetamide for 1 hr at 36°C in the dark. The concentrated urea/protein mixture was diluted 12 times with 50 mM ammonium bicarbonate buffer, and proteins were digested for 18 hr at 36°C, using trypsin/LysC mixture (Promega) in 1:50 (w/w) enzyme to protein ratio. Protein digests were desalted on 10×4.0 mm C18 cartridge (Restek, cat# 917450210) using Agilent 1260 Bio-inert HPLC system with a fraction collector. Purified peptides were speed vacuum dried and stored at −80°C until further processing. A subset of 8 muscle samples (100 µg) each corresponding to 4 controls and 4 TG^AC8^ LVs and one averaged reference sample were labeled with 10-plex tandem mass spectrometry tags (TMT) using standard TMT labeling protocol (Thermo Fisher). 200 femtomole (fM) of bacterial beta-galactosidase digest (SCIEX) was spiked into each sample prior to TMT labeling to control for labeling efficiency and overall instrument performance. Labeled peptides from 10 different TMT channels were combined into one experiment and fractionated.

### Statistics

Data are presented as median ±range or median ±interquartile range, except for the echocardiography, expressed as mean along with all the data points. Representative TEM and IHC images, and western blots were chosen based on the resemblance to the average values. The number of samples per group are indicated in each legend. Statistical analyses were performed in GraphPad Prism 10 and with R. ROUT analyses were applied to identify outliers, which were excluded from group comparison analyses. Student *t* test with Welch’s correction was used to compare data between 2 groups. Two-way ANOVA followed by Original FDR method of Benjamini and Hochberg was used for multiple comparisons^106^. For datasets with matching LC3I and LC3II from the same animal, a repeated-measures approach with three-way mixed ANOVA or mixed-effect analysis for repeated-measures was taken, followed by Original FDR method of Benjamini and Hochberg. Statistical significance was assumed at p<0.05.

## ACKNOWLEDGMENTS

The study was designed by M.G.P. and E.G.L. The experiments were conducted by M.G.P., M.C.B., D.R.R., G.A., A.P., I.A., K.C., Y.S.T, and H.K.. The data were analyzed by M.G.P., M.C.B., D.R.R, G.A., C.H.M., I.A., K.V.T, and J.Q. The article was written by M.G.P., M.C.B. and E.G.L., and approved by all authors.

## EXTENDED DATA FIGURE LEGENDS

**Extended Data Figure 2 - Table.**
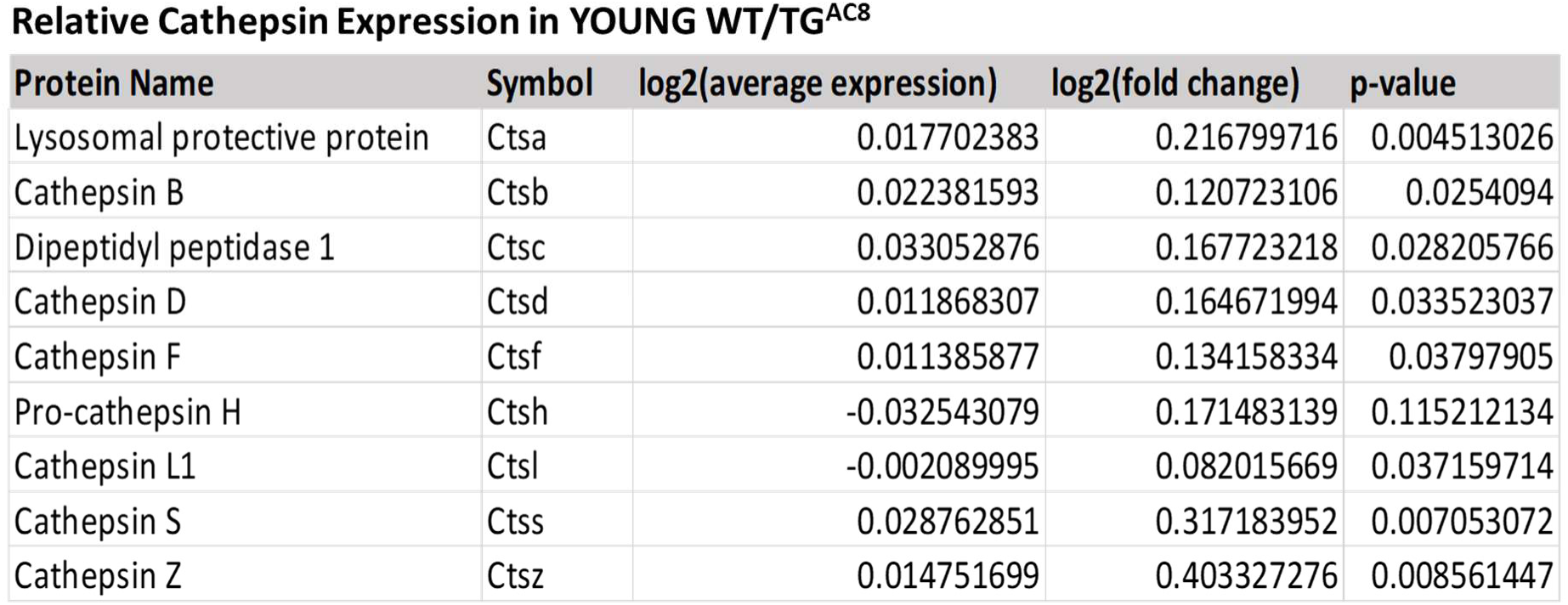
Relative Cathepsin Expression in TG^AC8^ and WT littermates at 3-4 months of age. MA plot of relative cathepsin expression (proteOMICs) in TG^AC8^ LVs at 3-4 months of age.

**Extended Data Figure 3 - Figure S1.**
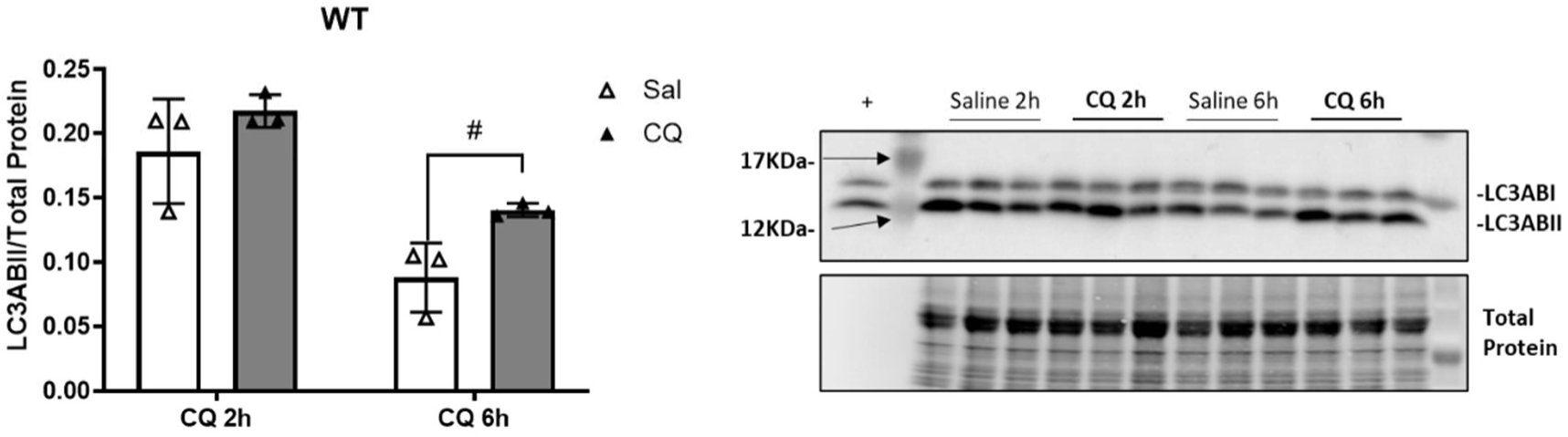
Chloroquine (CQ) time-course in WT at 3-4 months of age. Bar graphs and Western blot analysis of LC3II in WT (3-4 months of age) treated by intraperitoneal administration (IP) of chloroquine (CQ) (50 mg/kg) or saline. LVs were collected 2 hours and 6 hours after drug administration and snap-frozen (n=3 mice per group). A two-way ANOVA, followed by Original FDR method of Benjamini and Hochberg post-hoc multi-comparison test was used. Data are presented as the median ±interquartile range. **^#^** indicates significant (p<0.05) differences between treatments (CQ vs saline).

**Extended Data Figure 4 - Figure S2.**
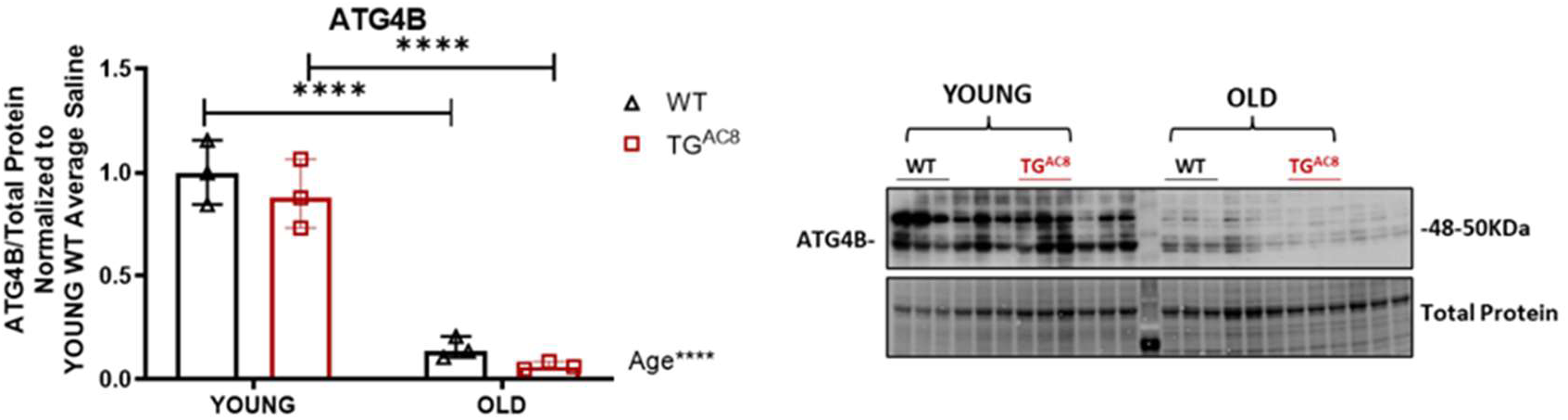
ATG4B is reduced with age in TG^AC8^ and WT littermates. Bar graphs and Western blot analysis of ATG4B in TG^AC8^ and WT at 3-4 months (n=3-5 mice per group) and 21 months of age (n=5 mice per group). A two-way ANOVA, followed by Original FDR method of Benjamini and Hochberg post-hoc multi-comparison test was used. Data are presented as the median ±range. Symbols **\*** indicate significant (p<0.05) differences between ages (young vs old).

**Extended Data Figure 5 - Figure S3.**
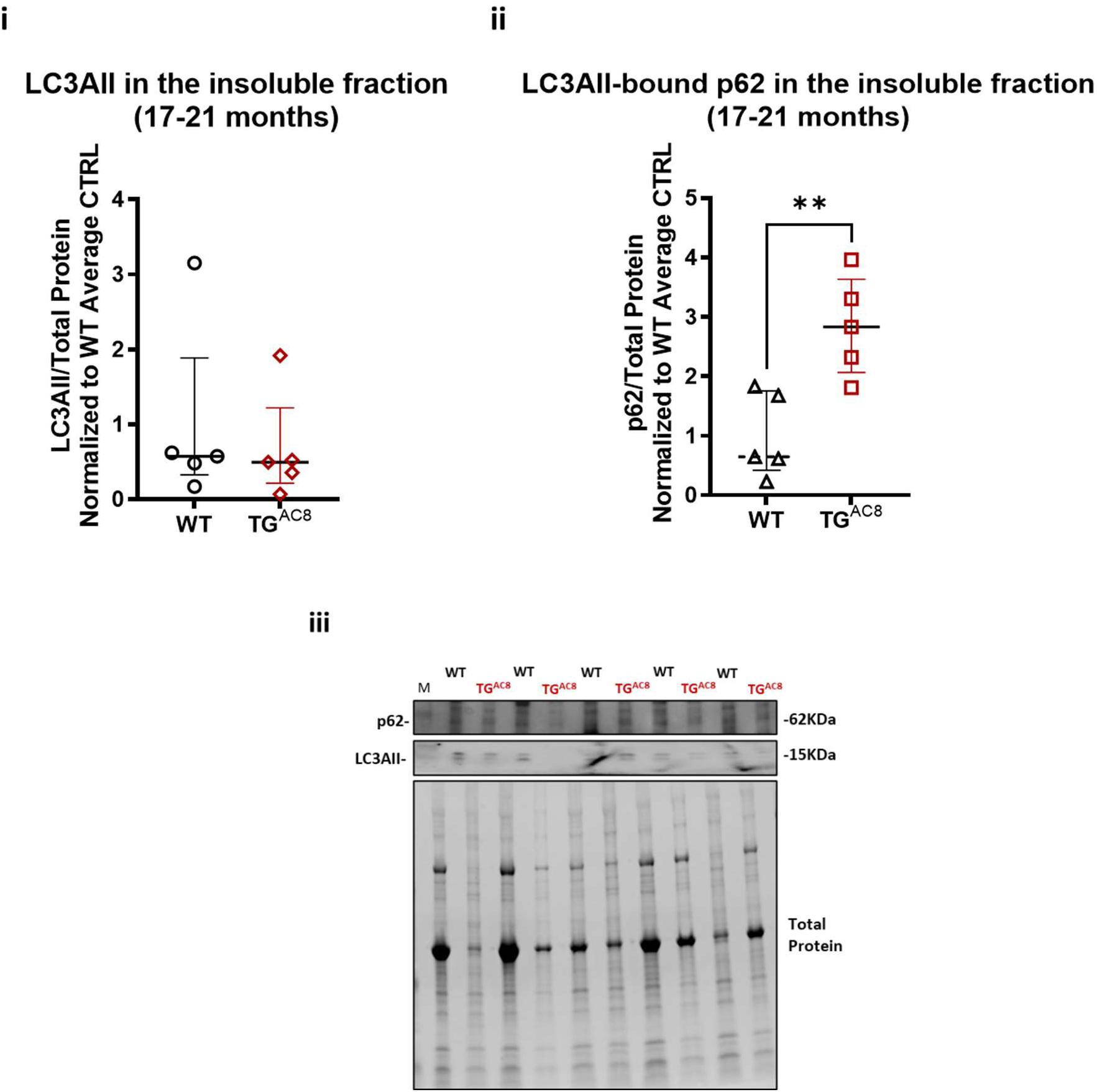
Assessment of LC3AII and *LC3AII-bound* p62 in insoluble fractions (pellets) in aged TG^AC8^ and WT littermates (17-21 months of age) Pellets from total protein lysates (RIPA) were lysed with WB loading buffer and protein resolved with SDS-PAGE. Proteins were then transferred onto PVDF and probed with LC3A and p62 antibodies, as indicated in the WB methods section. N=5 mice per group, TG^AC8^ and WT littermates (21 months of age). Unpaired 2-tailed Student t test with Welch’s correction was used. Data are presented as the median ±interquartile range. Symbols * indicate significant (p<0.05) differences between genotypes (WT vs TG^AC8^). **i**), LC3AII; **ii**), LC3AII-bound p62; **iii**), Paired WB blot image.

**Extended Data Figure 5 - Figure S4.**
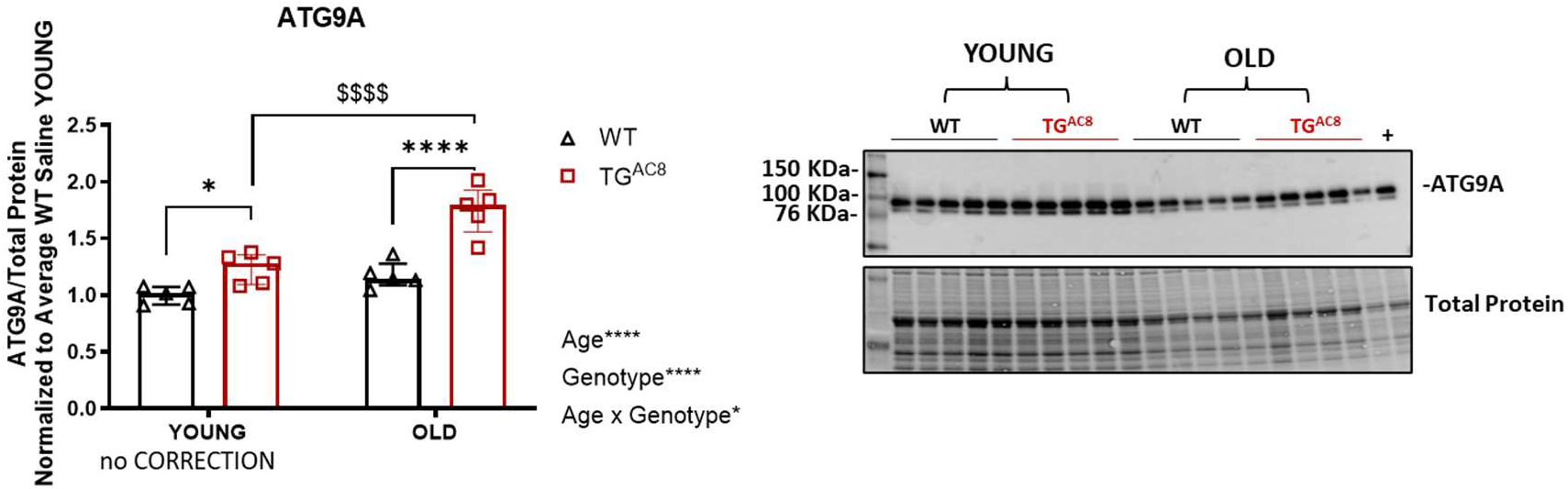
ATG9A is increased in old TG^AC8^ heart. Bar graphs and Western blot analysis of ATG9A in TG^AC8^ and WT at 3-4 months and 21 months of age (n=5 mice per group). A two-way ANOVA, followed by Original FDR method of Benjamini and Hochberg post-hoc multi-comparison test was used. Data are presented as the median ±interquartile range. Symbols **\*** ^$^ indicate significant (p<0.05) differences between genotypes* (WT vs TG^AC8^), between ages^$^ (young vs old).

**Extended Data Figure 5 - Figure S5.**
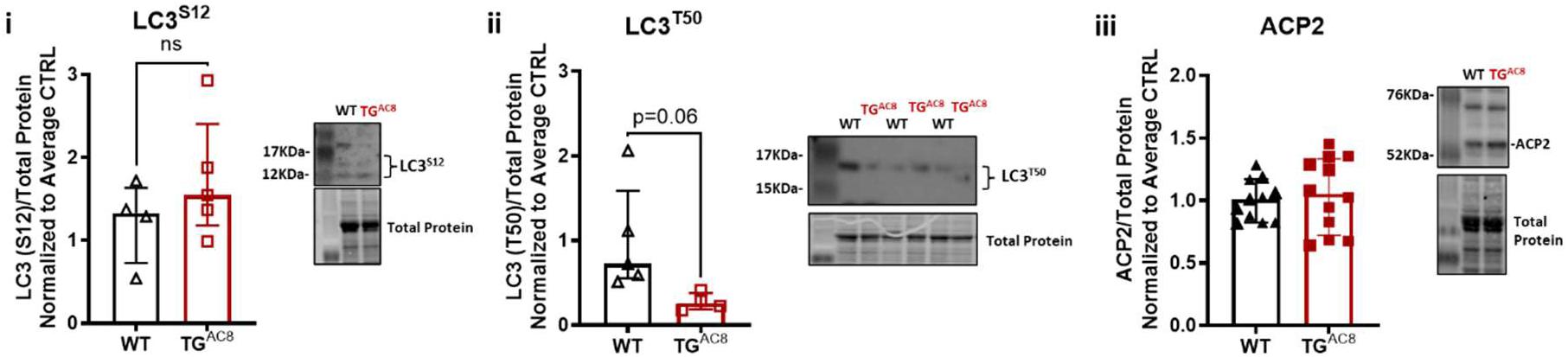
LC3^S12^, LC3^T50^ and ACP2 assessment. Bar graphs and Western blot analysis of autophagy markers in old (19 months) TG^AC8^ and WT littermates (n=5-15 mice per group) **i**), LC3^S12^; **ii**), LC3^T50^; **iii**) ACP2. Unpaired 2-tailed Student t test with Welch’s correction was used. Data are presented as the median ±interquartile range.

**Extended Data Figure 5 - Figure S6.**
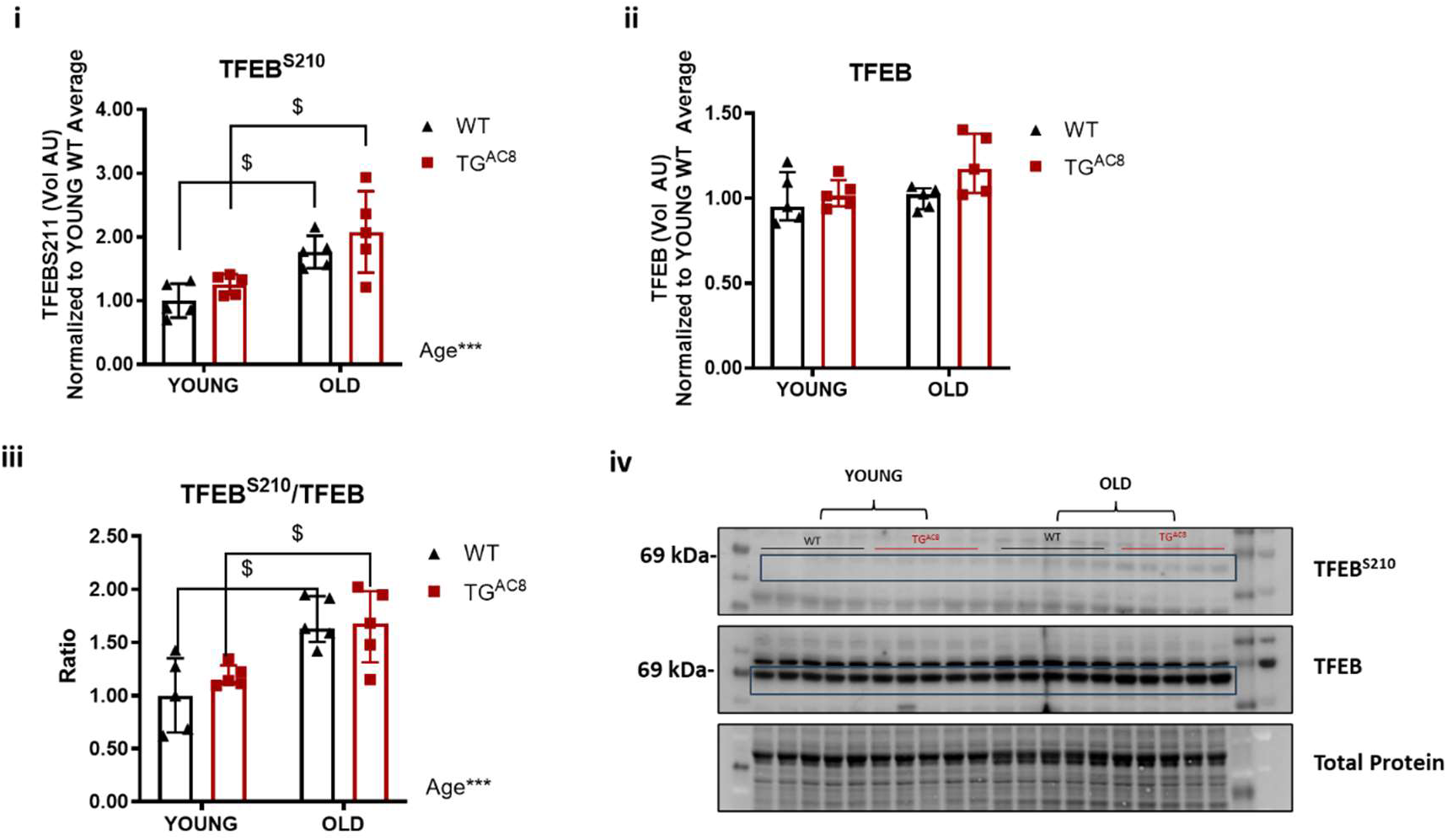
TFEB^S210^ and its ratio to total TFEB increase with aging both in TG^AC8^ and age-matched WT littermates. Bar graphs and Western blot analysis of TFEB and its phosphoform TFEB^S210^ in TG^AC8^ and WT at 3-4 months and 21 months of age (n=5 mice per group). **i**), TFEB^S210^; **ii**), total TFEB; **iii**), TFEB^S210^/TFEB ratio; **iv**), WB representative image. A two-way ANOVA, followed by Original FDR method of Benjamini and Hochberg post-hoc multi-comparison test was used. Data are presented as the median ±interquartile range. Symbols ^$^ indicate significant (p<0.05) differences between ages (young vs old).

## SUPPLEMENTAL DATA FIGURE LEGENDS

**Table 1.**
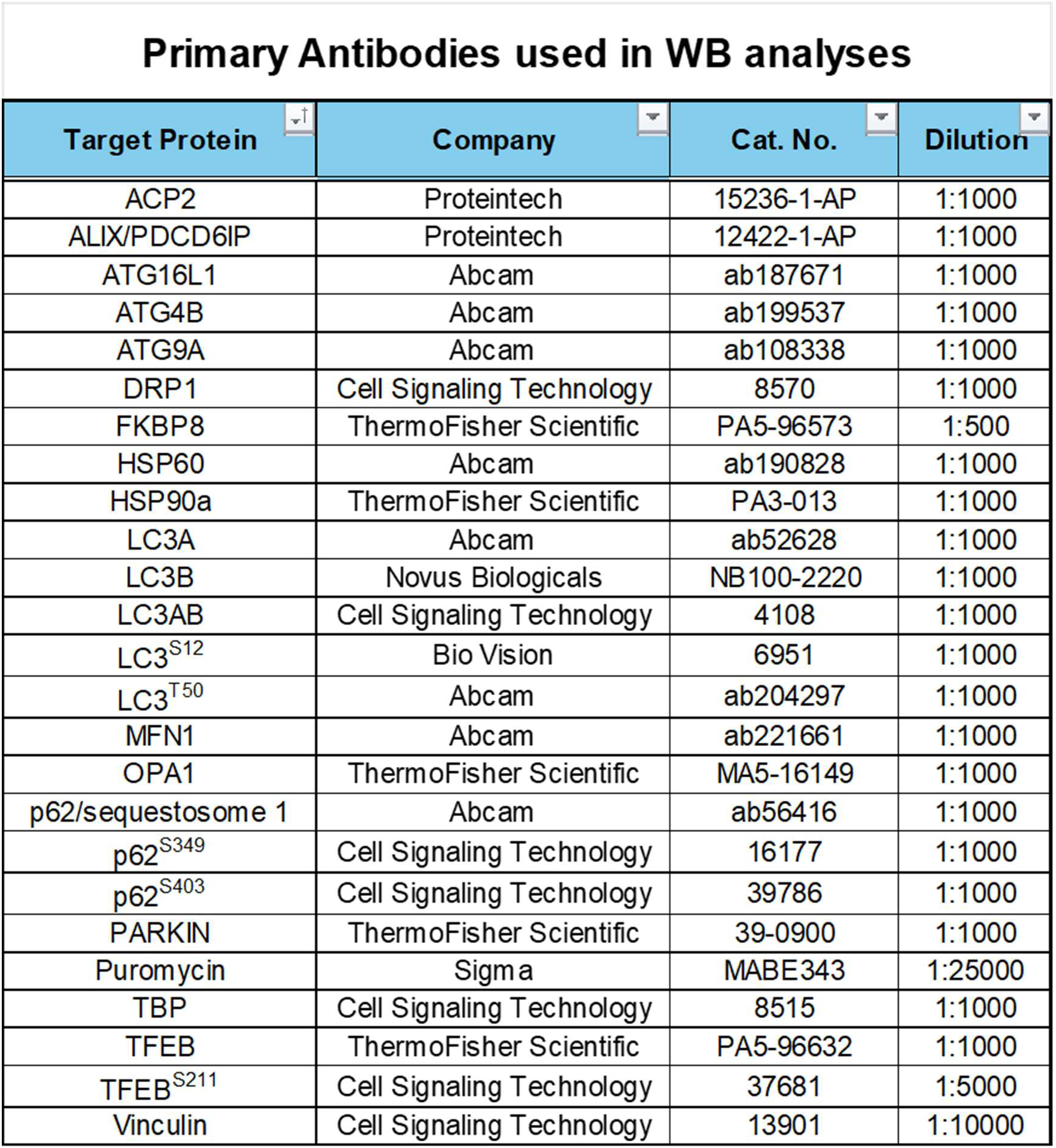
Primary antibodies used in WB.

**SUPPLEMENT DATA – TABLE 1**
| Primary Antibodies used in WB analyses |  |  |  |
| --- | --- | --- | --- |
| Target Protein | Company | Cat. No. | Dilution |
| ACP2 | Proteintech | 15236-1-AP | 1:1000 |
| ALIX/PDCD6IP | Proteintech | 12422-1-AP | 1:1000 |
| ATG16L1 | Abcam | ab187671 | 1:1000 |
| ATG4B | Abcam | ab199537 | 1:1000 |
| ATG9A | Abcam | ab108338 | 1:1000 |
| DRP1 | Cell Signaling Technology | 8570 | 1:1000 |
| FKBP8 | ThermoFisher Scientific | PA5-96573 | 1:500 |
| HSP60 | Abcam | ab190828 | 1:1000 |
| HSP90a | ThermoFisher Scientific | PA3-013 | 1:1000 |
| LC3A | Abcam | ab52628 | 1:1000 |
| LC3B | Novus Biologicals | NB100-2220 | 1:1000 |
| LC3AB | Cell Signaling Technology | 4108 | 1:1000 |
| LC3<sup>S12</sup> | Bio Vision | 6951 | 1:1000 |
| LC3<sup>T50</sup> | Abcam | ab204297 | 1:1000 |
| MFN1 | Abcam | ab221661 | 1:1000 |
| OPA1 | ThermoFisher Scientific | MA5-16149 | 1:1000 |
| p62/sequestosome 1 | Abcam | ab56416 | 1:1000 |
| p62<sup>S349</sup> | Cell Signaling Technology | 16177 | 1:1000 |
| p62<sup>S403</sup> | Cell Signaling Technology | 39786 | 1:1000 |
| PARKIN | ThermoFisher Scientific | 39-0900 | 1:1000 |
| Puromycin | Sigma | MABE343 | 1:25000 |
| TBP | Cell Signaling Technology | 8515 | 1:1000 |
| TFEB | ThermoFisher Scientific | PA5-96632 | 1:1000 |
| TFEB<sup>S211</sup> | Cell Signaling Technology | 37681 | 1:5000 |
| Vinculin | Cell Signaling Technology | 13901 | 1:10000 |

**Table 2.**
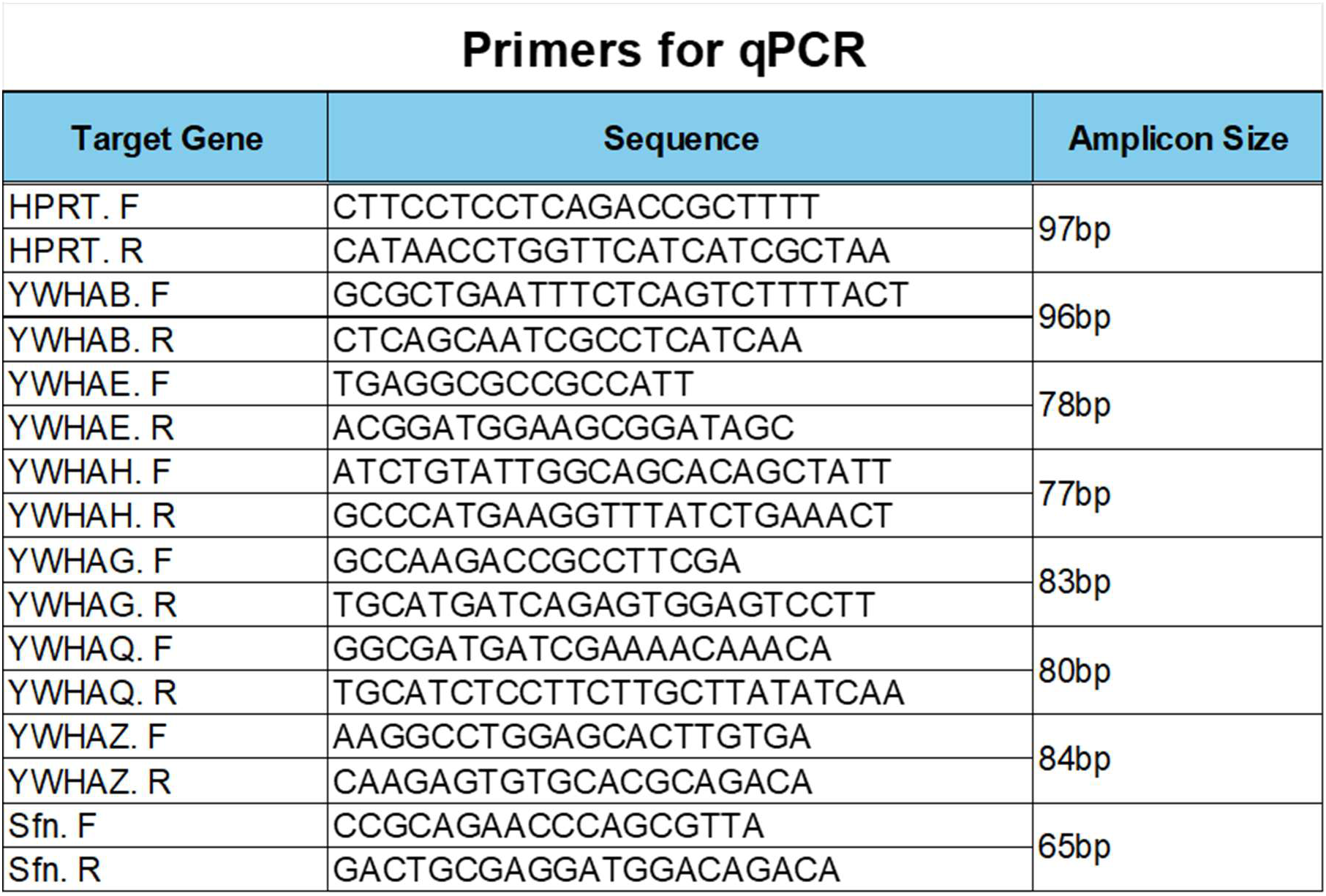
List of primers used in qPCR.

**Table 3.** List of players (proteOMICs) in the <u>Autophagy Pathway</u> in TG^AC8^ and WT littermates at 3-4 months of age.

**SUPPLEMENT DATA – TABLE 3****Autophagy Pathway (ProteOMICs) in YOUNG TG<sup>AC8</sup> vs WT**
| Symbol | Entrez Gene Name | Entrez G | Expr Ot | Expr p-value |
| --- | --- | --- | --- | --- |
| ATG3 | autophagy related 3 | Atg3 | 11.349 | 0.0153 |
| ATG9A | autophagy related 9A | Atg9a | 30.993 | 0.016 |
| ATG13 | autophagy related 13 | Atg13 | 21.004 | 0.0255 |
| BECN1 | beclin 1 | Becn1 | 14.637 | 0.000037 |
| CTSA | cathepsin A | Ctsa | 16.215 | 0.00451 |
| CTSB | cathepsin B | Ctsb | 8.728 | 0.0254 |
| CTSC | cathepsin C | Ctsc | 12.328 | 0.0282 |
| CTSD | cathepsin D | Ctsd | 12.091 | 0.0335 |
| CTSF | cathepsin F | Ctsf | 9.745 | 0.038 |
| CTSS | cathepsin S | Ctss | 24.59 | 0.00705 |
| CTSV | cathepsin V | Ctsl | 5.85 | 0.0372 |
| CTSZ | cathepsin Z | Ctsz | 32.255 | 0.00856 |
| LAMP1 | lysosomal associated membrane protein 1 | Lamp1 | 9.427 | 0.0149 |
| LAMP2 | lysosomal associated membrane protein 2 | Lamp2 | 17.389 | 0.00142 |
| MAP1LC3A | microtubule associated protein 1 light chain 3 alpha | Map1lc3a | -45.27 | 0.0423 |
| PIK3C3 | phosphatidylinositol 3-kinase catalytic subunit type 3 | Pik3c3 | 16.209 | 0.00709 |
| PIK3R4 | phosphoinositide-3-kinase regulatory subunit 4 | Pik3r4 | 12.184 | 0.00452 |
| RB1CC1 | RB1 inducible coiled-coil 1 | Rb1cc1 | 8.639 | 0.047 |
| SQSTM1 | sequestosome 1 | Sqstm1 | 22.553 | 0.00374 |
| STX17 | syntaxin 17 | Stx17 | 21.775 | 0.0379 |
| ULK1 | unc-51 like autophagy activating kinase 1 | Ulk1 | 14.444 | 0.04 |

**Table 4.** List of players (proteOMICs) in the <u>Phagosome Formation Pathway</u> in TG^AC8^ and WT littermates at 3-4 months of age.

**SUPPLEMENT DATA – TABLE 4****Phagosome Formation Pathway (ProteOMICs) in YOUNG WT/TG<sup>AC8</sup>**
| Symbol | Entrez Gene Name | Entrez Gene | Expr Ot | Expr p- |
| --- | --- | --- | --- | --- |
| FCER1G | Fc epsilon receptor lg | Fcer1g | 48.805 | 0.00259 |
| FN1 | fibronectin 1 | Fn1 | 19.588 | 0.00179 |
| FNBP1 | formin binding protein 1 | Fnbp1 | 13.954 | 0.000824 |
| ITGA2 | integrin subunit alpha 2 | Itga2 | 33.08 | 0.0254 |
| ITGA5 | integrin subunit alpha 5 | Itga5 | 15.639 | 0.0272 |
| ITGB2 | integrin subunit beta 2 | Itgb2 | 26.988 | 0.00412 |
| MBL2 | mannose binding lectin 2 | Mbl2 | 21.695 | 0.0033 |
| MRC1 | mannose receptor C-type 1 | Mrc1 | 26.984 | 0.0147 |
| MRC2 | mannose receptor C type 2 | Mrc2 | 14.806 | 0.0297 |
| PDIA3 | protein disulfide isomerase family A member 3 | Pdia3 | 13.635 | 0.00757 |
| PIK3C3 | phosphatidylinositol 3-kinase catalytic subunit type 3 | Pik3c3 | 16.209 | 0.00709 |
| PIK3CB | phosphatidylinositol-4,5-bisphosphate 3-kinase catalytic subunit bet | Pik3cb | 13.834 | 0.0183 |
| PIK3R1 | phosphoinositide-3-kinase regulatory subunit 1 | Pik3r1 | 8.8 | 0.0168 |
| PIK3R4 | phosphoinositide-3-kinase regulatory subunit 4 | Pik3r4 | 12.184 | 0.00452 |
| PLCG2 | phospholipase C gamma 2 | Plcg2 | 14.581 | 0.00011 |
| PRKCI | protein kinase C iota | Prkci | 23.901 | 0.0307 |
| RAC2 | Rac family small GTPase 2 | Rac2 | 34.574 | 0.0489 |
| RHOB | ras homolog family member B | Rhob | 9.77 | 0.0167 |
| RHOG | ras homolog family member G | Rhog | 14.423 | 0.0036 |
| RHOT1 | ras homolog family member T1 | Rhot1 | 8.752 | 0.0124 |
| RHOT2 | ras homolog family member T2 | Rhot2 | 5.576 | 0.018 |
| SYK | spleen associated tyrosine kinase | Syk | 16.644 | 0.0139 |
| VTN | vitronectin | Vtn | 33.51 | 0.00764 |

**Table 5.** List of players (proteOMICs) in the <u>Phagosome Maturation Pathway</u> in TG^AC8^ and WT littermates at 3-4 months of age.

| Symbol | Entrez Gene Name | Entrez Gene/ | Expr Ot | Expr p-vi |
| --- | --- | --- | --- | --- |
| ATP6V0A1 | ATPase H+ transporting V0 subunit a1 | Atp6v0a1 | 11.991 | 0.0486 |
| ATP6V0D1 | ATPase H+ transporting V0 subunit d1 | Atp6v0d1 | 16.118 | 0.00755 |
| ATP6V1A | ATPase H+ transporting V1 subunit A | Atp6v1a | 10.447 | 0.0188 |
| ATP6V1B2 | ATPase H+ transporting V1 subunit B2 | Atp6v1b2 | 7.786 | 0.00817 |
| ATP6V1F | ATPase H+ transporting V1 subunit F | Atp6v1f | 8.739 | 0.047 |
| ATP6V1H | ATPase H+ transporting V1 subunit H | Atp6v1h | 12.95 | 0.00459 |
| B2M | beta-2-microglobulin | B2m | 16.883 | 0.0403 |
| CANX | calnexin | Canx | 10.298 | 0.00522 |
| CTSA | cathepsin A | Ctsa | 16.215 | 0.00451 |
| CTSB | cathepsin B | Ctsb | 8.728 | 0.0254 |
| CTSC | cathepsin C | Ctsc | 12.328 | 0.0282 |
| CTSD | cathepsin D | Ctsd | 12.091 | 0.0335 |
| CTSF | cathepsin F | Ctsf | 9.745 | 0.038 |
| CTSS | cathepsin S | Ctss | 24.59 | 0.00705 |
| CTSV |  | Ctsl | 5.85 | 0.0372 |
| CTSZ | cathepsin Z | Ctsz | 32.255 | 0.00856 |
| DCTN4 | dynactin subunit 4 | Dctn4 | 9.453 | 0.00641 |
| DYNC1LI2 | dynein cytoplasmic 1 light intermediate chain 2 | Dync1li2 | 10.201 | 0.0127 |
| GPAA1 | glycosylphosphatidylinositol anchor attachment 1 | Gpaa1 | 14.343 | 0.00188 |
| HLA-A | major histocompatibility complex, class I, A | H2-D1 | 8.857 | 0.039 |
| LAMP1 | lysosomal associated membrane protein 1 | Lamp1 | 9.427 | 0.0149 |
| LAMP2 | lysosomal associated membrane protein 2 | Lamp2 | 17.389 | 0.00142 |
| NAPA | NSF attachment protein alpha | Napa | 8.606 | 0.00126 |
| NAPG | NSF attachment protein gamma | Napg | 7.666 | 0.0255 |
| NCF1 | neutrophil cytosolic factor 1 | Ncf1 | 55.062 | 0.0231 |
| NSF | N-ethylmaleimide sensitive factor, vesicle fusing ATPase | Nsf | 5.425 | 0.0259 |
| PIK3C3 | phosphatidylinositol 3-kinase catalytic subunit type 3 | Pik3c3 | 16.209 | 0.00709 |
| PRDX1 |  | Prdx1 | 16.381 | 9.22E-07 |
| PRDX2 | peroxiredoxin 2 | Prdx2 | 6.909 | 0.00636 |
| PRDX5 | peroxiredoxin 5 | Prdx5 | -3.968 | 0.0034 |
| RAB5B | RAB5B, member RAS oncogene family | Rab5b | 5.72 | 0.0287 |
| STX16 | syntaxin 16 | Stx16 | 9.904 | 0.0404 |
| TSG101 | tumor susceptibility 101 | Tsg101 | 6.717 | 0.00715 |
| TUBA8 | tubulin alpha 8 | Tuba8 | 5.722 | 0.0253 |
| TUBB | tubulin beta class I | Tubb5 | 14.606 | 0.014 |
| TUBB4B | tubulin beta 4B class IVb | Tubb4b | 9.131 | 0.00805 |
| TUBB6 | tubulin beta 6 class V | Tubb6 | 21.975 | 0.00626 |
| VPS37A | VPS37A subunit of ESCRT-I | Vps37a | 6.268 | 0.0305 |

---

Pathway 3 for Table 3. <u>Autophagy Pathway</u> (proteOMICs) in TG^AC8^ and WT littermates at 3-4 months of age.

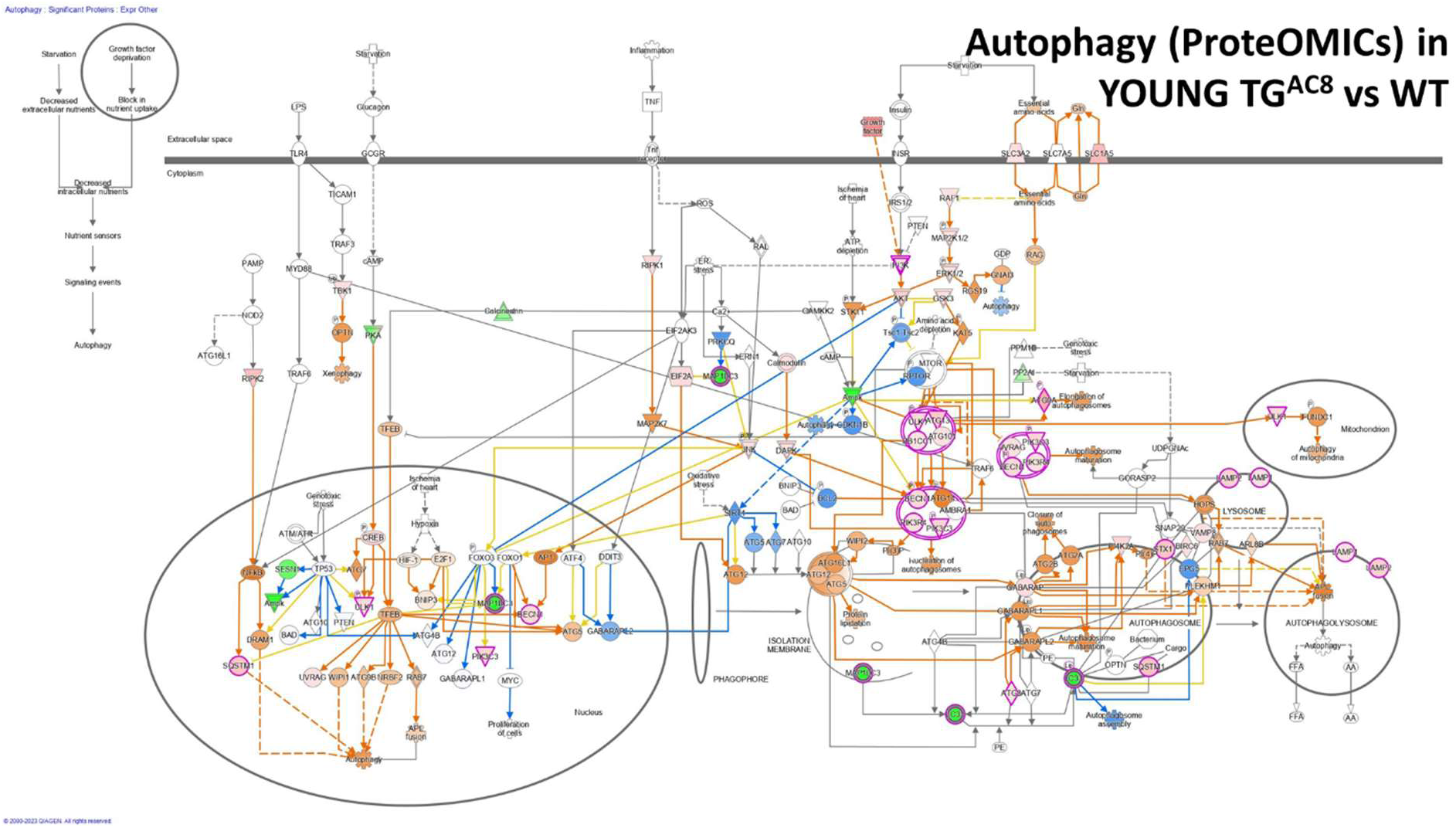

Pathway 4 for Table 4. <u>Phagosome Formation Pathway</u> (proteOMICs) in TG^AC8^ and WT littermates at 3-4 months of age.

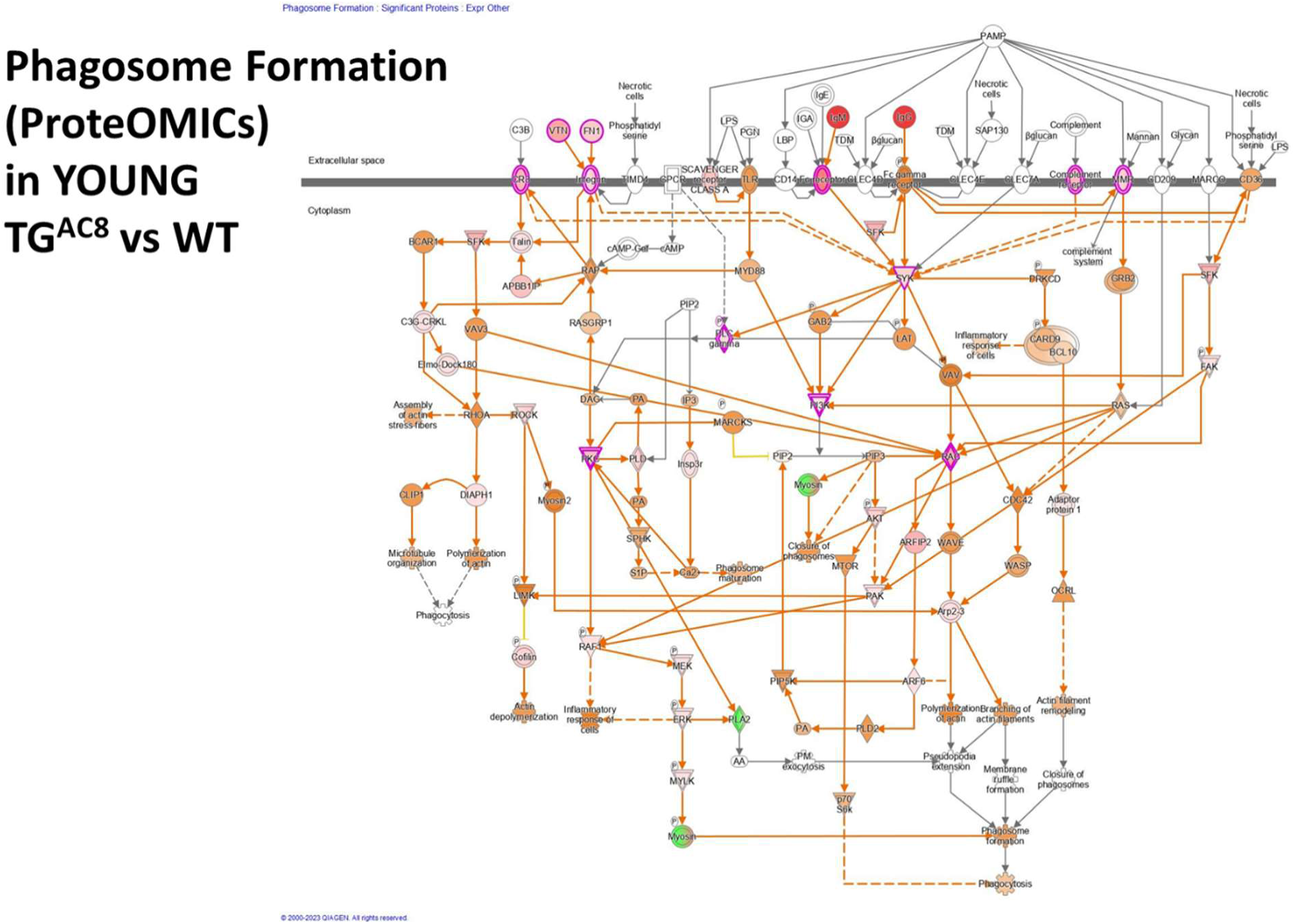

Pathway 5 for Table 5. <u>Phagosome Maturation Pathway</u> (proteOMICs) in TG^AC8^ and WT littermates at 3-4 months of age.

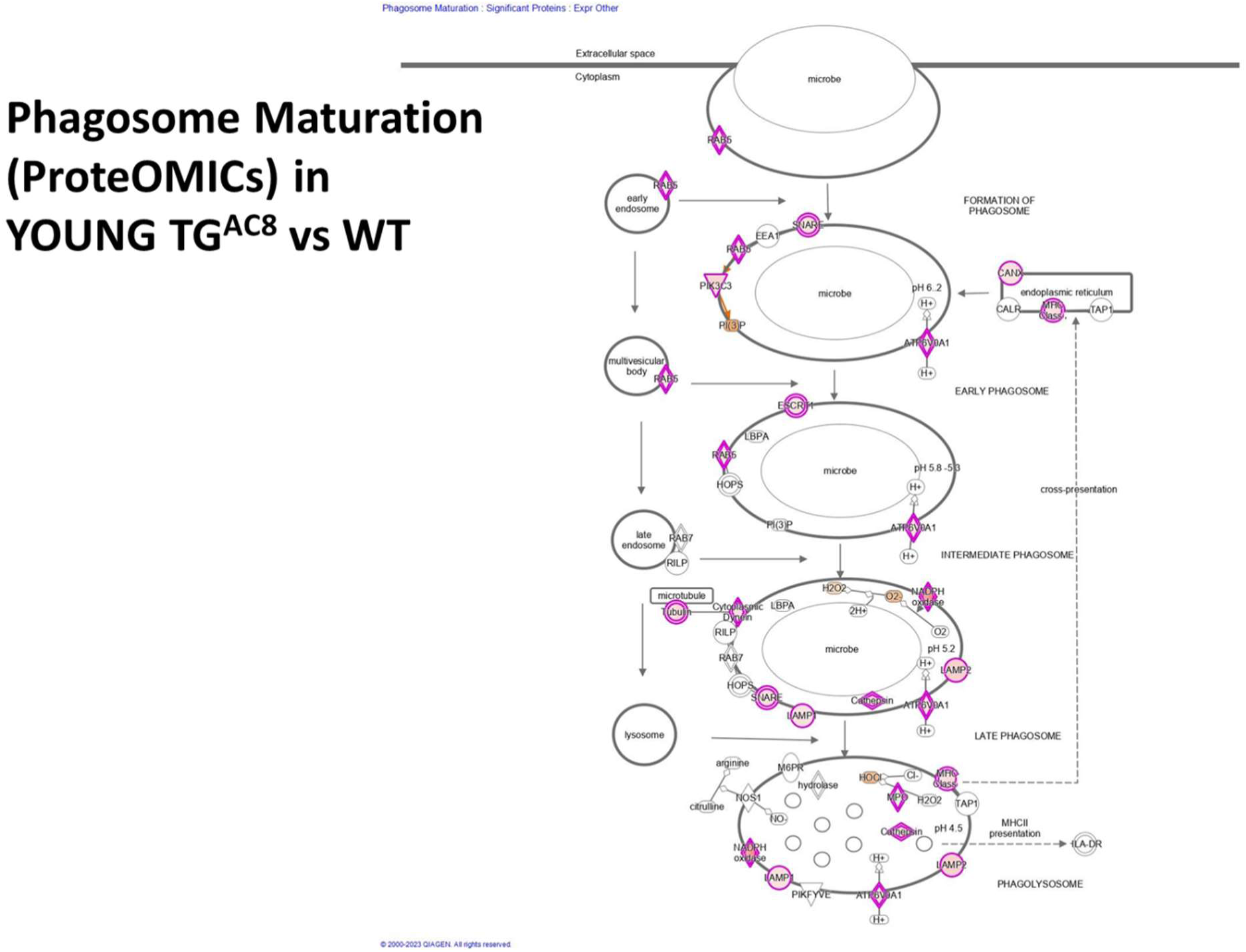

